# Spatially-targeted proteomics of the host-pathogen interface during staphylococcal abscess formation

**DOI:** 10.1101/2020.09.01.267773

**Authors:** Emma R. Guiberson, Andy Weiss, Daniel J. Ryan, Andrew J. Monteith, Kavya Sharman, Danielle B. Gutierrez, William J. Perry, Richard M. Caprioli, Eric P. Skaar, Jeffrey M. Spraggins

## Abstract

*Staphylococcus aureus* is a common cause of invasive and life-threatening infections that are often multi-drug resistant. To develop novel treatment approaches, a detailed understanding of the complex host-pathogen interactions during infection is essential. This is particularly true for the molecular processes that govern the formation of tissue abscesses, as these heterogeneous structures are important contributors to staphylococcal pathogenicity. To fully characterize the developmental process leading to mature abscesses, temporal and spatial analytical approaches are required. Spatially targeted proteomic technologies, such as micro liquid extraction surface analysis, offer insight into complex biological systems including detection of bacterial proteins, and their abundance in the host environment. By analyzing the proteomic constituents of different abscess regions across the course of infection, we defined the immune response and bacterial contribution to abscess development through spatial and temporal proteomic assessment. The information gathered was mapped to biochemical pathways to characterize the metabolic processes and immune strategies employed by the host. These data provide insights into the physiological state of bacteria within abscesses and elucidate pathogenic processes at the host-pathogen interface.

## Introduction

*Staphylococcus aureus* is one of the leading causes of bloodstream infections worldwide^1^. In the United States alone, this bacterium is responsible for more than 600,000 hospitalizations annually^2^ and patients with *S. aureus* bacteremia have a 30-day mortality rate of 20%^3^. *S. aureus* bacteremia often spreads to other body sites leading to the formation of abscesses, most commonly affecting the liver, kidneys, brain, and heart tissues^4^.

The formation of organ abscesses is a critical strategy to ensure *S. aureus* survival within the host^5^. Abscesses offer a temporary refuge for *S. aureus*, allowing the enclosed bacteria to resist the actions of the immune system, thereby securing persistence within host tissues^6,7^. In a murine model of systemic infection, the formation of soft tissue abscesses follows distinct phases (stages I-IV)^5^, driven by the recruitment of immune cells (e.g. neutrophils) ~2 days post infection (dpi)^7^, and the development of a bacterial nidus in the center of the abscess ~4-5 dpi (staphylococcal abscess colony, SAC)^7^. The developing SAC is surrounded by necrotic tissue, a fibrin pseudocapsule^7,8^, and an outer microcolony-associated meshwork^6^. At the end of abscess development (~15-30 dpi), the persistent and increasingly larger lesions rupture and release bacteria that seed new abscesses or cause secondary infections^7^.

Abscess formation requires the involvement of both host and bacterial factors. Several staphylococcal proteins, including both coagulases, Coa and vWbp, are essential for fibrin pseudocapsule formation and abscess development^9^. Additionally, a few other *S. aureus* proteins, associated with virulence, (e.g. Emp, Eap, Hla, IsdA, IsdB, SdrC, SrtA, Spa) have been found to be required for abscess formation^6–8,10,11^. The majority of these data describing bacterial contributions to abscess development were generated from histological stains or characterization of isogenic *S. aureus* mutants and their ability to persist within host tissues. However, unbiased studies that aim to assess the bacterial abscess proteome are sparse. This is similarly true for proteinaceous host factors in proximity to the abscess. While it is established that specific cell types (e.g. fibroblasts and neutrophils^12^) or immune proteins (e.g. the metal scavenging protein calprotectin^13^) play essential roles during the immune response to *S. aureus*, we lack a detailed understanding of specific cellular processes involved in the host response to tissue colonization by *S. aureus*. Although a previous study from our laboratory addressed some of these shortcomings by assessing the proteinaceous composition of kidneys infected with *S. aureus*^14^, the current study also examines the temporal and spatial aspects of abscess development.

Various proteomic technologies offer insight into complex biological systems, including the interplay between host and pathogen during infection^15,16^. Recent technological advances have enabled pairing of spatially targeted surface sampling with high-performance mass spectrometry for molecular analysis of tissue. Imaging mass spectrometry (IMS) technology, such as matrix-assisted laser desorption/ionization (MALDI) IMS, offers the unique combination of high molecular specificity and high spatial resolution imaging^17–19^. However, the sensitivity of MALDI IMS for analysis of large proteins (> 30 kDa) can be limited due to low ionization efficiency of proteins from tissue and the poor transmission efficiency of high mass-to-charge ratio (m/z) ions^19,20^. The identification of proteins with MALDI IMS is also challenging due to inefficient fragmentation of low charge state species. To facilitate this process, we have investigated the utility of complementary surface sampling technologies for discovery-based proteomic studies.

Traditional liquid chromatography coupled with tandem mass spectrometry (LC-MS/MS) of peptides derived from proteolytic digestion provides the greatest proteomic sensitivity ^21^. LC-MS/MS requires liquid samples, usually through homogenization of tissue, limiting spatial information from the sample of interest. To address this, we recently introduced a spatially targeted, bottom-up proteomics workflow used in analysis of *S. aureus* kidney infection^22^. Specific foci were targeted using picoliter-sized droplets of trypsin protease and the resulting proteolytic peptides were sampled using liquid extraction surface analysis (LESA). The entire process, termed microLESA, is histologically guided using autofluorescence microscopy. Herein, we expand on this work using microLESA to analyze regions from the abscess community (SAC), host-pathogen interface surrounding the community, and regions of cortical tissue of *S. aureus* infected kidneys. Spatial and temporal proteomic changes were examined by sampling various time points allowing us to follow responses of both the pathogen and the immune system over the course of infection. By studying three defined regions, and their dynamics over the course of the infection, changes in both host and pathogen can be observed across the organ.

## Results and Discussion

### Identification of bacterial and host proteins in infected tissue

To explore the role of host and bacterial proteins throughout the development of staphylococcal organ abscesses, we analyzed proteins in the abscesses and surrounding areas over time. We focused on kidneys as one of the most commonly utilized model systems for staphylococcal abscess formation. Samples were extracted from one of three defined regions (SAC, interface, and cortex; shown in Fig. 1A) at 4 and 10 dpi (Fig. 1B). These extractions were guided by fluorescence microscopy images (transgenic fluorophore and autofluorescence), which highlights the fluorescent bacterial communities and overall tissue morphology that allowed for differentiation between SAC and adjacent regions. The two time points were selected to ensure sufficient abscess size for extraction, as at 4 dpi abscesses are consistently large enough for micoLESA extraction and at 10 dpi the infection has greatly progressed. We hypothesized that the unique spatial and temporal proteomic analysis would discern i) *S. aureus* physiology and production of virulence determinants within the abscess microenvironment (SAC), ii) onset of the immune response including infection-mediated influx and action of immune cells (interface), as well as iii) organ-wide responses to infection (cortex). A total of 2,399 proteins were identified across all three regions of interest (ROI) and two time points (Table S1 and S2), with an average of 1,153 proteins per time point and ROI (Fig. 1B, Table S1 and S2). Of the proteins detected, 31 were of bacterial origin (Table S2). We identified an additional 26 bacterial proteins that were present in only one set of serial sections or one biological replicate and are not included in Figure 1B due to lower confidence (Table S3). This variability in detection could result from low protein abundance or heterogeneity amongst abscesses, a known challenge when studying abscess formation^23,24^. Nevertheless, we conclude the identification of these proteins is a strength of the method, as bacterial proteins from *S. aureus* infections are difficult to detect and measure due to ion suppression effects from abundant host proteins within tissues. Since we spatially targeted the abscess region, the inherent ‘chemical noise’ from the highly abundant host proteins was reduced, greatly improving the sensitivity for the bacterial proteins. Improving the coverage of the bacterial proteome detectable within the tissue microenvironment provided a more complete description of how *S. aureus* molecular machinery contributes to abscess formation and progression.

**Figure 1:**
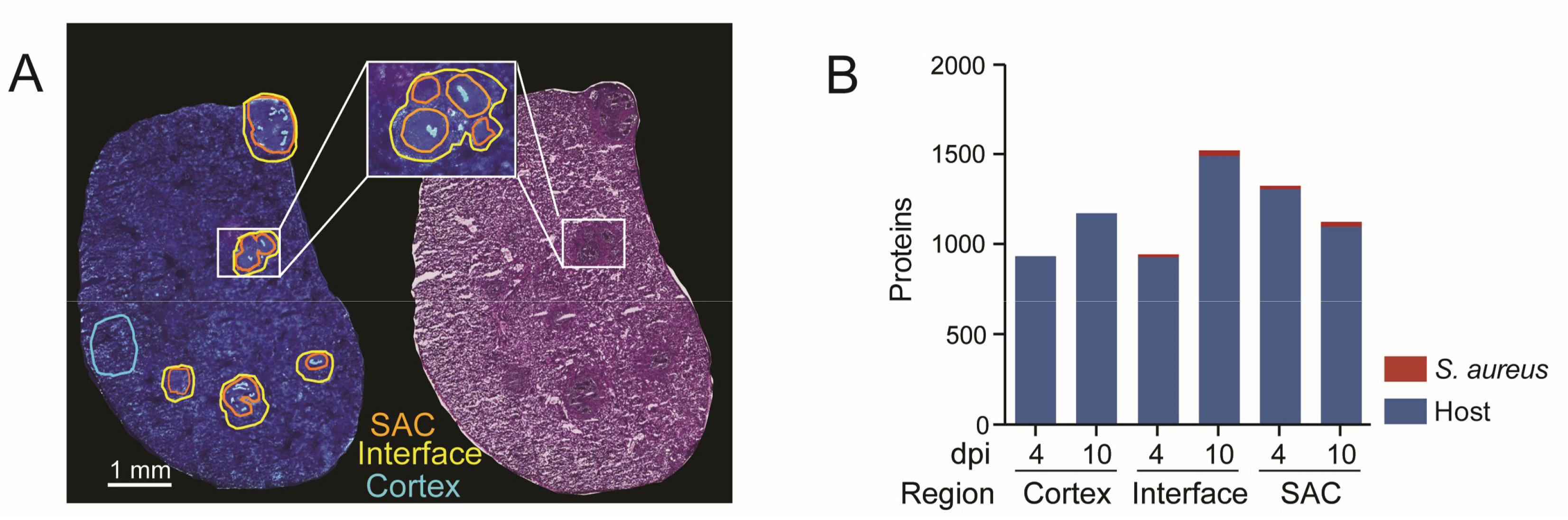
microLESA sampling strategy and results. A) Schematic of regions selected for collection shown using autofluorescence and H&E. B) Overview of bacterial and host proteins identified at different timepoints and across different regions within infected renal tissue. Total host proteins: 2399. Total bacterial proteins: 31.

### Bacterial factors

#### Detection of high abundance proteins related to translation

The microLESA workflow detected 31 bacterial proteins in the defined SAC and interface regions, several of which are involved in the translation process (Fig. 2). These include ribosomal proteins (50S ribosomal proteins L7/L12 [RplL] and L17 [RplQ]), a tRNA-ligase (ThrS) as well as elongation factors Tu (Tuf)^25^ and Ts (Tsf)^26^. Several of these intracellular highly abundant proteins were detected in the SAC region and also in the abscess interface (Fig. 2). While we cannot exclude that a small fraction of the bacterial population resides outside of the observable SAC, or the potential for these proteins to ‘leak’ from the SAC into adjacent regions during the sample preparation process, cell death or autolysin (Atl)-mediated lysis of a staphylococcal subpopulation may cause the release of bacterial factors into the abscess microenvironment^27^. It was previously shown that decreased autolysis by *S. aureus* results in a moderate decrease in renal bacterial numbers in a murine model of systemic infection at 7 dpi^28^, supporting the notion that Atl plays a role during for spread to or colonization of this organ. While our data does not allow us to finally conclude why we detect cellular bacterial proteins outside of the SAC, it is intriguing to speculate that (auto)lytic processes are important during abscess formation, potentially aiding infection through the release of intracellular factors.

**Figure 2:**
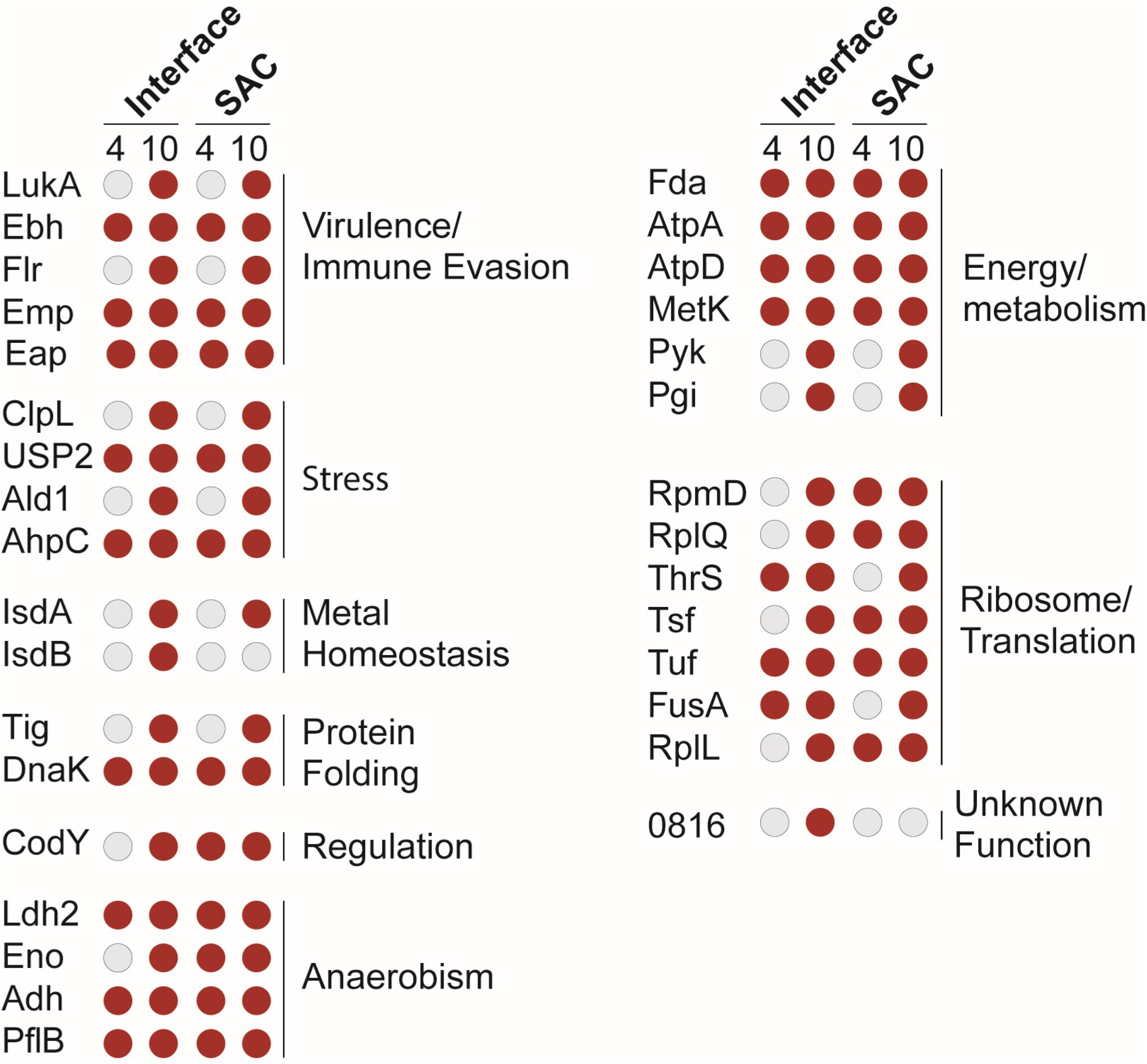
Overview of *S. aureus* proteins identified in the different abscess regions. Staphylococcal proteins were found in the interface and SAC of renal abscesses at 4 or 10 dpi. Red circles denote proteins that were present at the specific timepoint/region, while grey circles depict the absence of a specific protein. General functional categories for proteins and protein groups are shown. Further information concerning proteins in this list can be found in Table S2. 0816: SAUSA300_0816; 1656: SAUSA300_1656.

#### S. aureus *heme metabolism proteins in the abscess*

In response to the presence of *S. aureus* in tissue, the host immune system initiates a variety of anti-bacterial strategies. Nutritional immunity is one such approach in which the immune system limits bacterial access to essential micronutrients (e.g. transition metals) to hinder bacterial growth and stall infection progression^29^. Host-imposed Fe-starvation within the abscess was recently shown by our group using *in vivo* imaging and IMS strategies^23,24^. Bacterial pathogens have evolved a number of mechanisms to overcome this Fe limitation^29^. To ensure sufficient levels of the Fe-containing co-factor heme, *S. aureus* imports host heme through the action of the Isd heme uptake system^30^. Additionally, heme can be synthesized *de novo* via a coproporphyrin-dependent bacterial heme biosynthesis pathway^31,32^. In our analysis we observed components of the Isd system (IsdA and IsdB: Fig. 2, Table S2) as well as ChdC (identified with lower confidence, Table S3), a member of the heme biosynthetic pathway, to be present within the abscess^33^. Identification of proteins involved in heme uptake as well as *de novo* heme synthesis suggests that i) *S. aureus* within renal abscesses is indeed heme starved and ii) that multiple strategies are employed to overcome this limitation. These proteins were only detected at 10 dpi (Fig. 2), indicating that metal starvation and corresponding bacterial responses are likely more abundant at later time points during infection.

#### Bacterial factors related to protein stress

Other host-imposed stresses, including production of reactive oxygen species (ROS), reactive nitrogen species (RNS), and elevated temperatures during fever, can damage the bacterial protein pool^34–37^. To protect the staphylococcal proteins from host-derived stresses, bacterial inducible heat shock proteins (Hsps)^38^ assist with protein folding and proteostasis^39^. Although this study cannot distinguish between basal and stress-induced expression of such factors, the detection of the major bacterial Hsp DnaK at both the 4 dpi and 10 dpi time points (Fig. 2, Table S2) is in accordance with the established roles of Hsps during stress and contact with the host. DnaK is not only involved in maintenance of cellular protein pools, but also plays a role in the folding of *de novo* synthesized proteins, e.g. when adapting to changing environmental conditions^40^. This function is performed in concert with other chaperones, such as the ribosome-associated intracellular peptidyl prolyl cis/trans isomerase (PPIase)^41^, referred to as Trigger factor (TF, Tig)^42^, detected in the SAC at 10 dpi (Fig. 2, Table S2). The cooperative nature of these two proteins during stress conditions is demonstrated by both *dnak* and *tig* mutants being viable under laboratory conditions, but a DnaK/TF double mutant being synthetic lethality at temperatures above 30°C. This lethality is likely due to increased protein misfolding under these conditions^42^. In line with the cooperative nature of these proteins, we also found TF to be associated with the SAC at 10 dpi (Fig. 2, Table S2). To our knowledge, the roles of DnaK and TF in *S. aureus* virulence have not been investigated *in vivo*, and it is intriguing to speculate about the importance of these systems during abscess development. The roles of TF are of particular interest, since another *S. aureus* PPIase, PpiB, was previously shown to impact *S. aureus* virulence through a potential function in secretion of nucleolytic and hemolytic proteins^43^.

#### S. aureus *metabolism is shaped by the host environment*

*S. aureus* can circumvent cellular pathways that have been disrupted by macrophage-derived nitrosative stress^41,44^ or hypoxia at sites of infection^45^. Specifically, fermentative pathways^46^ can be employed during glucose catabolism under hypoxia or if oxidative phosphorylation is impaired due to the damage of terminal oxidases by radical nitric oxide^47^.

Fermentation of pyruvate to lactate by *S. aureus* is facilitated by different lactate dehydrogenases (i.e. Ldh1, Ldh2, Ddh)^46^. We detected Ldh2 in the SAC and interface at 4 and 10 dpi (Fig. 2, Table S2), and Ldh1, a nitric oxide-inducible lactate dehydrogenase, in the abscesses at 4 and 10 dpi, as well as in the interface at 10 dpi (Table S3). It was previously shown that loss of *ldh1* increases staphylococcal sensitivity to nitrosative stress and decreases the ability to form renal abscesses^46^. Furthermore, the additional loss of *ldh2* was found to augment the latter phenotype^46^, highlighting the importance of fermentative metabolism for *S. aureus* pathogenicity. In addition to maintaining the cells ability to generate energy under conditions encountered in the abscess, the creation of lactate as a byproduct of fermentation was recently shown to aid in staphylococcal immune evasion^48^. Briefly, bacterial-derived lactate causes alterations of gene expression in host immune cells, e.g. stimulates the production of the anti-inflammatory cytokine Il-10, allowing for persistence in host tissue. These exciting findings further emphasize how bacterial metabolism shapes to its behavior as a pathogen and ultimately its interaction with the immune system.

Activity of metabolic pathways in *S. aureus* is dependent on environmental conditions and controlled by a large number of transcriptional regulators^49^. Analogously, various regulatory proteins govern pathogenesis of the bacterium^50^. Because the production of the vast array of virulence factors encoded by *S. aureus* is energetically costly, cellular nutritional status and virulence are intimately linked, e.g. through the action of transcriptional regulators that sense and respond to alterations in nutrient availability^51^. A prime example of such a connection is the global transcriptional regulator CodY^52,53^. This bifunctional regulator senses the availability of branched chain amino acids^54,55^ and GTP^56^, and responds by controlling a large number of metabolic and virulence-related traits. We detected CodY, as only one of two *bona fide* bacterial transcriptional regulators in our dataset, at 4 and 10 dpi in the SAC, and at 10 dpi in the interface (Fig. 2, Table S2). While physical presence of the protein itself is not indicative of its regulatory state, the detection of CodY serves as reminder that *S. aureus* pathophysiology is directly affected by the conditions encountered in the abscess microenvironment, and that sensing of these stimuli guides virulence of the bacterium within the abscess.

#### S. aureus *secreted proteins*

In addition to the intracellular factors discussed thus far, we identified several staphylococcal proteins that are actively secreted into the abscess environment. Two adhesins belonging to the group of secretable expanded repertoire adhesive molecules (SERAMs)^57^ were detected: the extracellular matrix binding protein (Emp)^58,59^ and the extracellular adhesion protein (Eap)^60^. Both proteins were among the most consistently observed molecules in the SAC as well as in the interface (Fig. 2, Table S2). Our dataset was manually screened for the presence of Eap as the encoding sequence is found in the USA300 LAC genome, but it is not annotated as a protein. Using this approach, we identified peptides for Eap, indicating production of this protein within host tissue. The presence of Emp and Eap is concurrent with their roles in abscess formation and maintenance^61^. Both proteins were detected in all samples at 4 and 10 dpi, supporting the notion that these factors are abundant and important in both the early and late stages of abscess development^7^.

The action of neutrophils is essential for the immune system to clear staphylococcal infection. Neutrophils are recruited to the site of infection after the recognition of host or pathogen derived factors, including C5a, bacterial formylated peptides, or antimicrobial peptides. Corresponding receptors for these signals are the neutrophil receptor C5aR, the formyl peptide receptor (FPR), and the formyl peptide receptor-like-1 (FPRL1)^62,63^. To decrease the negative impact of neutrophil recruitment, *S. aureus* secretes chemotaxis inhibitory proteins that limit the infiltration of immune cells to the site of infection. These secreted factors bind to neutrophil receptors, therefore blocking the recognition of infection-related signal molecules and ultimately preventing neutrophil recruitment. Staphylococcal anti-chemotactic factors include, amongst others^64^, the chemotaxis inhibitory protein of *S. aureus* (CHIPS) that antagonizes C5aR and FPR^65^, and the FPRL1 inhibitory protein (FLIPr) that blocks the FPRL1 and FPR receptors^66^. Both proteins were identified by our analysis (FLIPr: high confidence, Fig. 2, Table S2; CHIPS: lower confidence, Table S3). Notably, we were only able to detect CHIPS and FLIPr at 10 dpi, suggesting that these factors are expressed or accumulate at advanced stages of abscess development. To our knowledge, this is the first time that production of these chemotaxis inhibitory proteins has been detected by an unbiased *in vivo* screen.

Another staphylococcal strategy to minimize the impact of the recruited immune cells is the secretion of pore-forming toxins^67^. Upon secretion by *S. aureus*, these proteins insert into and disrupt the plasma membrane of host cells (e.g. neutrophils), ultimately leading to immune cell death. We identified both components of the bi-component leukotoxin LukAB^68,69^ (LukA: high confidence, LukB: lower confidence) at 10 dpi (Fig. 2, Tables S2 and S3). Presence of these proteins was recently correlated to abscess formation, further validating our method^68^. LukB, as well as several other proteins identified by our microLESA approach (i.e. Chp, Emp and Eap) are expressed under the control of the major virulence regulatory system SaeRS^70^, whose activity can be modulated in response to host-derived signals (e.g. Zn-bound calprotectin or human neutrophil peptides)^71,72^. These results indicate that the SaeRS system may be active during adaptation to the abscess microenvironment and further highlight the interconnected nature of the host-pathogen interface, where the actions of host and bacteria are inseparably intertwined.

### Host-derived factors

#### Spatial and Temporal Changes in the Host Proteome

The majority of proteins identified in this study were of host origin. The large number of murine proteins detected (2,368) allowed for the probing of the relationships between different tissue regions as well as the determination of the proteomic changes within regions over time. At 4 dpi, the majority of identified proteins (676) are common among all three regions. This suggests that 4 days may not be sufficient for the full immune response to be observed and proteinaceously distinct abscess regions to form. The most unique region at this time point is the SAC (Fig. 3). The SAC region also displayed the least changes over the course of infection, where the vast majority of proteins in the region were present at both time points (964, Figure 3B). This indicates that once an abscess community has been formed, the makeup of this region appears to remain fairly stable. At 4 dpi the interface and SAC show high degrees of similarity with an overlap of 211 proteins. As the infection progresses this overlapping region becomes one of the most prominent groups represented in our dataset (635) after proteins unique to the cortex (657), while the overlap between all three regions is less than at 4 dpi (441 vs. 676). This indicates that as the infection continues, the site of infection (SAC and interface) and the cortex grow increasingly more distinct. As depicted in Figure 3B, only half of the proteins detected in the cortex were found at 4 and 10 dpi, which is in stark contrast to the trend seen in the SAC. These large-scale changes in the cortex suggest organ-wide effects of infection by *S. aureus.* We hypothesize that this increase in unique proteins in the cortex is likely due to the resolution of the immune response, in regions that are not in direct contact with the pathogen. This hypothesis is discussed further in the host response section regarding metabolism.

**Figure 3:**
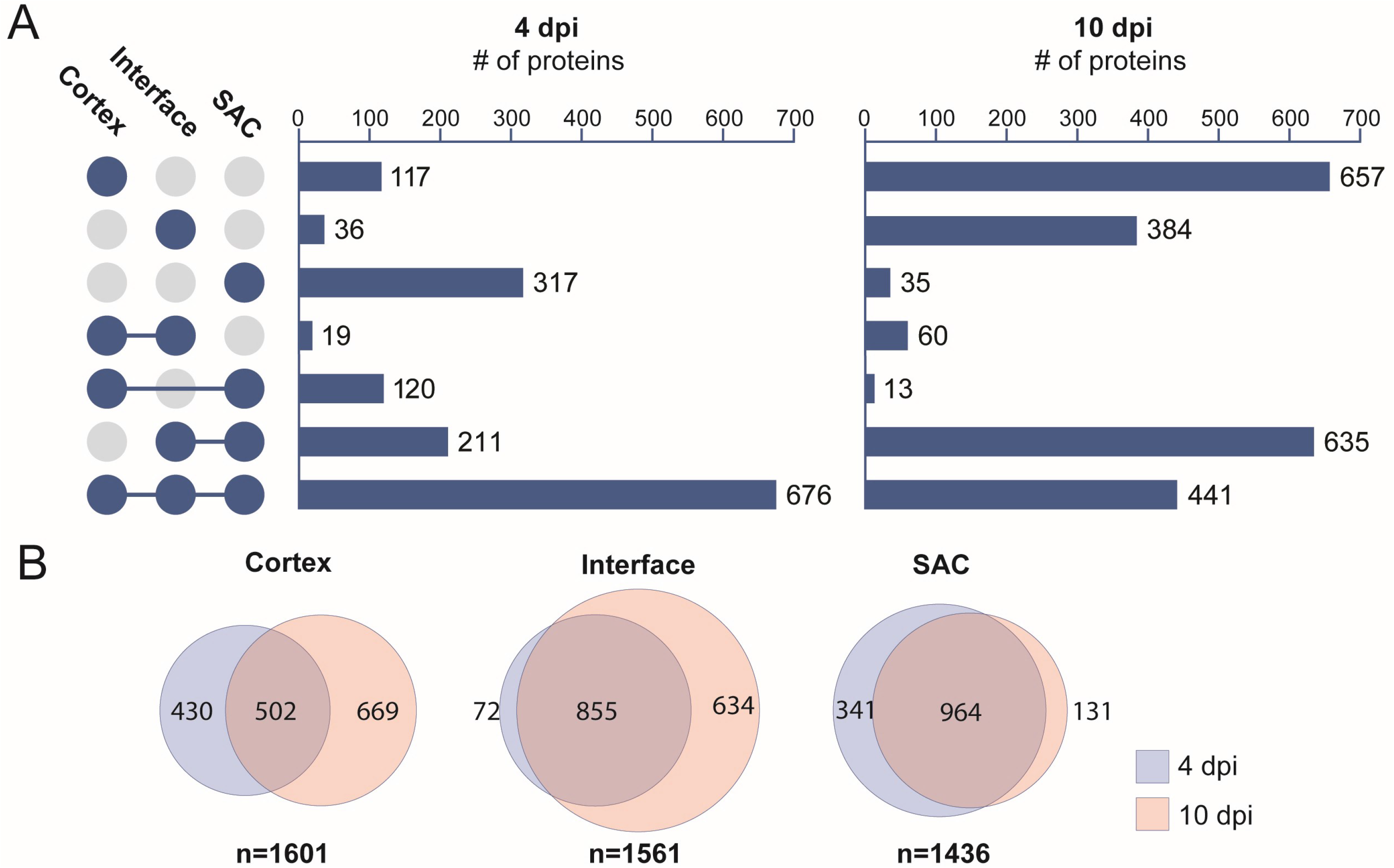
Comparison of the proteome from the different regions in and around staphylococcal tissue abscesses. Data is combined host and bacterial proteins. A) UpSet plot displaying unique and shared proteins identified from the three sampled locations. B) Comparison of the proteome from individual regions over the course of infections (4 dpi vs. 10 dpi).

Amongst the different investigated regions, the most notable trends are in the interface. The number of unique proteins detected only in the interface showed dramatic change over the course of the experiment, increasing greater than ten-fold from 4 dpi (36 proteins) to 10 dpi (384 proteins) as shown in Fig. 3. Additionally, many of the proteins found in the SAC at 4 dpi are shared with the interface by 10 dpi (635 proteins), suggesting that the interface becomes a major site of the competition between host and pathogen. The finding that many proteins were uniquely detected at the interface suggests that we are able to assess the proteome of this region without unwanted contamination from neighboring regions. In the case of unintended cross-contamination between the SAC and the interface, host proteins would likely be common between the two regions leading to a similar makeup, which was not observed in this data set. These findings also support the observation of bacterial cytosolic proteins detected at the interface indeed is biologically relevant and not merely an artifact of sample acquisition. Our results indicate that the interface is a unique and deeply informative tissue environment for understanding staphylococcal pathogenesis.

#### Immune cell distributions

Similar to the rich biology seen from our analysis of bacterial proteins, thousands of identified murine proteins characterize the host response to *S. aureus* infection. A summary of all identified host-derived proteins, including information about localization and time of identification, can be found in Table S1. Consistent with a predicted presence of immune cells in the abscess, the pan-leukocyte marker Receptor-type tyrosine-protein phosphatase C (Ptprc or CD45) is present in the SAC and interface at 4 and 10 dpi (Fig. 4). Curiously, other proteins thought to be specifically expressed by cells of hematopoietic lineages do not follow the same pattern as CD45. Lymphocyte-specific protein 1 (Lsp1) is also found in the cortical regions of the kidney both 4 and 10 dpi, while dedicator of cytokinesis protein 2 (Dock2) is only found in the abscess and interface 10 dpi (Fig. 4). This suggests subtle spatial and temporal differences in the leukocyte populations during infection.

**Figure 4:**
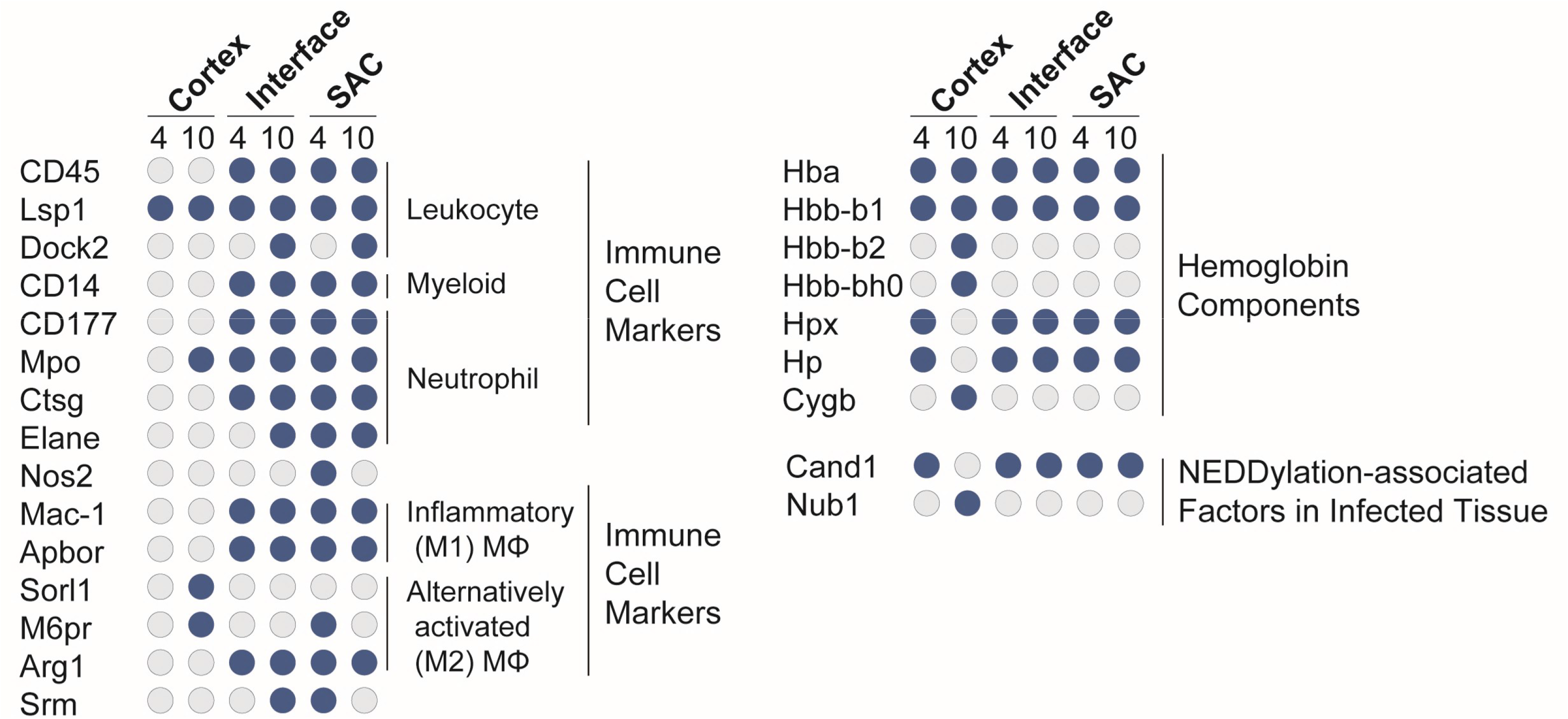
Spatiotemporal distribution of immune cell markers, hemoglobin components, NEDDylation-associated factors in infected tissue. Host immune proteins were found in the interface and SAC of renal abscessess at 4 and 10 dpi. Blue circles denote proteins that were present at the specified timepoint/region, while grey circles depict the absence of a specific protein.

During abscess formation, neutrophils are recruited to the site of infection in high numbers^73–75^. We identified the myeloid cell marker CD14 and neutrophil marker CD177 within the abscess and interface (Fig. 4). Coinciding with the neutrophil surface markers, multiple neutrophil-specific antimicrobial factors are also found in the abscess and interface regions including myeloperoxidase (Mpo), cathepsin G (Ctsg), and neutrophil elastase (Elane) (Fig. 4). In addition, neutrophils use NADPH oxidase to generate high levels of ROS in response to *S. aureus*, and an abundance of NADPH oxidase proteins are also specifically within the abscess and interface. The inducible nitric oxide synthase (Nos2), however, is only found in the abscess 4 dpi (Fig. 4). While it is not possible to fully exclude that Nos2 is present in low concentrations below our limit of detection at this later time point, Nos2 was reliably detected in nearly all samples at the 4 dpi time point. This suggests that even if Nos2 is not absent from the abscess at 10 dpi, the concentration decreases over time.

Relative to neutrophils, macrophages make up a smaller percentage of the immune cells in the abscess^76^. Nevertheless, the presence of integrin alpha-M (Itgam or Mac-1) indicates that activated inflammatory macrophages are found in the abscess and interface (Fig. 4). The expression of apolipoprotein B receptor (Apobr) by macrophages is critical to combating *S. aureus*, as it suppresses activation of the Agr system, a major component of the staphylococcal regulatory landscape^76^ (Fig. 4). Consistent with an integral role in combating *S. aureus*, Apobr is present both within the abscess and interface at 4 and 10 dpi. The presence of sortilin-related receptor (Sorl1), and mannose-6-phosphate receptor (M6pr) at 10 dpi in the cortex suggests the presence of alternatively activated M2-like macrophages involved in tissue repair and remodeling outside of the abscess (Fig. 4). The classical M2-like macrophage marker arginase-1 (Arg1) is not detected in the cortex and is present ubiquitously in the abscess and interface (Fig. 4). Despite Arg1 being linked to M2 polarization, Arg1 is only produced by a quarter of all M2-like macrophages^77^ and is furthermore up-regulated in inflammatory M1-like macrophages^78,79^. Arg1 is necessary for the production of spermine and spermidine, which are uniquely toxic to most methicillin-resistant *S. aureus* (MRSA) strains and integral to killing *S. aureus* during the tissue resolution phase of skin infections^80^. The presence of spermidine synthase (Srm) in the abscess at 4 dpi suggests spermine and spermidine production may play a critical antimicrobial function within abscesses in the kidney^81^ (Fig. 4). These data suggest that during *S. aureus* infection of kidneys, Arg1 may be present in M1 macrophages in the abscess and might not be a reliable marker for the M2 macrophages associated with tissue repair in the cortex 10 dpi.

#### Heme distribution

Heme acts as a critical source of iron for *S. aureus* at the site of infection^82^. While hemoglobin subunit alpha (Hba) and beta-1 (Hbb-b1) are ubiquitously present during infection, hemoglobin subunit beta-2 (Hbb-b2) and beta-H0 (Hbb-b0) are only found in the cortex at 10 dpi (Fig. 4). Importantly, the host can restrict free heme and hemoglobin from *S. aureus* through binding and sequestration by hemopexin (Hpx)^83^ and haptoglobin (Hp)^84^, respectively. Both Hpx and Hp were found in the abscess, interface, and cortex at 4 dpi, but not in the cortex at 10 dpi (Fig. 4). This suggests that while free heme and hemoglobin may be present at the site of infection in the kidney, the simultaneous presence of Hpx and Hp may render heme and hemoglobin biologically unavailable to *S. aureus*. The presence of host factors that limit heme availability to *S. aureus* also serve as an explanation for the previously discussed presence of different staphylococcal proteins aimed to counteract heme limitation during infection (i.e. IsdA, IsdB and ChdC) (Fig. 2). Another heme-containing protein, cytoglobin (Cygb), is exclusively present in the cortex at 10 dpi. Cygb contributes to oxygen diffusion for collagen synthesis during wound healing^85^, regulating nitric oxide levels^86,87^ and detoxifying reactive oxygen species^88^ (Fig. 4). Cygb mirrors the presence of M2 macrophages in the cortex at 10 dpi, that suggests Cygb plays a role in tissue repair and abscess resolution during *S. aureus* infection.

#### Host signaling

The power of microLESA is not just in confirming the presence or absence of proteins, but in allowing for unbiased observations about the regulatory states of the cells. NEDDylation and ubiquitylation are posttranslational modifications that regulate protein degradation by the proteasome, thereby influencing signal transduction^89–94^, inflammasome activation^95,96,97^, autophagy^98,99^, and cell death^100,101^. The cullin-associated NEDD8-dissociated protein 1 (Cand1) regulates NEDD8 activity by sterically inhibiting the assembly of cullin-RING ubiquitin ligases and preventing NEDDylation^102^, while the NEDD8 ultimate buster 1 (Nub1) specifically recruits NEDD8 to the proteasome for degradation^103,104^. In the abscess, interface (both at 4 and 10 dpi), and cortex (at 4 dpi) Cand1 was present and Nub1 absent, suggesting that NEDDylation is being regulated by Cand1 (Fig. 4). However, in the cortex at 10 dpi, the phenotype reversed with the presence of Nub1, and absence of Cand1, suggesting that NEDDylation is impaired by the Nub1-mediated degradation of NEDD8. The reduction of total NEDD8 protein would have significant implications in multiple signal transduction pathways, including NFκB and HIF1α, and suggests the signaling environment within the cortex at 10 dpi is unique. Enzymes necessary for ubiquitination and proteasomal degradation of proteins, including the ubiquitin activating enzymes (E1) that catalyze the first step in the ubiquitination reaction, and ubiquitin ligases (E3) that catalyze the transfer of ubiquitin from the E2 enzyme to the protein substrate, show similar spatial and temporal patterns. Since, E1 and E3 enzymes interact directly with the protein substrate, it is possible that varying complexes of E1, E2, and E3 enzymes exhibit unique activities to exert diverse biological functions (Table S1).

#### Host metabolism: Glycolysis and glucogenesis

Not only does signal transduction vary spatially and temporally, but the metabolic niche changes at different renal locations when comparing 4 and 10 dpi (Fig. 4 and S2). We utilized a systems biology pathway analysis tool, SIMONE, to visualize the interactions between proteins of interest (Fig. S3). These proteins were determined using external pathway mapping tools, entered into Reactome^105^ and the resulting list of proteins associated with metabolism were input into SIMONE^106^ as seed proteins. This tool constructs networks using the MAGINE framework^107^, which derives protein connection information from multiple databases (Fig. S2). By uncovering how these proteins are connected, pathways can be predicted from spatiotemporal proteomics data. These pathways were summarized and combined in Figure 5. While we observed alterations in abundance of various proteins related to different metabolic processes (discussed below), the enzymes necessary for glycolysis are generally present during infection in the abscess, interface, and cortex, suggesting glucose conversion to pyruvate. Many of the enzymes necessary for the pentose phosphate pathway that runs parallel to glycolysis and enzymes necessary to breakdown fructose that feed into glycolysis are not detected in the cortex at 10 dpi. However, the enzyme bisphosphoglycerate mutase (Bpgm) is solely present in the cortex. This enzyme is necessary to form 2,3-bisphosphoglycerate from the glycolysis intermediate 1,3-bisphophoglycerate. The absence of Bpgm in the abscess and interface is consistent with a hypoxic environment in the abscess, as 2,3-bisphosphoglycerate binds hemoglobin at a high affinity and causes a conformational change resulting in the release of oxygen^108,109^ (Fig. 5 and S2).

**Figure 5:**
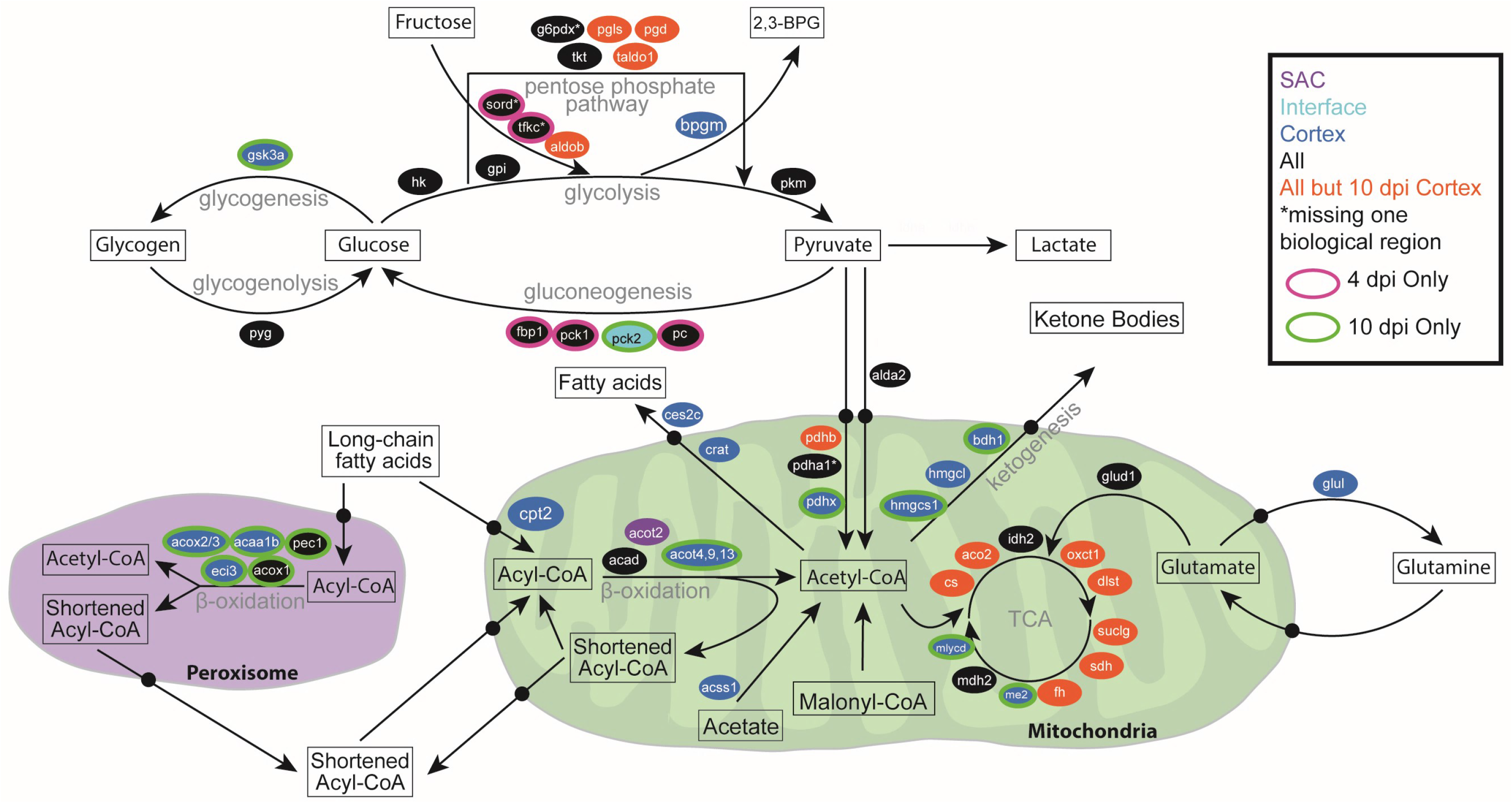
Overview of spatiotemporal distribution of proteins involved in central metabolism. Arrows denote metabolic pathways, ovals indicate genes, and colors indicate time point and region where proteins were detected.

The enzymes necessary for host gluconeogenesis, a process that converts non-carbohydrate substrates into glucose, were only detected in the abscess and cortex at 4 dpi, suggesting that gluconeogenesis may not occur late in infection (Fig. 5 and S2). The enzymes necessary for glycogenolysis are present in the abscess, interface, and cortex; however, glycogen synthase kinase-3 alpha (Gsk3a), an enzyme that contributes to glycogenesis, is not detected in the abscess and interface (Fig. 5 and S2). The lack of enzymes for glucose formation and long-term storage in the form of glycogen, and the presence of enzymes necessary for the breakdown of glycogen into glucose in the abscess and interface is consistent with high levels of glycolysis during inflammation (reviewed in^110^). The presence of Gsk3a exclusively in the cortex at 10 dpi suggests a decreased metabolic demand for glucose and possible glycogen formation. In addition, Gsk3a plays a central role in regulating the transition between pro-inflammatory and immune-suppressive response to *S. aureus* by controlling cytokine production^111^. These results suggest a specific role for Gsk3a in altering the metabolic and cytokine landscape of the cortex 10 dpi during *S. aureus* infection of the kidney.

#### Host Metabolism: Metabolism in the abscess

While the metabolic enzymes present in the abscess and interface suggest the formation of pyruvate, the complex necessary for conversion of pyruvate into acetyl coenzyme A (acetyl-CoA) in the mitochondria may not be fully formed. Pyruvate dehydrogenase protein X component (Pdhx) tethers the E3 dimers to the E2 core of the pyruvate dehydrogenate complex^112,113^ (Fig. 5 and S2). Therefore, lacking detectable Pdhx in the abscess and interface suggests that the pyruvate dehydrogenase complex would not be functional and that conversion of pyruvate into acetyl-CoA in the mitochondria would be impaired. Pyruvate can also be converted into acetyl-CoA in the cytosol^114^, but the acetyl-coenzyme A synthetase (Acss2), which is necessary for transport into the mitochondria, is also not detected within the abscess and interface (Fig. 5 and S2). Instead, the ubiquitous expression of the lactate dehydrogenase (Ldha and Ldhb) suggests that in the abscess and interface, the resulting pyruvate from glycolysis is being converted to lactate, consistent with anaerobic glycolysis in the hypoxic environment of the abscess (Fig. 5 and S2).

Many of the enzymes associated with the tricarboxylic acid (TCA) cycle are present in the abscess and interface during infection, but pyruvate does not seem to be fueling downstream oxidative phosphorylation and ATP generation due to the lack of Pdhx and Acss2 (Fig. 5 and S2). Many of the enzymes necessary for β-oxidation in the mitochondria are present in the abscess and interface, and the resulting formation of acetyl-CoA could fuel the TCA cycle (reviewed^115^). However, long-chain fatty acids (LCFAs) would likely not be able to serve as the carbon source for the TCA cycle. The O-palmitoyltransferase 2 (Cpt2), catalyzes the formation of palmitoyl-CoA from palmitoylcarnitine, a necessary step for LCFAs prior to β-oxidation (reviewed in^115^). Thus, the lack Cpt2 suggest that LCFAs cannot undergo β-oxidation in the mitochondria and therefore are not feeding into the TCA cycle in the abscess and interface (Fig. 5 and S2). This does not exclude the possibility that short-chain fatty acids (SCFAs), which passively gain access to the mitochondria, could be fueling the TCA cycle. Carnitine O-acetyltransferase (Crat) blunts acetyl-CoA from fueling the TCA cycle by forming acetyl-carnitine for transport into the cytosol, and once in the cytosol, acylcarnitine hydrolase (Ces2c) liberates fatty acids from L-carnitine. Both Crat and Ces2c are absent in the in the abscess and interface supporting the possibility that SCFAs undergoing β-oxidation could serve as a carbon source for the TCA cycle (Fig. 5 and S2). Glutamine may also act as an alternative carbon source for the TCA cycle ^116^. Glutamate dehydrogenase 1 (Glud1) catalyzes the oxidative deamination of glutamate into α-ketoglutarate, which feeds into the TCA cycle. The presence of Glud1 and absence of glutamine synthase (Glul), which converts glutamate into glutamine, in the abscess and interface suggests that glutamine may also be fueling the TCA cycle in the abscess environment (Fig. 5 and S2).

#### Host Metabolism: Metabolism in the cortex

By contrast, in the cortex at 10 dpi many of the enzymes necessary to run the TCA cycle are not detected. Only isocitrate dehydrogenase (Idh2), NAD-dependent malic enzyme (Me2), and malate dehydrogenase (Mdh2) are detected (Fig. 5 and S2), suggesting that in the cortex at 10 dpi, the full TCA cycle may not be used. The cortex at 10 dpi is the only region to contain all the acyl-CoA dehydrogenases as well as almost exclusive presence of the enzymes necessary for β-oxidation in the peroxisome. Unlike the abscess and interface, Cpt2, is detected in the cortex suggesting that β-oxidation of LCFAs may occur, but the presence of Ces2c and Crat suggest that the resulting acetyl-CoA is shunted out of the mitochondria rather than feeding into the TCA cycle. M2 macrophages in the tissue surrounding the abscess (Fig. 5 and S2) are necessary for the conversion to the resolution phase by cleaning up apoptotic cells and cellular debris, and are consistent with previous studies assessing abscess biology in the skin and soft tissue^80^. Cellular debris contains high concentrations of lipids, thereby requiring β-oxidation for its degradation. The high presence of lipids in resolving damaged tissue outside the abscess could account for the increased presence of proteins necessary for β-oxidation.

Acetyl-coenzyme A synthetase 2-like (Acss1) and malonyl-CoA decarboxylase (Mlycd) are also present in the cortex 10 days-post infection (Fig. 5 and S2). These enzymes shunt mitochondrial acetate and malonyl-CoA away from lipid synthesis and toward the formation of acetyl-CoA. The combined activity of β-oxidation, Acss1, and Mlycd with a limited presence of TCA cycle enzymes could cause an accumulation of acetyl-CoA. The presence of Crat and Cesc2c may relieve some of the acetyl-CoA burden by exporting acetyl-CoA as free fatty acids into the cytosol. Alternatively, many of the enzymes necessary for ketogenesis are present in the cortex at 10 dpi suggesting that the abundance of acetyl-CoA could also be converted to ketone bodies (Fig. 5 and S2).

A byproduct of β-oxidation is oxidative stress, and consistent with increased β-oxidation, the cortex at 10 dpi is the only region to express the full array of glutathione S-transferases, synthase, reductase, and peroxidase to combat oxidative stress (Table S1). Since oxidative phosphorylation from the TCA cycle can create oxidative stress, this could explain the absence of TCA cycle enzymes in the cortex at 10 dpi (Fig. 5 and S2). Finally, many of the downstream enzymes for β-oxidation in the mitochondria and peroxisome are not detected in the cortex at 10 dpi that are present at 4 days or in the abscess and interface (Fig. 5 and S2). This suggests that while β-oxidation of LCFAs is occurring in the cortex at 10 dpi, the lipids are not fully broken down. In addition, acyl-coenzyme A thioesterases (Acot) are thought to maintain a sufficient CoA pool for β-oxidation by terminating β-oxidation after a set number of cycles (reviewed in^117^). The specific presence of Acot4 in the peroxisome and Acot9 and Acot13 in the mitochondria could result in the formation of specific lipid products that is important in maintaining β-oxidation in high lipid environments in order to avoid oxidative stress (Fig. 5 and S2). The metabolic environment in the cortex at 10 dpi is consistent with an immune response that has altered to a resolving phase that is cleaning up lipid-dense apoptotic cells and cellular debris following inflammation.

## Summary

We performed spatially and temporally targeted proteomics, aided by a robust systems biology data processing workflow, to molecularly investigate staphylococcal abscess formation and development. This approach is suited to reliably identify bacterial proteins as well as enables characterization of the host proteome during infection. Such a comprehensive assessment of the entire abscess proteome allows us to molecularly characterize the host-pathogen interface over time. By pairing information about the spatiotemporal distribution of bacterial proteins with data defining the host proteome, we can elucidate how pathogen and host shape the abscess (micro)environment and how, in turn, both parties react to these biomolecular changes.

Using our microLESA workflow, we characterized the metabolic niche at the site of infection. Our findings indicate the influx of immune cells (i.e. neutrophils and different macrophage populations), while also elucidating specific metabolic processes employed by these host cell populations. The action of macrophages was not limited to the SAC and interface. We also detected markers for M2 macrophages in the renal cortex, suggesting a role in tissue repair and remodeling. In contrast, multiple neutrophil-specific antimicrobial factors (e.g. Mpo, Ctsg, Elane) were found to be expressed in proximity to the staphylococcal abscess. We found *S. aureus* produces CHIPS and FLIPr, which antagonize the host neutrophil receptors to prevent neutrophilic entrance into the abscess. Staphylococcal leukotoxins (LukAB) produced during abscess formation were also identified, again aiding in immune evasion and persistence in the host tissue. Indicative for the onset of nutritional immunity, we observed bacterial heme uptake and biosynthetic proteins present in the abscess. This limitation could be explained by the presence of host proteins that restrict the availability of heme for *S. aureus* (Hpx and Hp), particularly in the SAC and interface.

We describe several other additional mechanisms of how host and pathogen directly affect each other during infection and how both sides counteract the challenges presented to them. These data highlight the powerful nature of our experimental setup and offer insights into the processes at the host-pathogen interface, beyond the specific examples discussed here. The ability to elucidate when, where and how pathogens and immune systems interact during abscess formation and disease progression, is an invaluable resource to identify potential points of intervention when developing new anti-staphylococcal therapeutic strategies.

## Materials and Methods

### Murine model of systemic S. aureus infection

6-8 week old female C67BL/6J mice were anesthetized with tribromoethanol (Avertin) and retro-orbitally infected with ~1.5-2 × 10^7^ CFUs of *S. aureus* USA300 LAC constitutively expressing sfGFP from the genome (P*sarA*-sfGFP integrated at the SaPI1 site)^22^. Infections were allowed to progress for 4 or 10 days before animals were humanely euthanized and organs removed for subsequent analysis.

### Chemicals

Acetonitrile, acetic acid, formic acid, trifluoracetic acid, ethanol, ammonium bicarbonate, and chloroform were purchased from Fisher Scientific (Pittsburgh, PA). Mass spectrometry sequence-grade trypsin from porcine pancreas was purchased from Sigma-Aldrich Chemical, Co. (St. Louis, MO).

### Micro-Digestions

All sample preparation was completed using the method previously described by Ryan *et al.*^22^. Excised murine kidneys infected with *S. aureus* USA300 LAC carrying a constitutive fluorescent reporter were fresh frozen on dry ice and stored at −80°C. Tissues were then cryosectioned at 10 μm thickness (Leica Microsystems, Buffalo Grove, IL) and thaw mounted onto custom fiducial microscope slides (Delta Technology, Loveland, CO). For all samples, an autofluorescence microscopy image (Carl Zeiss Microscopy, White Plains, NY) was acquired at 10x magnification (0.92 μm/pixel) prior to spotting using FITC and DAPI filters (FITC excitation λ, 465-495 nm; emission λ, 515-555 nm; DAPI excitation λ, 340-380 nm; emission λ, 435-485 nm). Microscopy exposure time was set to 200 ms for the DAPI filter and 100 ms for the FITC filter. Samples were washed with graded ethanol washes for 30s (70% EtOH, 100% EtOH, Carnoy’s Fluid, 100% EtOH, H2O, 100% EtOH) and Carnoy’s fluid for 3min (6 ethanol: 3 chloroform: 1 acetic acid) to remove salts and lipids. Samples were allowed to dry under vacuum prior to trypsin digestion. Regions targeted for digestion were annotated on microscopy images using ImageJ (U.S. National Institutes of Health, Bethesda, MD) and converted into instrument coordinates for the piezoelectric spotter using an in-house script (Fig. S1). Trypsin was dissolved in ddH_2_O to a final concentration of 0.048 μg/mL. A piezoelectric spotting system (sciFLEXARRAYER S3, Princeton, NJ) was used to dispense ~250 pL droplets of trypsin. Trypsin was dispensed on regions of interest (ROIs) 20 times with one drop per run in order to reduce spot size, with 4 spots per ROI. The four trypsin spots were positioned so that all tryptic peptides for an ROI could be collected using a single LESA experiment. Following trypsin deposition, samples were incubated at 37°C for three hours in 300 μL ammonium bicarbonate.

### LESA

Liquid surface extraction was completed using a TriVersa NanoMate (Advion Inc., Ithaca, NY) with the LESAplusLC modification. Digested samples were scanned using a flatbed scanner and uploaded to the Advion ChipSoft software. 5 μL of extraction solvent (2:8 acetonitrile/water with 0.1% formic acid) was aspirated into the glass capillary. The capillary was then lowered to a height of approximately 0.5 mm above the sample surface and 2.5 μL of solvent was dispensed onto ROIs. Contact with the surface was maintained for 10s and 3.0 μL of solvent was re-aspirated into the capillary. The initial 5 μL volume was dispensed into a 96-well plate containing 200 μL of water/0.1% formic acid. The LESA extraction was repeated twice at the same ROI and combined. Three wash cycles of the instrument were completed between each ROI set to prevent carryover from other biological regions. Resulting extracts were then dried down under vacuum and stored at −80°C until LC-MS/MS analysis.

### LC-MS/MS

Dried peptide samples were reconstituted in 10 μL of water/0.1% formic acid prior to analysis. Tryptic peptides from tissue extracts were injected and gradient eluted on a pulled tip emitter column packed in-house with C18 material (Waters BEH C18). The column was heated to 60°C with a flow rate of 400 nL/min using an Easy-nLC 1000 UHPLC (Thermo Scientific, San Jose, CA). Mobile phase A consisted of H_2_O with 0.1% formic acid, and mobile phase B consisted of acetonitrile with 0.1% formic acid. Peptides were eluted on a linear gradient of 2-20% B for 100 minutes, followed by 20-32% B for 20 minutes, and lastly 32-95% B for one minute. Eluting peptides were analyzed using an Orbitrap Fusion Tribrid mass spectrometer (Thermo Scientific, San Jose, CA). MS1 scans were acquired at 120,000 resolving power at *m/z* 200 using the Orbitrap, with a mass range of *m/z* 400-1600, and an automatic gain control target of 1.0×10^6^.

### Data Analysis

For peptide identification, tandem mass spectra were searched using Protalizer software (Vulcan Analytic, Birmingham, Alabama) and MaxQuant^118^ against a database containing both the mouse and *S. aureus* strain USA300 LAC proteome created from the UniprotKB^119^. Modifications such as glycosylation, phosphorylation, methionine, oxidation, and deamidation were included in the Protalizer database search with an FDR of 1%. For MaxQuant analysis, raw files were processed using a label-free quantification method implemented in MaxQuant version 1.6.7. Spectra were searched against mouse and *Staphylococcus aureus* (strain USA300) reference databases downloaded from UniProt KB. These were supplemented with the reversed sequences and common contaminants automatically (decoy database) and used for quality control and estimation of FDR by MaxQuant. Carbamidomethylation was set as a fixed modification and acetyl (protein N-term) and oxidation (M) were set as variable modifications. Minimal peptide length was 7 amino acids. Peptide and protein FDRs were both set to 1%. The resultant protein groups file from MaxQuant was analyzed for outliers by calculating z-scores for each sample based on number of protein groups identified; 3 samples out of 42 were excluded as outliers. Proteins identified as “reverse”, “only identified by site”, or “potential contaminants” were also removed.

Proteins identified by Protalizer and MaxQuant were filtered based on the following criteria: 2 unique peptides contributed to the protein identification, proteins were detected in 2+ biological replicates, and proteins were detected in 2+ serial sections. Bacterial proteins also met these criteria, with the exception of those listed in Table S3 (designated as ‘lower confidence’) that only met 1 of the 2 replicate requirements criteria but did meet the 2+ unique peptide criteria. We believe these proteins did not fulfil the requirements for high confidence due to potentially relevant differences in abundance or due to heterogeneity between abscesses. Since these proteins have previously been studied in a primarily targeted manner rather than our discovery-based approach, we include these data but additional confirmation is required. The lists generated by each dataset were cross compared for all proteins, to aid in identification from the protein groups generated by MaxQuant. Tables S1 and S2 include the full list of identified proteins, and include the search algorithm(s) were used for identification

Pathway analysis was conducted using a similar workflow described previously by Gutierrez and colleagues^106^. Briefly, data were uploaded into a central in-house database and organized by identifiers such as tissue location and infection time. Project data were exported into a custom data analysis and visualization tool for network construction (SIMONE) in a data driven manner. Data was entered into Reactome^105^, a free online pathway mapping software, and proteins associated with metabolism were selected from the Reactome output. The genes encoding for these proteins were then input into the network construction software as root nodes. Network construction was performed as described^107^ in conjunction with a custom user interface that provided visualization (through the use of Cytoscape24) of the constructed data networks. Resulting networks are shown in Fig. S1 and S2.

## Data and Software Availability

The mass spectrometry proteomics data have been deposited to the ProteomeXchange Consortium via the PRIDE^120^ partner repository with the dataset identifier PXD019920.

## Acknowledgements

This work was funded by the NIH National Institute of Allergy and Infectious Diseases (R01AI138581 awarded to E.P.S and J.M.S. and R01AI069233, R01AI073843 awarded to E.P.S.) and NIH National Institute of General Medical Sciences (2P41 GM103391-07 awarded to R.M.C.). A.W. is supported by American Heart Association (18POST33990262) and the NIH National Institute of Environmental Health Sciences (T32ES007028).

## Author Contributions

E.R.G., A.W., A.J.M, D.B.G., R.M.C., E.P.S., J.M.S wrote the manuscript. A.W. performed animal infection models. W.J.P aided with sample preparation. D.J.R. designed experiments and collected data. E.R.G. and K.S. performed data analysis and pathway-mapping results.

---

Figure S1: Examples of microLESA from different regions. Locations of microLESA sampling (colored dots) are shown on autofluorescence images of infected kidneys (10 dpi). Regions are shown by color-coded circles.

Figure S2: Spatiotemporal distribution of metabolic proteins during infection. Blue circles denote proteins that were present at the specified timepoint/region, while grey circles depict the absence of a specific protein.

Table S1: List of host-derived proteins found by presence at time point and biological region, including platform used for identification (P = Protalizer, MQ = MaxQuant).

Table S2: List of bacterial proteins detected and their localizations based on stringent search criteria from Protalizer and MaxQuant. No bacterial proteins were detected in the cortex.

Table S3: List of bacterial proteins detected and their localizations with lower identification confidence.

