## Supplemental Figures and Tables for "Spatially-targeted proteomics of the host-pathogen interface during staphylococcal abscess formation"

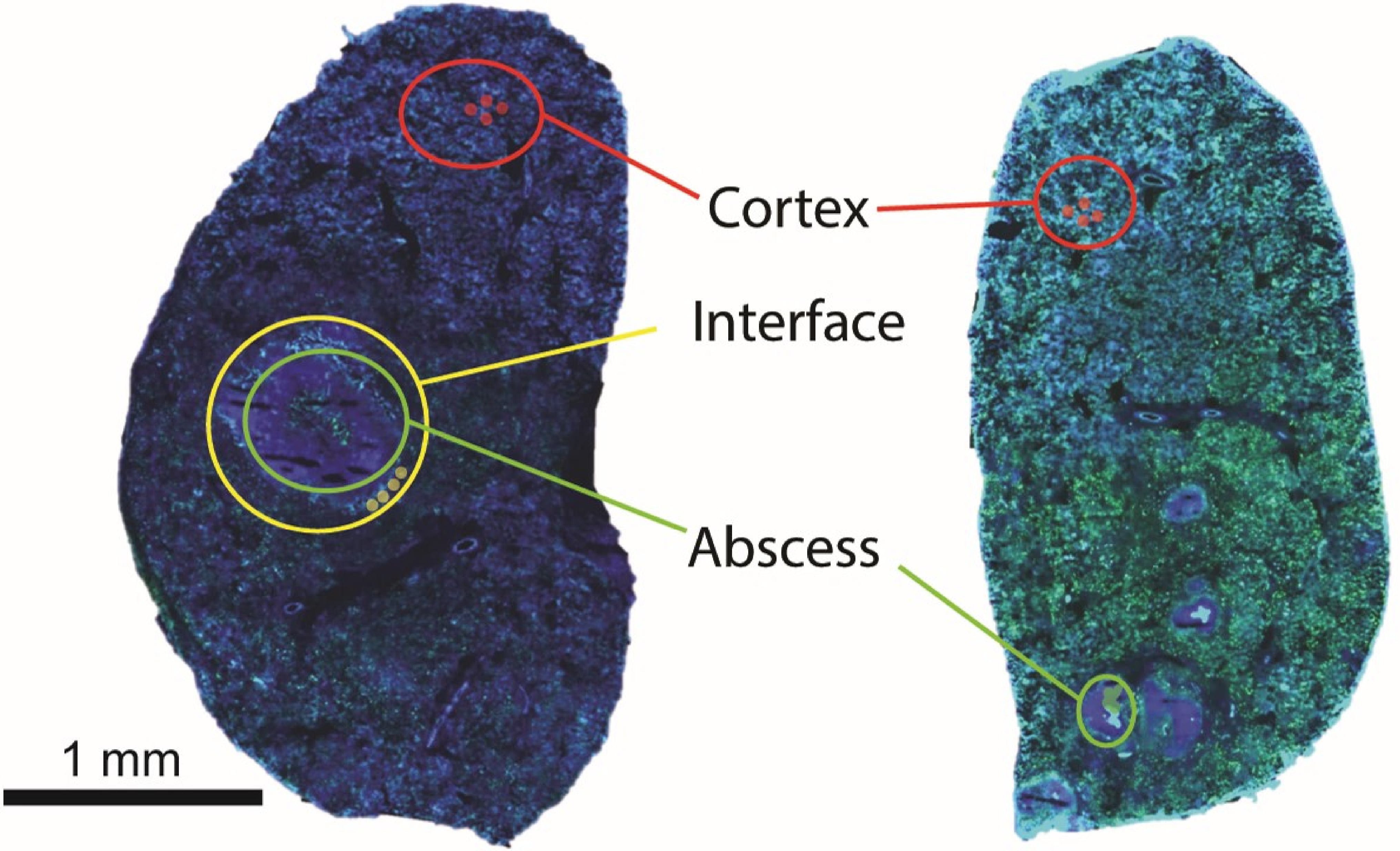
Cortex
Interface
Abscess
1 mm

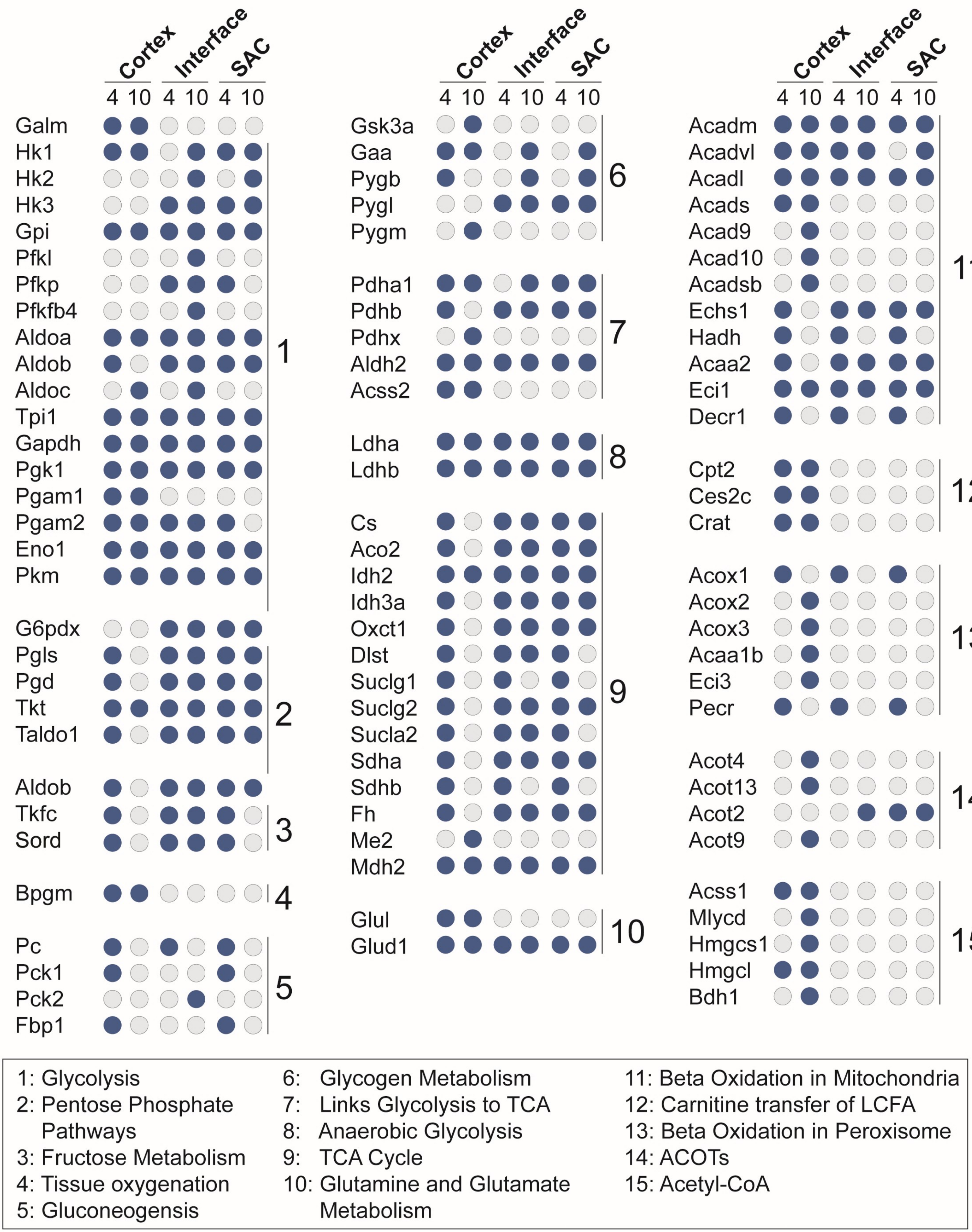
Cortex
Interface
SAC
4 10 4 10 4 10
Galm
Hk1
Hk2
Hk3
Gpi
Pfkl
Pfkp
Pfkfb4
Aldoa
Aldob
Aldoc
Tpi1
Gapdh
Pgk1
Pgam1
Pgam2
Eno1
Pkm
1
G6pdx
Pgls
Pgd
Tkt
Taldo1
2
Aldob
Tkfc
Sord
3
Bpgm
4
Pc
Pck1
Pck2
Fbp1
5
Gsk3a
Gaa
Pygb
Pygl
Pygm
6
Pdha1
Pdhb
Pdhx
Aldh2
Acss2
7
Ldha
Ldhb
8
Cs
Aco2
Idh2
Idh3a
Oxct1
Dlst
Suclg1
Suclg2
Sucla2
Sdha
Sdhb
Fh
Me2
Mdh2
9
Glul
Glud1
10
Acadm
Acadvl
Acadl
Acads
Acad9
Acad10
Acadsb
Echs1
Hadh
Acaa2
Eci1
Decr1
11
Cpt2
Ces2c
Crat
12
Acox1
Acox2
Acox3
Acaa1b
Eci3
Pecr
13
Acot4
Acot13
Acot2
Acot9
14
Acss1
Mlycd
Hmgcs1
Hmgcl
Bdh1
15
1: Glycolysis
2: Pentose Phosphate Pathways
3: Fructose Metabolism
4: Tissue oxygenation
5: Gluconeogensis
6: Glycogen Metabolism
7: Links Glycolysis to TCA
8: Anaerobic Glycolysis
9: TCA Cycle
10: Glutamine and Glutamate Metabolism
11: Beta Oxidation in Mitochondria
12: Carnitine transfer of LCFA
13: Beta Oxidation in Peroxisome
14: ACOTs
15: Acetyl-CoA

| Protein Name | Ascension Number | 4 dpi Co | 10 dpi Co | 4 dpi INT | 10 dpi INT | 4 dpi SAC | 10 dpi SAC | Platform |
|---|---|---|---|---|---|---|---|---|
| HK2 Hexokinase-2 | O08528 | NO | NO | NO | YES | NO | YES | P |
| GSTO1 Glutathione S-transferase omega-1 | O09131 | YES | YES | YES | YES | YES | YES | P/MQ |
| PGAM2 Phosphoglycerate mutase 2 | O70250 | YES | YES | YES | YES | YES | NO | P/MQ |
| ALDOC Fructose-bisphosphate aldolase C | P05063 | NO | YES | NO | YES | NO | NO | P |
| ALDOA Fructose-bisphosphate aldolase A | P05064 | YES | YES | YES | YES | YES | YES | P/MQ |
| G6PI Glucose-6-phosphate isomerase | P06745 | YES | YES | YES | YES | YES | YES | P/MQ |
| MDH2 Malate dehydrogenase, mitochondrial | P08249 | YES | YES | YES | YES | YES | YES | P/MQ |
| PGK1 Phosphoglycerate kinase 1 | P09411 | YES | YES | YES | YES | YES | YES | P/MQ |
| GSTA2 Glutathione S-transferase A2 | P10648 | NO | YES | NO | NO | YES | NO | P/MQ |
| GSTM1 Glutathione S-transferase Mu 1 | P10649 | YES | YES | YES | YES | YES | YES | P/MQ |
| GPX1 Glutathione peroxidase 1 | P11352 | YES | YES | YES | YES | YES | YES | P/MQ |
| PFKL ATP-dependent 6-phosphofructokinase, liver type | P12382 | NO | NO | NO | YES | NO | NO | P |
| GSTA1 Glutathione S-transferase A1 | P13745 | NO | YES | YES | NO | NO | NO | P/MQ |
| GADPH Glyceraldehyde-3-phosphate dehydrogenase | P16858 | YES | YES | YES | YES | YES | YES | P/MQ |
| Eno1 Alpha-enolase | P17182 | YES | YES | YES | YES | YES | YES | P/MQ |
| HK1 Hexokinase-1 | P17710 | YES | YES | NO | YES | YES | YES | P/MQ |
| TPI1 Triosephosphate isomerase | P17751 | YES | YES | YES | YES | YES | YES | P/MQ |
| GSTP1 Glutathione S-transferase P 1 | P19157 | YES | YES | YES | YES | YES | YES | P/MQ |
| GSTA4 Glutathione S-transferase A4 | P24472 | YES | YES | NO | NO | NO | NO | P |
| GSTA3 Glutathione S-transferase A3 | P30115 | YES | YES | YES | NO | YES | NO | P |
| TKT Transketolase | P40142 | YES | YES | YES | YES | YES | YES | P/MQ |
| ECI1 Enoyl-CoA delta isomerase 1, mitochondrial | P42125 | YES | YES | YES | YES | YES | YES | P/MQ |
| ACADM Medium-chain specific acyl-CoA dehydrogenase, mitochondrial | P45952 | YES | YES | YES | YES | YES | YES | P/MQ |
| GPX3 Glutathione peroxidase 3 | P46412 | YES | YES | NO | YES | YES | NO | P/MQ |
| GSHR Glutathione reductase, mitochondrial | P47791 | YES | YES | YES | YES | YES | YES | P/MQ |
| ACADVL Very long-chain specific acyl-CoA dehydrogenase, mitochondrial | P50544 | YES | YES | YES | YES | NO | YES | P/MQ |
| ACADL Long-chain specific acyl-CoA dehydrogenase, mitochondrial | P51174 | YES | YES | YES | YES | YES | YES | P/MQ |
| GSHB Glutathione synthetase | P51855 | YES | YES | NO | NO | NO | NO | P |
| KPYM Pyruvate kinase PKM | P52480 | YES | YES | YES | YES | YES | YES | P/MQ |
| IDH2 Isocitrate dehydrogenase [NADP], mitochondrial | P54071 | YES | YES | YES | YES | YES | YES | P/MQ |
| FH Fumarate hydratase, mitochondrial | P97807 | YES | NO | YES | YES | YES | YES | P/MQ |
| G6PDX Glucose-6-phosphate 1-dehydrogenase X | Q00612 | NO | NO | YES | YES | YES | YES | P/MQ |
| ACADS Short-chain specific acyl-CoA dehydrogenase, mitochondrial | Q07417 | YES | YES | NO | NO | NO | NO | P |
| HK3 Hexokinase-3 | Q3TRM8 | NO | NO | YES | YES | YES | YES | P/MQ |

| | | | | | | | | |
|---|---|---|---|---|---|---|---|---|
| GSTT2 Glutathione S-transferase theta-2 | Q61133 | NO | YES | NO | NO | NO | NO | P |
| HADH Hydroxyacyl-coenzyme A dehydrogenase, mitochondrial | Q61425 | YES | NO | YES | NO | YES | NO | P |
| ECI3 Enoyl-CoA delta isomerase 3, peroxisomal | Q78JN3 | NO | YES | NO | NO | NO | NO | P |
| ECHM Enoyl-CoA hydratase, mitochondrial | Q8BH95 | YES | NO | YES | YES | YES | YES | P/MQ |
| ACAA2 3-ketoacyl-CoA thiolase, mitochondrial | Q8BWT1 | YES | NO | YES | YES | YES | YES | P/MQ |
| ACAD9 Acyl-CoA dehydrogenase family member 9, mitochondrial | Q8JZN5 | NO | YES | NO | NO | NO | NO | P |
| ACAD10 Acyl-CoA dehydrogenase family member 10 | Q8K370 | NO | YES | NO | NO | NO | NO | P |
| ACAA1B 3-ketoacyl-CoA thiolase B, peroxisomal | Q8VCH0 | NO | YES | NO | NO | NO | NO | P/MQ |
| ALDOB Fructose-bisphosphate aldolase B | Q91Y97 | YES | NO | YES | YES | YES | YES | P/MQ |
| ACO2 Aconitate hydratase, mitochondrial | Q99KI0 | YES | NO | YES | YES | YES | YES | P/MQ |
| GSTT3 Glutathione S-transferase theta-3 | Q99L20 | NO | YES | NO | NO | NO | NO | P |
| PECR Peroxisomal trans-2-enoyl-CoA reductase | Q99MZ7 | YES | NO | YES | NO | YES | NO | P/MQ |
| PGLS 6-phosphogluconolactonase | Q9CQ60 | YES | NO | YES | YES | YES | YES | P/MQ |
| DECR1 2,4-dienoyl-CoA reductase, mitochondrial | Q9CQ62 | YES | NO | YES | NO | YES | NO | P/MQ |
| CS Citrate synthase, mitochondrial | Q9CZU6 | YES | NO | YES | YES | YES | YES | P/MQ |
| OXCT1 Succinyl-CoA:3-ketoacid coenzyme A transferase 1, mitochondrial | Q9D0K2 | YES | NO | YES | YES | YES | YES | P/MQ |
| PGAM1 Phosphoglycerate mutase 1 | Q9DBJ1 | YES | NO | YES | YES | YES | YES | P/MQ |
| ACADSB Short/branched chain specific acyl-CoA dehydrogenase, mitochondrial | Q9DBL1 | NO | YES | NO | NO | NO | NO | P |
| PGD 6-phosphogluconate dehydrogenase, decarboxylating | Q9DCD0 | YES | NO | YES | YES | YES | YES | P/MQ |
| GSTK1 Glutathione S-transferase kappa 1 | Q9DCM2 | NO | YES | NO | NO | NO | NO | P |
| ACOX3 Peroxisomal acyl-coenzyme A oxidase 3 | Q9EPL9 | NO | YES | NO | NO | NO | NO | P |
| ACOX2 Peroxisomal acyl-coenzyme A oxidase 2 | Q9QXD1 | NO | YES | NO | NO | NO | NO | P |
| ACOX1 Peroxisomal acyl-coenzyme A oxidase 1 | Q9R0H0 | YES | NO | YES | NO | YES | NO | P/MQ |
| PFKP ATP-dependent 6-phosphofructokinase, platelet type | Q9WUA3 | NO | NO | YES | YES | YES | NO | P/MQ |
| SUCLG1 Succinate--CoA ligase [ADP/GDP-forming] subunit alpha, mitochondrial | Q9WUM5 | YES | NO | YES | NO | YES | NO | P |
| SUCLG2 Succinate--CoA ligase [GDP-forming] subunit beta, mitochondrial | Q9Z2I8 | YES | NO | YES | YES | YES | YES | P/MQ |
| SUCLA2 Succinate--CoA ligase [ADP-forming] subunit beta, mitochondrial | Q9Z2I9 | YES | NO | NO | YES | YES | NO | P |
| Immunoglobulin heavy constant mu | A0A075B6A0 | YES | YES | YES | YES | YES | YES | MQ |
| Fibronectin | A0A087WSN6 | YES | YES | YES | YES | YES | YES | MQ |
| FCGR4 Low affinity immunoglobulin gamma Fc region receptor IV | A0A0B4J1G0 | NO | NO | NO | YES | NO | NO | P |
| Histone H2A | A0A0N4SV66 | YES | YES | YES | YES | YES | YES | MQ |
| Guanine nucleotide-binding protein subunit gamma | A0A0N4SVT3 | YES | YES | YES | YES | YES | YES | MQ |
| Leucine-rich repeat flightless-interacting protein 2 | A0A0R4J169 | YES | YES | YES | YES | YES | YES | MQ |
| S-adenosylmethionine synthase | A0A0U1RNT6 | YES | YES | YES | YES | YES | YES | MQ |

| | | | | | | | | |
|---|---|---|---|---|---|---|---|---|
| Phosphatidylinositol-binding clathrin assembly protein | A0A140LHQ8 | YES | YES | YES | YES | YES | YES | MQ |
| HIG1 domain family member 1A, mitochondrial | A0A1L1SR69 | YES | YES | YES | YES | YES | YES | MQ |
| MAPK-interacting and spindle-stabilizing protein-like | A0A2I3BRS9 | YES | YES | YES | YES | YES | YES | MQ |
| Kininogen 2 | A0A338P699 | YES | YES | YES | YES | YES | YES | MQ |
| High mobility group protein HMG-I/HMG-Y | A0A338P6G6 | YES | YES | YES | YES | YES | YES | MQ |
| Myosin-11 | A0A338P6K2 | YES | YES | YES | YES | YES | YES | MQ |
| Receptor expression-enhancing protein 5 | A0A494BBE3 | YES | YES | YES | YES | YES | YES | MQ |
| K1522 Uncharacterized protein KIAA1522 | A2A7S8 | NO | YES | NO | NO | NO | NO | P |
| SPD2B SH3 and PX domain-containing protein 2B | A2AAY5 | NO | NO | NO | YES | NO | NO | P |
| DDI2 Protein DDI1 homolog 2 | A2ADY9 | YES | YES | YES | YES | YES | YES | P/MQ |
| CKAP5 Cytoskeleton-associated protein 5 | A2AGT5 | NO | NO | NO | YES | YES | YES | P |
| MA7D1 MAP7 domain-containing protein 1 | A2AJI0 | NO | YES | NO | YES | NO | NO | P |
| QCR2 Cytochrome b-c1 complex subunit 2, mitochondrial | A2AKK5 | YES | YES | NO | NO | NO | NO | P |
| UBR4 E3 ubiquitin-protein ligase UBR4 | A2AN08 | NO | NO | NO | NO | YES | NO | P |
| LRP2 Low-density lipoprotein receptor-related protein 2 | A2ARV4 | YES | YES | YES | NO | YES | NO | P/MQ |
| AGRIN Agrin | A2ASQ1 | NO | YES | YES | YES | YES | NO | P |
| ARI1A AT-rich interactive domain-containing protein 1A | A2BH40 | NO | NO | NO | YES | YES | NO | P |
| RPN2 Dolichyl-diphosphooligosaccharide--protein glycosyltransferase subunit 2 | A3KMP2 | YES | YES | NO | NO | NO | NO | P |
| MMRN2 Multimerin-2 | A6H6E2 | NO | YES | NO | NO | NO | NO | P |
| ITIH4 Inter alpha-trypsin inhibitor, heavy chain 4 | A6X935 | YES | YES | YES | YES | YES | YES | P/MQ |
| THOC2 THO complex subunit 2 | B1AZI6 | NO | NO | NO | YES | NO | NO | P |
| ANR44 Serine/threonine-protein phosphatase 6 regulatory ankyrin repeat subunit B | B2RXR6 | NO | NO | NO | YES | NO | NO | P |
| PLXB2 Plexin-B2 | B2RXS4 | NO | YES | NO | NO | NO | NO | P |
| RBM25 RNA-binding protein 25 | B2RY56 | NO | NO | YES | YES | NO | NO | P |
| OSBL8 Oxysterol-binding protein-related protein 8 | B9EJ86 | NO | YES | NO | YES | NO | NO | P |
| H2A1D Histone H2A type 1-D | C0HKE3 | NO | NO | NO | YES | NO | NO | P/MQ |
| RNT2A Ribonuclease T2-A | C0HKG5 | NO | YES | NO | NO | NO | NO | P |
| SAFB1 Scaffold attachment factor B1 | D3YXK2 | NO | NO | NO | YES | YES | NO | P |
| CCDC6 Coiled-coil domain-containing protein 6 | D3YZP9 | NO | NO | NO | NO | YES | NO | P |
| BIN2 Bridging integrator 2 | D3Z6Q9 | NO | YES | YES | YES | YES | YES | P/MQ |
| PGAM1 Phosphoglycerate mutase 1 | D3Z7P3 | YES | YES | NO | NO | NO | NO | P |
| FIBA Fibrinogen alpha chain | E9PV24 | YES | YES | YES | YES | YES | YES | P/MQ |
| GCN1 eIF-2-alpha kinase activator GCN1 | E9PVA8 | NO | YES | NO | YES | NO | NO | P |
| KI67 Proliferation marker protein Ki-67 | E9PVX6 | NO | NO | NO | YES | YES | YES | P |
| PARP4 Protein mono-ADP-ribosyltransferase PARP4 | E9PYK3 | NO | YES | NO | YES | NO | NO | P |
| AKP13 A-kinase anchor protein 13 | E9Q394 | NO | YES | NO | YES | YES | NO | P |
| APOB Apolipoprotein B-100 | E9Q414 | NO | YES | NO | NO | NO | NO | P |

| | | | | | | | | |
|---|---|---|---|---|---|---|---|---|
| RN213 E3 ubiquitin-protein ligase RNF213 | E9Q555 | NO | NO | YES | YES | YES | YES | P |
| DESP Desmoplakin | E9Q557 | NO | YES | NO | NO | NO | NO | P |
| NOLC1 Nucleolar and coiled-body phosphoprotein 1 | E9Q5C9 | NO | NO | NO | YES | NO | NO | P |
| BD1L1 Biorientation of chromosomes in cell division protein 1-like 1 | E9Q6J5 | NO | NO | NO | YES | NO | NO | P |
| CDHR2 Cadherin-related family member 2 | E9Q7P9 | NO | YES | NO | NO | NO | NO | P |
| SC16A Protein transport protein Sec16A | E9QAT4 | NO | YES | NO | YES | NO | NO | P |
| TPR Nucleoprotein TPR | F6ZDS4 | YES | YES | YES | YES | YES | YES | P |
| MORC3 MORC family CW-type zinc finger protein 3 | F7BJB9 | NO | NO | NO | YES | NO | NO | P |
| MYPT1 Protein phosphatase 1 regulatory subunit 12A | F8VPU2 | YES | YES | NO | NO | NO | NO | P/MQ |
| AT2B1 Plasma membrane calcium-transporting ATPase 1 | G5E829 | YES | YES | YES | YES | NO | YES | P |
| TRIPC E3 ubiquitin-protein ligase TRIP12 | G5E870 | NO | NO | NO | YES | NO | NO | P |
| ANK3 Ankyrin-3 | G5E8K5 | YES | YES | YES | NO | YES | NO | P/MQ |
| Tropomyosin alpha-1 chain | G5E8R2 | YES | YES | YES | YES | YES | YES | MQ |
| Signal recognition particle 19 kDa protein | G5E8T3 | YES | YES | YES | YES | YES | YES | MQ |
| ADP-ribosylation factor-interacting protein 1 | G5E8V9 | YES | YES | YES | YES | YES | YES | MQ |
| AP complex subunit beta | H3BKM0 | YES | YES | YES | YES | YES | YES | MQ |
| CAN2 Calpain-2 catalytic subunit | O08529 | YES | YES | YES | YES | YES | YES | P |
| BIN1 Myc box-dependent-interacting protein 1 | O08539 | NO | NO | NO | YES | NO | NO | P |
| SC22B Vesicle-trafficking protein SEC22b | O08547 | NO | YES | NO | YES | YES | YES | P/MQ |
| DPYL2 Dihydropyrimidinase-related protein 2 | O08553 | YES | YES | YES | YES | YES | YES | P/MQ |
| LEG9 Galectin-9 | O08573 | NO | YES | NO | NO | NO | NO | P |
| EMD Emerin | O08579 | NO | NO | NO | YES | NO | NO | P |
| THOC4 THO complex subunit 4 | O08583 | YES | NO | YES | YES | YES | YES | P/MQ |
| CLCA Clathrin light chain A | O08585 | YES | YES | YES | YES | YES | YES | P/MQ |
| NPL N-acetylneuraminate lyase | O08600 | YES | YES | NO | NO | NO | NO | P |
| MTP Microsomal triglyceride transfer protein large subunit | O08601 | NO | YES | NO | NO | NO | NO | P |
| MYH11 Myosin-11 | O08638 | YES | YES | YES | NO | YES | YES | P |
| MAP2 Methionine aminopeptidase 2 | O08663 | NO | NO | NO | YES | NO | NO | P |
| ARGI2 Arginase-2, mitochondrial | O08691 | NO | YES | NO | NO | NO | NO | P |
| NGP Neutrophilic granule protein | O08692 | YES | YES | YES | YES | YES | YES | P/MQ |
| PRDX6 Peroxiredoxin-6 | O08709 | YES | YES | YES | YES | YES | YES | P/MQ |
| DLDH Dihydrolipoyl dehydrogenase, mitochondrial | O08749 | YES | YES | YES | YES | YES | YES | P/MQ |
| ATP5H ATP synthase subunit d, mitochondrial | O08756 | YES | YES | NO | NO | YES | NO | P |
| UBE3A Ubiquitin-protein ligase E3A | O08759 | NO | YES | NO | NO | NO | NO | P |
| TCOF Treacle protein | O08784 | NO | YES | NO | YES | YES | YES | P/MQ |
| GLU2B Glucosidase 2 subunit beta | O08795 | YES | YES | YES | YES | YES | YES | P/MQ |
| DIAP1 Protein diaphanous homolog 1 | O08808 | NO | NO | YES | YES | YES | YES | P |
| U5S1 116 kDa U5 small nuclear ribonucleoprotein component | O08810 | YES | YES | NO | NO | YES | YES | P |

| | | | | | | | | |
|---|---|---|---|---|---|---|---|---|
| AIP AH receptor-interacting protein | O08915 | NO | YES | YES | YES | YES | YES | P |
| FLOT1 Flotillin-1 | O08917 | NO | NO | NO | YES | YES | YES | P/MQ |
| S22A1 Solute carrier family 22 member 1 | O08966 | NO | YES | NO | NO | NO | NO | P |
| SDCB1 Syntenin-1 | O08992 | NO | NO | NO | NO | YES | YES | P/MQ |
| ATOX1 Copper transport protein ATOX1 | O08997 | YES | YES | YES | YES | YES | YES | P |
| NAPSA Napsin-A | O09043 | NO | YES | NO | NO | NO | NO | P |
| SNP23 Synaptosomal-associated protein 23 | O09044 | YES | YES | NO | YES | NO | YES | P/MQ |
| PSB1 Proteasome subunit beta type-1 | O09061 | NO | NO | YES | YES | YES | YES | P |
| NDUBB NADH dehydrogenase [ubiquinone] 1 beta subcomplex subunit 11, mitochondrial | O09111 | NO | YES | NO | NO | NO | NO | P |
| MA2B1 Lysosomal alpha-mannosidase | O09159 | NO | YES | NO | YES | NO | YES | P/MQ |
| SODE Extracellular superoxide dismutase [Cu-Zn] | O09164 | NO | YES | NO | YES | YES | NO | P |
| RL21 60S ribosomal protein L21 | O09167 | YES | YES | YES | YES | YES | YES | P/MQ |
| GSH0 Glutamate--cysteine ligase regulatory subunit | O09172 | NO | NO | NO | YES | YES | NO | P |
| HGD Homogentisate 1,2-dioxygenase | O09173 | NO | YES | NO | NO | NO | NO | P |
| AMACR Alpha-methylacyl-CoA racemase | O09174 | YES | YES | NO | NO | NO | NO | P |
| PHB2 Prohibitin-2 | O35129 | YES | YES | YES | YES | YES | YES | P/MQ |
| ATIF1 ATPase inhibitor, mitochondrial | O35143 | YES | YES | YES | YES | YES | YES | P |
| COFA1 Collagen alpha-1(XV) chain | O35206 | NO | YES | NO | YES | NO | NO | P |
| DOPD D-dopachrome decarboxylase | O35215 | YES | YES | YES | NO | YES | NO | P/MQ |
| PSMD4 26S proteasome non-ATPase regulatory subunit 4 | O35226 | NO | YES | YES | YES | NO | NO | P |
| DHX15 Pre-mRNA-splicing factor ATP-dependent RNA helicase DHX15 | O35286 | YES | YES | YES | YES | YES | YES | P/MQ |
| PURB Transcriptional activator protein Pur-beta | O35295 | YES | YES | NO | YES | YES | YES | P/MQ |
| SRSF5 Serine/arginine-rich splicing factor 5 | O35326 | NO | YES | NO | NO | NO | NO | P |
| CAN1 Calpain-1 catalytic subunit | O35350 | NO | YES | YES | YES | YES | YES | P/MQ |
| AN32A Acidic leucine-rich nuclear phosphoprotein 32 family member A | O35381 | YES | YES | YES | YES | YES | YES | P/MQ |
| EXOC4 Exocyst complex component 4 | O35382 | NO | YES | NO | YES | NO | NO | P/MQ |
| FOLH1 Glutamate carboxypeptidase 2 | O35409 | NO | YES | NO | NO | NO | NO | P |
| ECH1 Delta(3,5)-Delta(2,4)-dienoyl-CoA isomerase, mitochondrial | O35459 | YES | YES | NO | NO | YES | NO | P/MQ |
| FKBP8 Peptidyl-prolyl cis-trans isomerase FKBP8 | O35465 | NO | YES | NO | NO | NO | NO | P |
| S27A2 Very long-chain acyl-CoA synthetase | O35488 | YES | YES | NO | NO | YES | NO | P |
| PSDE 26S proteasome non-ATPase regulatory subunit 14 | O35593 | NO | YES | NO | YES | YES | NO | P/MQ |
| FYB1 FYN-binding protein 1 | O35601 | NO | NO | NO | YES | NO | YES | P/MQ |
| SCAM3 Secretory carrier-associated membrane protein 3 | O35609 | NO | YES | NO | YES | YES | NO | P |
| ANXA3 Annexin A3 | O35639 | YES | YES | YES | YES | YES | YES | P/MQ |
| AP1B1 AP-1 complex subunit beta-1 | O35643 | NO | YES | NO | YES | NO | NO | P |
| NEUR1 Sialidase-1 | O35657 | NO | YES | NO | NO | NO | NO | P |

| | | | | | | | | |
|---|---|---|---|---|---|---|---|---|
| C1QBP Complement component 1 Q subcomponent-binding protein, mitochondrial | O35658 | NO | NO | NO | NO | YES | NO | P/MQ |
| NUDC Nuclear migration protein nudC | O35685 | YES | YES | YES | YES | YES | YES | P/MQ |
| PININ Pinin | O35691 | NO | YES | NO | NO | NO | NO | P |
| SPTC1 Serine palmitoyltransferase 1 | O35704 | NO | YES | NO | NO | NO | NO | P |
| HNRH1 Heterogeneous nuclear ribonucleoprotein H | O35737 | NO | NO | YES | YES | YES | YES | P/MQ |
| CHIL3 Chitinase-like protein 3 | O35744 | YES | YES | YES | YES | YES | YES | P/MQ |
| API5 Apoptosis inhibitor 5 | O35841 | NO | YES | NO | YES | YES | YES | P |
| BCAT2 Branched-chain-amino-acid aminotransferase, mitochondrial | O35855 | NO | YES | NO | NO | NO | NO | P |
| TIM44 Mitochondrial import inner membrane translocase subunit TIM44 | O35857 | NO | YES | NO | YES | YES | NO | P |
| CALU Calumenin | O35887 | YES | YES | YES | YES | YES | YES | P |
| SP100 Nuclear autoantigen Sp-100 | O35892 | NO | YES | NO | YES | NO | NO | P/MQ |
| PSB10 Proteasome subunit beta type-10 | O35955 | NO | YES | YES | YES | NO | YES | P/MQ |
| RM23 39S ribosomal protein L23, mitochondrial | O35972 | NO | YES | NO | NO | NO | NO | P |
| SDC4 Syndecan-4 | O35988 | YES | YES | NO | NO | NO | NO | P/MQ |
| CAVN1 Caveolae-associated protein 1 | O54724 | YES | YES | YES | YES | NO | NO | P/MQ |
| OST48 Dolichyl-diphosphooligosaccharide--protein glycosyltransferase 48 kDa subunit | O54734 | YES | YES | YES | YES | YES | YES | P |
| AP3D1 AP-3 complex subunit delta-1 | O54774 | NO | NO | NO | YES | NO | YES | P |
| IL16 Pro-interleukin-16 | O54824 | NO | NO | NO | YES | YES | YES | P |
| HMGB3 High mobility group protein B3 | O54879 | NO | YES | NO | YES | YES | YES | P |
| REPS1 RalBP1-associated Eps domain-containing protein 1 | O54916 | NO | YES | NO | NO | NO | NO | P |
| AKAP2 A-kinase anchor protein 2 | O54931 | YES | YES | YES | YES | YES | YES | P |
| MTG16 Protein CBFA2T3 | O54972 | NO | NO | NO | YES | NO | NO | P |
| ASNA ATPase Asna1 | O54984 | NO | NO | NO | YES | NO | NO | P |
| SLK STE20-like serine/threonine-protein kinase | O54988 | YES | YES | YES | YES | YES | YES | P/MQ |
| PROM1 Prominin-1 | O54990 | NO | YES | NO | NO | NO | NO | P |
| TPPC3 Trafficking protein particle complex subunit 3 | O55013 | NO | YES | NO | NO | NO | NO | P |
| PGRC1 Membrane-associated progesterone receptor component 1 | O55022 | YES | YES | NO | YES | NO | NO | P |
| IMPA1 Inositol monophosphatase 1 | O55023 | YES | YES | YES | YES | YES | YES | P |
| COPB2 Coatomer subunit beta' | O55029 | YES | YES | YES | YES | YES | YES | P |
| SYUA Alpha-synuclein | O55042 | YES | YES | NO | NO | YES | NO | P |
| STK10 Serine/threonine-protein kinase 10 | O55098 | NO | NO | YES | YES | YES | YES | P/MQ |
| STRN Striatin | O55106 | NO | YES | NO | YES | YES | YES | P |
| O55125 | O55125 | YES | NO | NO | NO | NO | NO | P |
| SEPT7 Septin-7 | O55131 | YES | YES | YES | YES | YES | YES | P/MQ |
| IF6 Eukaryotic translation initiation factor 6 | O55135 | NO | YES | YES | YES | YES | NO | P/MQ |
| RL35A 60S ribosomal protein L35a | O55142 | YES | YES | NO | NO | YES | YES | P |

| | | | | | | | | |
|---|---|---|---|---|---|---|---|---|
| AT2A2 Sarcoplasmic/endoplasmic reticulum calcium ATPase 2 | O55143 | YES | YES | YES | YES | YES | YES | P/MQ |
| SPT5H Transcription elongation factor SPT5 | O55201 | NO | NO | NO | YES | NO | NO | P |
| ILK Integrin-linked protein kinase | O55222 | NO | NO | YES | YES | YES | YES | P |
| PSB5 Proteasome subunit beta type-5 | O55234 | NO | YES | NO | NO | NO | NO | P |
| DHX9 ATP-dependent RNA helicase A | O70133 | NO | YES | YES | YES | YES | YES | P/MQ |
| MMP8 Neutrophil collagenase | O70138 | NO | NO | NO | YES | NO | YES | P |
| NCF2 Neutrophil cytosol factor 2 | O70145 | NO | NO | YES | YES | YES | YES | P/MQ |
| AIF1 Allograft inflammatory factor 1 | O70200 | NO | NO | NO | YES | NO | NO | P |
| EF1B Elongation factor 1-beta | O70251 | YES | YES | NO | YES | YES | YES | P/MQ |
| NMT1 Glycylpeptide N-tetradecanoyltransferase 1 | O70310 | NO | NO | NO | YES | YES | YES | P |
| E41L2 Band 4.1-like protein 2 | O70318 | YES | YES | YES | YES | YES | YES | P/MQ |
| RNH1 Ribonuclease H1 | O70338 | NO | YES | NO | NO | NO | NO | P |
| PDLI1 PDZ and LIM domain protein 1 | O70400 | NO | YES | NO | NO | YES | NO | P |
| VAMP8 Vesicle-associated membrane protein 8 | O70404 | NO | YES | YES | YES | YES | YES | P/MQ |
| FHL2 Four and a half LIM domains protein 2 | O70433 | NO | YES | NO | NO | NO | NO | P |
| PSA3 Proteasome subunit alpha type-3 | O70435 | NO | YES | YES | YES | YES | YES | P/MQ |
| STX7 Syntaxin-7 | O70439 | YES | YES | YES | YES | YES | YES | P/MQ |
| 1433S 14-3-3 protein sigma | O70456 | NO | NO | NO | NO | YES | NO | P |
| UGDH UDP-glucose 6-dehydrogenase | O70475 | YES | YES | NO | YES | YES | YES | P/MQ |
| SNX3 Sorting nexin-3 | O70492 | YES | YES | YES | YES | YES | YES | P/MQ |
| SRPK1 SRSF protein kinase 1 | O70551 | NO | YES | NO | NO | NO | NO | P |
| PIGR Polymeric immunoglobulin receptor | O70570 | NO | NO | NO | NO | YES | NO | P |
| NSMA Sphingomyelin phosphodiesterase 2 | O70572 | NO | YES | NO | NO | NO | NO | P |
| S22A2 Solute carrier family 22 member 2 | O70577 | YES | YES | NO | NO | NO | NO | P |
| CSKP Peripheral plasma membrane protein CASK | O70589 | NO | YES | NO | NO | NO | NO | P |
| PFD2 Prefoldin subunit 2 | O70591 | NO | YES | NO | YES | NO | YES | P/MQ |
| CFDP1 Craniofacial development protein 1 | O88271 | NO | NO | NO | YES | YES | YES | P |
| ZN326 DBIRD complex subunit ZNF326 | O88291 | NO | YES | NO | NO | NO | NO | P |
| SORL1 Sortilin-related receptor | O88307 | NO | YES | NO | NO | NO | NO | P |
| NID2 Nidogen-2 | O88322 | YES | YES | NO | YES | YES | YES | P/MQ |
| CAD16 Cadherin-16 | O88338 | YES | YES | YES | NO | YES | YES | P/MQ |
| WDR1 WD repeat-containing protein 1 | O88342 | YES | YES | YES | YES | YES | YES | P/MQ |
| S4A4 Electrogenic sodium bicarbonate cotransporter 1 | O88343 | YES | YES | NO | NO | NO | NO | P/MQ |
| VTI1B Vesicle transport through interaction with t-SNAREs homolog 1B | O88384 | NO | NO | NO | YES | YES | YES | P/MQ |
| PAPS2 Bifunctional 3'-phosphoadenosine 5'-phosphosulfate synthase 2 | O88428 | YES | YES | NO | NO | YES | NO | P |
| CPNS1 Calpain small subunit 1 | O88456 | YES | YES | NO | YES | NO | NO | P/MQ |
| DC1I2 Cytoplasmic dynein 1 intermediate chain 2 | O88487 | YES | YES | YES | YES | YES | YES | P/MQ |
| NEMO NF-kappa-B essential modulator | O88522 | NO | NO | NO | YES | NO | NO | P |
| PPT1 Palmitoyl-protein thioesterase 1 | O88531 | NO | YES | NO | YES | NO | YES | P |

| | | | | | | | | |
|---|---|---|---|---|---|---|---|---|
| ZFR Zinc finger RNA-binding protein | O88532 | NO | NO | NO | YES | NO | NO | P/MQ |
| CSN4 COP9 signalosome complex subunit 4 | O88544 | YES | YES | YES | YES | NO | YES | P/MQ |
| ROA2 Heterogeneous nuclear ribonucleoproteins A2/B1 | O88569 | YES | YES | YES | YES | YES | YES | P/MQ |
| COMT Catechol O-methyltransferase | O88587 | NO | YES | NO | NO | NO | NO | P |
| PRS6A 26S proteasome regulatory subunit 6A | O88685 | NO | YES | YES | YES | YES | YES | P |
| CTBP1 C-terminal-binding protein 1 | O88712 | NO | YES | NO | NO | NO | NO | P |
| TOM1 Target of Myb protein 1 | O88746 | NO | NO | NO | YES | YES | NO | P |
| IDHC Isocitrate dehydrogenase [NADP] cytoplasmic | O88844 | YES | YES | YES | YES | YES | YES | P/MQ |
| LIN7C Protein lin-7 homolog C | O88952 | YES | YES | NO | NO | NO | NO | P |
| GNPI1 Glucosamine-6-phosphate isomerase 1 | O88958 | NO | YES | NO | NO | NO | NO | P |
| KBL 2-amino-3-ketobutyrate coenzyme A ligase, mitochondrial | O88986 | NO | YES | NO | NO | NO | NO | P |
| COR1A Coronin-1A | O89053 | YES | YES | YES | YES | YES | YES | P/MQ |
| COPE Coatomer subunit epsilon | O89079 | NO | YES | NO | YES | YES | NO | P/MQ |
| RBM3 RNA-binding protein 3 | O89086 | NO | YES | YES | YES | YES | YES | P |
| ADH1 Alcohol dehydrogenase 1 | P00329 | YES | YES | YES | NO | NO | NO | P |
| DYR Dihydrofolate reductase | P00375 | NO | YES | NO | NO | NO | NO | P |
| COX2 Cytochrome c oxidase subunit 2 | P00405 | YES | YES | YES | NO | NO | NO | P |
| HPRT Hypoxanthine-guanine phosphoribosyltransferase | P00493 | NO | NO | NO | NO | YES | NO | P |
| CAH2 Carbonic anhydrase 2 | P00920 | YES | YES | YES | YES | YES | YES | P/MQ |
| CO3 Complement C3 | P01027 | YES | YES | YES | YES | YES | YES | P/MQ |
| CO4B Complement C4-B | P01029 | YES | YES | NO | NO | YES | YES | P |
| IGJ Immunoglobulin J chain | P01592 | YES | YES | NO | NO | YES | NO | P |
| KV2A7 Ig kappa chain V-II region 26-10 | P01631 | NO | YES | NO | NO | NO | NO | P |
| KV5A3 Ig kappa chain V-V region K2 (Fragment) | P01635 | NO | YES | NO | NO | NO | NO | P |
| KV5A9 Ig kappa chain V-V region L7 (Fragment) | P01642 | NO | YES | NO | NO | NO | NO | P |
| KV5AA Ig kappa chain V-V region MOPC 173 | P01643 | NO | YES | NO | NO | NO | NO | P |
| KV3A1 Ig kappa chain V-III region PC 2880/PC 1229 | P01654 | NO | YES | NO | NO | NO | NO | P |
| KV3AI Ig kappa chain V-III region PC 6684 | P01670 | NO | NO | NO | YES | NO | NO | P |
| LV1A Ig lambda-1 chain V region | P01723 | NO | YES | NO | NO | NO | NO | P |
| HVM28 Ig heavy chain V-III region U61 | P01797 | NO | YES | NO | YES | NO | NO | P |
| IGKC Immunoglobulin kappa constant | P01837 | YES | YES | NO | YES | YES | YES | P |
| GCAB Ig gamma-2A chain C region secreted form | P01864 | NO | YES | NO | YES | NO | NO | P |
| IGG2B Ig gamma-2B chain C region | P01867 | NO | YES | NO | NO | NO | NO | P |
| IGHM Immunoglobulin heavy constant mu | P01872 | YES | YES | YES | YES | YES | YES | P/MQ |
| B2MG Beta-2-microglobulin | P01887 | NO | YES | NO | NO | NO | YES | P |
| HA11 H-2 class I histocompatibility antigen, D-B alpha chain | P01899 | NO | NO | YES | YES | YES | YES | P |
| HA12 H-2 class I histocompatibility antigen, D-D alpha chain | P01900 | NO | YES | NO | NO | NO | NO | P |

| | | | | | | | | |
|---|---|---|---|---|---|---|---|---|
| HA1B H-2 class I histocompatibility antigen, K-B alpha chain | P01901 | NO | YES | YES | YES | YES | NO | P |
| HB2D H-2 class II histocompatibility antigen, A-D beta chain | P01921 | NO | YES | NO | NO | NO | NO | P |
| HBA Hemoglobin subunit alpha | P01942 | YES | YES | YES | YES | YES | YES | P/MQ |
| HBB-b1 Hemoglobin subunit beta-1 | P02088 | YES | YES | YES | YES | YES | YES | P/MQ |
| HBB-b2 Hemoglobin subunit beta-2 | P02089 | NO | YES | NO | NO | NO | NO | P/MQ |
| LAMC1 Laminin subunit gamma-1 | P02468 | YES | YES | NO | YES | YES | YES | P/MQ |
| LAMB1 Laminin subunit beta-1 | P02469 | NO | YES | NO | NO | NO | NO | P |
| ADA Adenosine deaminase | P03958 | NO | NO | NO | YES | NO | NO | P |
| IGHG3 Ig gamma-3 chain C region | P03987 | NO | YES | NO | YES | NO | YES | P |
| FABP4 Fatty acid-binding protein, adipocyte | P04117 | NO | YES | NO | YES | YES | NO | P/MQ |
| CFAB Complement factor B | P04186 | YES | YES | YES | YES | YES | YES | P/MQ |
| HG2A H-2 class II histocompatibility antigen gamma chain | P04441 | NO | YES | NO | YES | NO | YES | P |
| HBB-h0 Hemoglobin subunit beta-H0 | P04443 | NO | YES | NO | NO | NO | NO | P |
| B3AT Band 3 anion transport protein | P04919 | YES | YES | YES | YES | YES | NO | P/MQ |
| AATC Aspartate aminotransferase, cytoplasmic | P05201 | YES | YES | YES | YES | YES | YES | P/MQ |
| AATM Aspartate aminotransferase, mitochondrial | P05202 | YES | YES | YES | YES | YES | YES | P/MQ |
| SAA1 Serum amyloid A-1 protein | P05366 | YES | YES | YES | YES | YES | YES | P/MQ |
| SAA2 Serum amyloid A-2 protein | P05367 | YES | YES | YES | NO | YES | NO | P |
| MAC-1 Integrin alpha-M | P05555 | NO | NO | YES | YES | YES | YES | P |
| K1C18 Keratin, type I cytoskeletal 18 | P05784 | YES | YES | YES | YES | YES | YES | P |
| LDHA L-lactate dehydrogenase A chain | P06151 | YES | YES | YES | YES | YES | YES | P/MQ |
| RENI1 Renin-1 | P06281 | NO | YES | NO | NO | NO | NO | P |
| APOBR Apolipoprotein A-IV | P06728 | YES | YES | YES | YES | YES | YES | P/MQ |
| CATL1 Cathepsin L1 | P06797 | NO | YES | NO | NO | NO | NO | P |
| CD45 Receptor-type tyrosine-protein phosphatase C | P06800 | NO | NO | YES | YES | YES | YES | P |
| MAOX NADP-dependent malic enzyme | P06801 | NO | YES | NO | NO | NO | NO | P |
| CFAH Complement factor H | P06909 | YES | YES | NO | NO | NO | YES | P |
| TTHY Transthyretin | P07309 | YES | YES | NO | YES | NO | YES | P |
| ANXA2 Annexin A2 | P07356 | YES | YES | YES | YES | YES | YES | P/MQ |
| A1AG2 Alpha-1-acid glycoprotein 2 | P07361 | YES | YES | NO | YES | YES | YES | P |
| ALBU Serum albumin | P07724 | YES | YES | YES | YES | YES | YES | P/MQ |
| A1AT1 Alpha-1-antitrypsin 1-1 | P07758 | NO | YES | NO | NO | NO | NO | P |
| SPA3K Serine protease inhibitor A3K | P07759 | YES | YES | YES | YES | YES | YES | P/MQ |
| HS90A Heat shock protein HSP 90-alpha | P07901 | YES | YES | YES | YES | YES | YES | P/MQ |
| PDIA4 Protein disulfide-isomerase A4 | P08003 | YES | YES | YES | YES | YES | YES | P/MQ |
| APT Adenine phosphoribosyltransferase | P08030 | NO | YES | NO | YES | YES | YES | P |
| SPTA1 Spectrin alpha chain, erythrocytic 1 | P08032 | YES | YES | NO | NO | YES | NO | P/MQ |
| TRFL Lactotransferrin | P08071 | YES | YES | YES | YES | YES | YES | P/MQ |
| HCK Tyrosine-protein kinase HCK | P08103 | NO | NO | NO | YES | YES | YES | P |

| | | | | | | | | |
|---|---|---|---|---|---|---|---|---|
| ENPL Endoplasmin | P08113 | YES | YES | YES | YES | YES | YES | P/MQ |
| CO3A1 Collagen alpha-1(III) chain | P08121 | NO | NO | NO | NO | YES | NO | P |
| CO4A2 Collagen alpha-2(IV) chain | P08122 | YES | YES | YES | YES | YES | YES | P/MQ |
| S10AA Protein S100-A10 | P08207 | NO | YES | NO | YES | YES | NO | P |
| APOE Apolipoprotein E | P08226 | YES | YES | YES | YES | YES | YES | P/MQ |
| SODC Superoxide dismutase [Cu-Zn] | P08228 | YES | YES | YES | NO | YES | YES | P/MQ |
| RASN GTPase NRas | P08556 | NO | NO | NO | YES | NO | NO | P |
| GNAI2 Guanine nucleotide-binding protein G(i) subunit alpha-2 | P08752 | YES | YES | YES | YES | YES | YES | P/MQ |
| RPB1 DNA-directed RNA polymerase II subunit RPB1 | P08775 | NO | YES | NO | YES | NO | YES | P/MQ |
| LYZ2 Lysozyme C-2 | P08905 | YES | YES | NO | YES | YES | YES | P/MQ |
| ISK1 Serine protease inhibitor Kazal-type 1 | P09036 | NO | YES | NO | NO | NO | NO | P |
| ITB1 Integrin beta-1 | P09055 | YES | YES | YES | YES | YES | YES | P/MQ |
| PDIA1 Protein disulfide-isomerase | P09103 | YES | YES | YES | YES | YES | YES | P/MQ |
| NUCL Nucleolin | P09405 | YES | YES | YES | YES | YES | YES | P/MQ |
| JUNB Transcription factor jun-B | P09450 | NO | NO | NO | YES | NO | NO | P |
| FRIH Ferritin heavy chain | P09528 | YES | YES | YES | YES | YES | YES | P/MQ |
| HMGN2 non-histone chromosomal protein HMG-17 | P09602 | NO | YES | YES | YES | YES | YES | P/MQ |
| SODM Superoxide dismutase [Mn], mitochondrial | P09671 | YES | YES | NO | YES | YES | YES | P/MQ |
| CADH1 Cadherin-1 | P09803 | YES | YES | NO | NO | NO | NO | P/MQ |
| APOA2 Apolipoprotein A-II | P09813 | YES | YES | YES | YES | YES | YES | P/MQ |
| H2AZ Histone H2A.Z | P0C0S6 | YES | YES | NO | YES | YES | YES | P/MQ |
| IFI5B Interferon-activable protein 205-B | P0DOV1 | NO | YES | NO | YES | NO | YES | P/MQ |
| CALM3 Calmodulin-3 | P0DP28 | YES | YES | YES | YES | YES | YES | P/MQ |
| Annexin A1 | P10107 | YES | YES | YES | YES | YES | YES | MQ |
| EF1A1 Elongation factor 1-alpha 1 | P10126 | YES | YES | YES | YES | YES | YES | P/MQ |
| CO4B Complement C4-B | P1029 | NO | NO | YES | NO | NO | NO | P |
| NID1 Nidogen-1 | P10493 | YES | YES | YES | YES | YES | YES | P/MQ |
| HEM2 Delta-aminolevulinic acid dehydratase | P10518 | NO | YES | NO | NO | YES | NO | P |
| TAU Microtubule-associated protein tau | P10637 | YES | YES | NO | NO | YES | NO | P/MQ |
| THIO Thioredoxin | P10639 | YES | YES | YES | YES | YES | YES | P/MQ |
| TCEA1 Transcription elongation factor A protein 1 | P10711 | NO | NO | YES | YES | YES | YES | P/MQ |
| CD14 Monocyte differentiation antigen CD14 | P10810 | NO | NO | YES | YES | YES | YES | P |
| 4F2 4F2 cell-surface antigen heavy chain | P10852 | YES | YES | YES | YES | YES | YES | P/MQ |
| H2B1M Histone H2B type 1-M | P10854 | NO | YES | NO | NO | NO | NO | P |
| H10 Histone H1.0 | P10922 | YES | YES | YES | YES | YES | YES | P/MQ |
| OSTP Osteopontin | P10923 | YES | YES | YES | YES | YES | YES | P/MQ |
| TCP4 Activated RNA polymerase II transcriptional coactivator p15 | P11031 | NO | YES | NO | YES | YES | YES | P |
| CO1A1 Collagen alpha-1(I) chain | P11087 | NO | YES | NO | YES | YES | YES | P/MQ |
| MPO Myeloperoxidase | P11247 | NO | YES | YES | YES | YES | YES | P |
| FABPH Fatty acid-binding protein, heart | P11404 | YES | YES | NO | NO | NO | NO | P |

| | | | | | | | | |
|---|---|---|---|---|---|---|---|---|
| HS90B Heat shock protein HSP 90-beta | P11499 | YES | YES | YES | YES | YES | YES | P/MQ |
| NGAL Neutrophil gelatinase-associated lipocalin | P11672 | YES | YES | YES | YES | YES | YES | P/MQ |
| K2C8 Keratin, type II cytoskeletal 8 | P11679 | YES | YES | YES | YES | YES | YES | P/MQ |
| ITB2 Integrin beta-2 | P11835 | NO | YES | YES | YES | YES | YES | P/MQ |
| ANGT Angiotensinogen | P11859 | NO | YES | NO | NO | NO | NO | P |
| GAS2 Growth arrest-specific protein 2 | P11862 | YES | YES | NO | NO | NO | NO | P |
| NUD19 Nucleoside diphosphate-linked moiety X motif 19 | P11930 | NO | YES | NO | NO | NO | NO | P |
| TCPA T-complex protein 1 subunit alpha | P11983 | YES | YES | YES | YES | YES | YES | P/MQ |
| A4 Amyloid-beta A4 protein | P12023 | NO | YES | NO | NO | NO | NO | P/MQ |
| SAMP Serum amyloid P-component | P12246 | YES | YES | YES | YES | YES | YES | P/MQ |
| BGLR Beta-glucuronidase | P12265 | NO | NO | NO | YES | NO | NO | P |
| CALB1 Calbindin | P12658 | YES | YES | YES | NO | YES | NO | P |
| FABPL Fatty acid-binding protein, liver | P12710 | NO | YES | NO | NO | NO | NO | P |
| COX5A Cytochrome c oxidase subunit 5A, mitochondrial | P12787 | YES | YES | YES | YES | YES | YES | P/MQ |
| RL7A 60S ribosomal protein L7a | P12970 | YES | YES | YES | YES | YES | YES | P/MQ |
| GELS Gelsolin | P13020 | YES | YES | YES | YES | YES | YES | P/MQ |
| UMPS Uridine 5'-monophosphate synthase | P13439 | NO | YES | NO | NO | NO | NO | P |
| Intercellular adhesion molecule 1 | P13597 | YES | YES | YES | YES | YES | YES | MQ |
| CAH1 Carbonic anhydrase 1 | P13634 | NO | NO | NO | NO | YES | NO | P |
| GPDA Glycerol-3-phosphate dehydrogenase [NAD(+)], cytoplasmic | P13707 | YES | YES | NO | NO | NO | NO | P |
| S10A6 Protein S100-A6 | P14069 | NO | YES | NO | YES | YES | NO | P |
| AT1B1 Sodium/potassium-transporting ATPase subunit beta-1 | P14094 | YES | YES | YES | NO | YES | NO | P/MQ |
| RL27A 60S ribosomal protein L27a | P14115 | YES | YES | YES | YES | YES | YES | P/MQ |
| RS16 40S ribosomal protein S16 | P14131 | YES | YES | YES | YES | YES | YES | P/MQ |
| RL7 60S ribosomal protein L7 | P14148 | YES | YES | YES | YES | YES | YES | P/MQ |
| MDH2 Malate dehydrogenase, cytoplasmic | P14152 | YES | YES | YES | YES | YES | YES | P/MQ |
| RSSA 40S ribosomal protein SA | P14206 | YES | YES | YES | YES | YES | YES | P/MQ |
| CALR Calreticulin | P14211 | YES | YES | YES | YES | YES | YES | P/MQ |
| FGR Tyrosine-protein kinase Fgr | P14234 | NO | NO | NO | YES | YES | YES | P |
| HSPB1 Heat shock protein beta-1 | P14602 | YES | YES | YES | YES | YES | YES | P/MQ |
| PSMD3 26S proteasome non-ATPase regulatory subunit 3 | P14685 | YES | YES | YES | YES | YES | YES | P |
| LMNB1 Lamin-B1 | P14733 | YES | YES | YES | YES | YES | YES | P/MQ |
| ANXA6 Annexin A6 | P14824 | YES | YES | YES | YES | YES | YES | P/MQ |
| RLA0 60S acidic ribosomal protein P0 | P14869 | YES | YES | YES | YES | YES | YES | P/MQ |
| HMOX1 Heme oxygenase 1 | P14901 | NO | NO | YES | YES | YES | YES | P |
| GLUL Glutamine synthetase | P15105 | YES | YES | NO | NO | NO | NO | P/MQ |
| PMGE Bisphosphoglycerate mutase | P15327 | YES | YES | NO | NO | NO | NO | P |
| CD44 CD44 antigen | P15379 | NO | YES | NO | NO | NO | NO | P |

| | | | | | | | | |
|---|---|---|---|---|---|---|---|---|
| NDKA Nucleoside diphosphate kinase A | P15532 | NO | YES | NO | NO | NO | NO | P/MQ |
| LEUK Leukosialin | P15702 | NO | NO | NO | YES | NO | NO | P |
| H12 Histone H1.2 | P15864 | YES | YES | NO | NO | YES | YES | P/MQ |
| KLK1 Kallikrein-1 | P15947 | YES | YES | NO | NO | YES | NO | P |
| LEG1 Galectin-1 | P16045 | NO | YES | NO | NO | NO | YES | P |
| LEG3 Galectin-3 | P16110 | YES | YES | YES | YES | YES | YES | P/MQ |
| LDHB L-lactate dehydrogenase B chain | P16125 | YES | YES | YES | YES | YES | YES | P/MQ |
| PH4H Phenylalanine-4-hydroxylase | P16331 | YES | YES | NO | NO | NO | NO | P |
| MUTA Methylmalonyl-CoA mutase, mitochondrial | P16332 | YES | YES | NO | NO | NO | NO | P |
| AMPE Glutamyl aminopeptidase | P16406 | YES | YES | YES | NO | YES | NO | P/MQ |
| Argininosuccinate synthase | P16460 | YES | YES | YES | YES | YES | YES | MQ |
| SPTN1 Spectrin alpha chain, non-erythrocytic 1 | P16546 | YES | YES | YES | YES | YES | YES | P |
| PPGB Lysosomal protective protein | P16675 | NO | YES | NO | YES | NO | NO | P |
| HMGA1 High mobility group protein HMG-I/HMG-Y | P17095 | NO | YES | NO | YES | YES | NO | P/MQ |
| PTBP1 Polypyrimidine tract-binding protein 1 | P17225 | YES | YES | YES | YES | YES | YES | P/MQ |
| AP2A1 AP-2 complex subunit alpha-1 | P17426 | NO | NO | NO | YES | NO | NO | P |
| AP2A2 AP-2 complex subunit alpha-2 | P17427 | NO | YES | NO | YES | NO | NO | P/MQ |
| SBP1 Methanethiol oxidase | P17563 | YES | YES | YES | NO | YES | NO | P |
| COX7C Cytochrome c oxidase subunit 7C, mitochondrial | P17665 | NO | YES | NO | NO | NO | NO | P |
| PPIA Peptidyl-prolyl cis-trans isomerase A | P17742 | YES | YES | YES | YES | YES | YES | P/MQ |
| GTR1 Solute carrier family 2, facilitated glucose transporter member 1 | P17809 | NO | NO | NO | NO | YES | NO | P |
| PCNA Proliferating cell nuclear antigen | P17918 | NO | NO | YES | YES | YES | NO | P/MQ |
| CATD Cathepsin D | P18242 | YES | YES | YES | YES | YES | YES | P/MQ |
| BASI Basigin | P18572 | YES | YES | NO | NO | YES | YES | P/MQ |
| HMGN1 non-histone chromosomal protein HMG-14 | P18608 | YES | NO | NO | YES | YES | YES | P/MQ |
| KS6A1 Ribosomal protein S6 kinase alpha-1 | P18653 | NO | YES | NO | NO | NO | NO | P |
| KS6A3 Ribosomal protein S6 kinase alpha-3 | P18654 | YES | YES | YES | YES | YES | YES | P |
| COF1 Cofilin-1 | P18760 | YES | YES | YES | YES | YES | YES | P/MQ |
| OXDA D-amino-acid oxidase | P18894 | NO | YES | NO | NO | NO | NO | P |
| K1C19 Keratin, type I cytoskeletal 19 | P19001 | NO | YES | NO | YES | NO | YES | P |
| FAS Fatty acid synthase | P19096 | YES | YES | NO | YES | YES | YES | P/MQ |
| THRB Prothrombin | P19221 | NO | NO | YES | NO | NO | NO | P |
| RL13A 60S ribosomal protein L13a | P19253 | YES | YES | YES | YES | YES | YES | P |
| SERPH Serpin H1 | P19324 | YES | YES | YES | YES | YES | YES | P/MQ |
| COX5B Cytochrome c oxidase subunit 5B, mitochondrial | P19536 | YES | YES | YES | YES | YES | YES | P/MQ |
| COX41 Cytochrome c oxidase subunit 4 isoform 1, mitochondrial | P19783 | YES | YES | YES | YES | YES | YES | P/MQ |
| LSP1 Lymphocyte-specific protein 1 | P19973 | YES | YES | YES | YES | YES | YES | P/MQ |
| BIP Endoplasmic reticulum chaperone BiP | P20029 | YES | YES | YES | YES | YES | YES | P/MQ |

| | | | | | | | | |
|---|---|---|---|---|---|---|---|---|
| HEXB Beta-hexosaminidase subunit beta | P20060 | NO | YES | YES | YES | YES | YES | P/MQ |
| TYB4 Thymosin beta-4 | P20065 | YES | YES | YES | YES | YES | YES | P/MQ |
| PRDX3 Thioredoxin-dependent peroxide reductase, mitochondrial | P20108 | YES | YES | YES | YES | YES | YES | P/MQ |
| VIME Vimentin | P20152 | YES | YES | YES | YES | YES | YES | P/MQ |
| FCERG High affinity immunoglobulin epsilon receptor subunit gamma | P20491 | NO | NO | YES | YES | YES | YES | P |
| PLMN Plasminogen | P20918 | YES | YES | YES | YES | YES | YES | P/MQ |
| TPM3 Tropomyosin alpha-3 chain | P21107 | YES | YES | YES | YES | YES | YES | P |
| GNA11 Guanine nucleotide-binding protein subunit alpha-11 | P21278 | NO | YES | NO | NO | NO | NO | P |
| VTDB Vitamin D-binding protein | P21614 | YES | YES | YES | YES | YES | NO | P/MQ |
| LMNB2 Lamin-B2 | P21619 | YES | YES | YES | YES | YES | YES | P/MQ |
| TAP1 Antigen peptide transporter 1 | P21958 | NO | NO | NO | YES | NO | NO | P |
| TGM2 Protein-glutamine gamma-glutamyltransferase 2 | P21981 | YES | YES | YES | YES | YES | YES | P/MQ |
| EMB Embigin | P21995 | NO | YES | NO | NO | YES | NO | P |
| HEMH Ferrochelatase, mitochondrial | P22315 | NO | YES | NO | NO | NO | NO | P |
| A1AT2 Alpha-1-antitrypsin 1-2 | P22599 | YES | YES | YES | NO | YES | NO | P/MQ |
| CBL E3 ubiquitin-protein ligase CBL | P22682 | NO | YES | NO | YES | NO | YES | P |
| EIF3A Eukaryotic translation initiation factor 3 subunit A | P23116 | YES | YES | YES | YES | YES | YES | P/MQ |
| CBX3 Chromobox protein homolog 3 | P23198 | YES | YES | YES | YES | YES | YES | P/MQ |
| PNPH Purine nucleoside phosphorylase | P23492 | NO | YES | YES | YES | YES | YES | P/MQ |
| FCL GDP-L-fucose synthase | P23591 | NO | YES | NO | YES | NO | NO | P |
| BGAL Beta-galactosidase | P23780 | NO | YES | NO | YES | NO | NO | P |
| CRYAB Alpha-crystallin B chain | P23927 | NO | YES | NO | NO | NO | NO | P |
| EST1C Carboxylesterase 1C | P23953 | YES | YES | YES | YES | YES | YES | P |
| CATA Catalase | P24270 | YES | YES | YES | YES | YES | YES | P/MQ |
| PPIB Peptidyl-prolyl cis-trans isomerase B | P24369 | YES | YES | YES | YES | YES | YES | P/MQ |
| CAPG Macrophage-capping protein | P24452 | YES | YES | YES | YES | YES | YES | P/MQ |
| LKHA4 Leukotriene A-4 hydrolase | P24527 | YES | YES | YES | YES | YES | YES | P/MQ |
| IMDH2 Inosine-5'-monophosphate dehydrogenase 2 | P24547 | NO | YES | YES | YES | YES | NO | P |
| AL1A1 Retinal dehydrogenase 1 | P24549 | NO | YES | NO | NO | NO | NO | P |
| M6PR Cation-dependent mannose-6-phosphate receptor | P24668 | NO | YES | NO | NO | YES | NO | P |
| IL1RA Interleukin-1 receptor antagonist protein | P25085 | NO | NO | NO | NO | YES | NO | P |
| MCM3 DNA replication licensing factor MCM3 | P25206 | NO | NO | NO | YES | NO | YES | P/MQ |
| RS2 40S ribosomal protein S2 | P25444 | YES | YES | YES | YES | YES | YES | P/MQ |
| LYN Tyrosine-protein kinase Lyn | P25911 | NO | NO | NO | YES | YES | NO | P |
| UBF1 Nucleolar transcription factor 1 | P25976 | NO | YES | YES | YES | YES | YES | P/MQ |
| TLN1 Talin-1 | P26039 | YES | YES | YES | YES | YES | YES | P/MQ |
| EZRI Ezrin | P26040 | YES | YES | YES | YES | YES | YES | P/MQ |

| | | | | | | | | |
|---|---|---|---|---|---|---|---|---|
| MOES Moesin | P26041 | YES | YES | YES | YES | YES | YES | P/MQ |
| RADI Radixin | P26043 | YES | YES | YES | YES | YES | YES | P/MQ |
| CTNA1 Catenin alpha-1 | P26231 | YES | YES | YES | YES | YES | YES | P/MQ |
| PTMA Prothymosin alpha | P26350 | YES | YES | YES | YES | YES | YES | P/MQ |
| U2AF2 Splicing factor U2AF 65 kDa subunit | P26369 | NO | NO | NO | YES | YES | NO | P |
| GLUD1 Glutamate dehydrogenase 1, mitochondrial | P26443 | YES | YES | YES | YES | YES | YES | P/MQ |
| PSMD7 26S proteasome non-ATPase regulatory subunit 7 | P26516 | NO | YES | NO | YES | NO | YES | P |
| SYSC Serine--tRNA ligase, cytoplasmic | P26638 | YES | YES | YES | YES | YES | YES | P |
| MARCS Myristoylated alanine-rich C-kinase substrate | P26645 | NO | YES | YES | YES | YES | YES | P/MQ |
| FKB1A Peptidyl-prolyl cis-trans isomerase FKBP1A | P26883 | NO | YES | NO | YES | YES | NO | P/MQ |
| MA2A1 Alpha-mannosidase 2 | P27046 | NO | NO | NO | YES | NO | NO | P |
| MAP4 Microtubule-associated protein 4 | P27546 | YES | YES | YES | YES | YES | YES | P/MQ |
| GNA13 Guanine nucleotide-binding protein subunit alpha-13 | P27601 | NO | NO | NO | YES | NO | NO | P |
| PLAP Phospholipase A-2-activating protein | P27612 | YES | YES | NO | YES | YES | YES | P/MQ |
| XRCC5 X-ray repair cross-complementing protein 5 | P27641 | NO | YES | NO | NO | NO | NO | P |
| RL3 60S ribosomal protein L3 | P27659 | YES | YES | YES | YES | YES | YES | P/MQ |
| H2AX Histone H2AX | P27661 | NO | NO | NO | YES | NO | NO | P/MQ |
| PDIA3 Protein disulfide-isomerase A3 | P27773 | YES | YES | YES | YES | YES | YES | P/MQ |
| VAV Proto-oncogene vav | P27870 | NO | NO | NO | YES | NO | NO | P |
| HNF1B Hepatocyte nuclear factor 1-beta | P27889 | NO | YES | NO | NO | NO | NO | P |
| PSB8 Proteasome subunit beta type-8 | P28063 | NO | NO | NO | YES | YES | YES | P |
| ACOC Cytoplasmic aconitate hydratase | P28271 | YES | YES | YES | YES | YES | YES | P/MQ |
| CTSG Cathepsin G | P28293 | NO | NO | YES | YES | YES | YES | P |
| APEX1 DNA-(apurinic or apyrimidinic site) lyase | P28352 | NO | YES | YES | YES | YES | YES | P/MQ |
| MAX Protein max | P28574 | NO | NO | NO | YES | NO | NO | P |
| PURA1 Adenylosuccinate synthetase isozyme 1 | P28650 | NO | NO | NO | YES | YES | YES | P |
| PGS1 Biglycan | P28653 | NO | YES | YES | YES | YES | YES | P/MQ |
| PGS2 Decorin | P28654 | NO | YES | NO | YES | NO | NO | P/MQ |
| NP1L1 Nucleosome assembly protein 1-like 1 | P28656 | YES | YES | YES | YES | YES | YES | P/MQ |
| MUG1 Murinoglobulin-1 | P28665 | YES | YES | NO | NO | YES | NO | P/MQ |
| MRP MARCKS-related protein | P28667 | NO | YES | YES | YES | YES | YES | P/MQ |
| MEP1A Meprin A subunit alpha | P28825 | NO | YES | NO | NO | NO | NO | P |
| DPP4 Dipeptidyl peptidase 4 | P28843 | YES | YES | NO | NO | NO | NO | P |
| KPCD Protein kinase C delta type | P28867 | NO | NO | NO | YES | NO | NO | P |
| PABP1 Polyadenylate-binding protein 1 | P29341 | YES | YES | YES | YES | YES | YES | P/MQ |
| PTN6 Tyrosine-protein phosphatase non-receptor type 6 | P29351 | YES | YES | YES | YES | YES | YES | P/MQ |
| FRIL1 Ferritin light chain 1 | P29391 | YES | YES | YES | YES | YES | YES | P/MQ |
| HEXA Beta-hexosaminidase subunit alpha | P29416 | NO | NO | NO | YES | YES | YES | P |
| NOS2 Nitric oxide synthase, inducible | P29477 | NO | NO | NO | NO | YES | NO | P |

| | | | | | | | | |
|---|---|---|---|---|---|---|---|---|
| VCAM1 Vascular cell adhesion protein 1 | P29533 | NO | YES | YES | YES | YES | NO | P/MQ |
| NEDD8 | P29595 | YES | YES | YES | YES | YES | YES | MQ |
| FETUA Alpha-2-HS-glycoprotein | P29699 | YES | YES | YES | YES | YES | YES | P/MQ |
| OAT Ornithine aminotransferase, mitochondrial | P29758 | YES | YES | YES | YES | YES | YES | P/MQ |
| VTNC Vitronectin | P29788 | NO | NO | YES | NO | YES | NO | P |
| KCRU Creatine kinase U-type, mitochondrial | P30275 | YES | YES | NO | NO | YES | NO | P |
| FKBP4 Peptidyl-prolyl cis-trans isomerase FKBP4 | P30416 | YES | YES | YES | YES | YES | YES | P/MQ |
| HMGB2 High mobility group protein B2 | P30681 | YES | NO | YES | YES | YES | YES | P/MQ |
| C5AR1 C5a anaphylatoxin chemotactic receptor 1 | P30993 | NO | NO | NO | YES | YES | YES | P |
| CTND1 Catenin delta-1 | P30999 | YES | YES | YES | YES | YES | YES | P/MQ |
| DESM Desmin | P31001 | YES | YES | YES | YES | YES | YES | P/MQ |
| AIMP1 Aminoacyl tRNA synthase complex-interacting multifunctional protein 1 | P31230 | NO | YES | NO | YES | YES | YES | P |
| DPEP1 Dipeptidase 1 | P31428 | YES | YES | NO | NO | NO | NO | P/MQ |
| S10A9 Protein S100-A9 | P31725 | YES | YES | YES | YES | YES | YES | P/MQ |
| ACBP Acyl-CoA-binding protein | P31786 | YES | YES | YES | YES | YES | YES | P/MQ |
| MP2K1 Dual specificity mitogen-activated protein kinase kinase 1 | P31938 | NO | YES | YES | YES | YES | YES | P/MQ |
| NLTP non-specific lipid-transfer protein | P32020 | YES | YES | YES | YES | YES | YES | P/MQ |
| LA Lupus La protein homolog | P32067 | YES | YES | YES | YES | YES | YES | P/MQ |
| ANT3 Antithrombin-III | P32261 | NO | YES | NO | NO | NO | NO | P/MQ |
| RASK GTPase KRas | P32883 | NO | YES | NO | NO | NO | NO | P |
| SYWC Tryptophan--tRNA ligase, cytoplasmic | P32921 | NO | NO | YES | YES | YES | NO | P/MQ |
| APOC3 Apolipoprotein C-III | P33622 | NO | YES | NO | NO | NO | NO | P |
| RANG Ran-specific GTPase-activating protein | P34022 | YES | YES | YES | YES | YES | YES | P/MQ |
| MIF Macrophage migration inhibitory factor | P34884 | YES | YES | YES | YES | YES | YES | P/MQ |
| H1 Bifunctional epoxide hydrolase 2 | P34914 | YES | YES | NO | NO | YES | NO | P |
| CYT2 Stefin-2 | P35174 | NO | NO | NO | YES | YES | YES | P |
| CYT1 Stefin-1 | P35175 | NO | NO | NO | YES | NO | NO | P |
| PTN11 Tyrosine-protein phosphatase non-receptor type 11 | P35235 | NO | YES | NO | NO | NO | NO | P |
| RAB5C Ras-related protein Rab-5C | P35278 | YES | YES | YES | YES | YES | YES | P/MQ |
| RAB6A Ras-related protein Rab-6A | P35279 | NO | NO | NO | YES | YES | NO | P/MQ |
| RAB21 Ras-related protein Rab-21 | P35282 | NO | NO | NO | YES | NO | YES | P |
| RAB18 Ras-related protein Rab-18 | P35293 | NO | YES | NO | NO | NO | NO | P |
| TSP1 Thrombospondin-1 | P35441 | NO | YES | NO | YES | NO | YES | P |
| ODPA Pyruvate dehydrogenase E1 component subunit alpha, somatic form, mitochondrial | P35486 | YES | YES | NO | YES | YES | YES | P/MQ |
| FAAA Fumarylacetoacetase | P35505 | YES | YES | YES | YES | YES | NO | P/MQ |
| FBRL rRNA 2'-O-methyltransferase fibrillarin | P35550 | YES | YES | YES | YES | YES | YES | P/MQ |
| CALX Calnexin | P35564 | YES | YES | NO | YES | YES | YES | P/MQ |

| | | | | | | | | |
|---|---|---|---|---|---|---|---|---|
| AP1M1 AP-1 complex subunit mu-1 | P35585 | NO | YES | YES | YES | YES | NO | P/MQ |
| PRDX1 Peroxiredoxin-1 | P35700 | YES | YES | YES | YES | YES | YES | P/MQ |
| PTN1 Tyrosine-protein phosphatase non-receptor type 1 | P35821 | NO | NO | NO | YES | NO | NO | P |
| PTN12 Tyrosine-protein phosphatase non-receptor type 12 | P35831 | NO | NO | NO | YES | NO | YES | P |
| RL12 60S ribosomal protein L12 | P35979 | YES | YES | YES | YES | YES | YES | P/MQ |
| RL18 60S ribosomal protein L18 | P35980 | YES | YES | YES | YES | YES | YES | P/MQ |
| TAP2 Antigen peptide transporter 2 | P36371 | NO | NO | NO | YES | NO | YES | P |
| PPM1B Protein phosphatase 1B | P36993 | NO | YES | NO | NO | YES | NO | P |
| NCPR NADPH--cytochrome P450 reductase | P37040 | YES | YES | YES | YES | YES | YES | P/MQ |
| TAGL Transgelin | P37804 | YES | YES | YES | YES | YES | YES | P/MQ |
| FBLN2 Fibulin-2 | P37889 | NO | NO | NO | YES | NO | YES | P |
| HMGCL Hydroxymethylglutaryl-CoA lyase, mitochondrial | P38060 | YES | YES | NO | NO | NO | NO | P |
| GRP75 Stress-70 protein, mitochondrial | P38647 | YES | YES | YES | YES | YES | YES | P/MQ |
| DYN2 Dynamin-2 | P39054 | YES | YES | YES | YES | YES | YES | P/MQ |
| COIA1 Collagen alpha-1(XVIII) chain | P39061 | YES | YES | YES | YES | YES | YES | P/MQ |
| MA1A2 Mannosyl-oligosaccharide 1,2-alpha-mannosidase IB | P39098 | NO | YES | NO | NO | NO | NO | P |
| ZO1 Tight junction protein ZO-1 | P39447 | YES | YES | NO | YES | YES | YES | P/MQ |
| CAP1 Adenylyl cyclase-associated protein 1 | P40124 | YES | YES | YES | YES | YES | YES | P/MQ |
| CHD1 Chromodomain-helicase-DNA-binding protein 1 | P40201 | NO | YES | NO | NO | NO | NO | P |
| VP26A Vacuolar protein sorting-associated protein 26A | P40336 | YES | YES | NO | NO | YES | NO | P |
| TFAM Transcription factor A, mitochondrial | P40630 | YES | NO | NO | NO | YES | NO | P |
| RL28 60S ribosomal protein L28 | P41105 | NO | YES | YES | YES | YES | YES | P/MQ |
| ACSL1 Long-chain-fatty-acid--CoA ligase 1 | P41216 | YES | YES | NO | NO | YES | NO | P/MQ |
| CSK Tyrosine-protein kinase CSK | P41241 | NO | NO | NO | YES | NO | NO | P |
| MMP9 Matrix metalloproteinase-9 | P41245 | NO | NO | YES | YES | NO | YES | P/MQ |
| SEPT2 Septin-2 | P42208 | YES | YES | YES | YES | YES | YES | P/MQ |
| STAT1 Signal transducer and activator of transcription 1 | P42225 | NO | YES | YES | YES | YES | YES | P/MQ |
| STAT3 Signal transducer and activator of transcription 3 | P42227 | NO | NO | NO | YES | NO | YES | P |
| EPS15 Epidermal growth factor receptor substrate 15 | P42567 | NO | YES | YES | YES | YES | YES | P/MQ |
| TCPQ T-complex protein 1 subunit theta | P42932 | YES | YES | YES | YES | YES | YES | P/MQ |
| H14 Histone H1.4 | P43274 | YES | YES | YES | YES | YES | YES | P/MQ |
| H11 Histone H1.1 | P43275 | YES | YES | YES | YES | YES | YES | P/MQ |
| H15 Histone H1.5 | P43276 | YES | YES | YES | YES | YES | YES | P/MQ |

| | | | | | | | | |
|---|---|---|---|---|---|---|---|---|
| H13 Histone H1.3 | P43277 | YES | YES | YES | YES | YES | YES | P/MQ |
| ITAV Integrin alpha-V | P43406 | NO | YES | NO | NO | NO | NO | P |
| PLIN2 Perilipin-2 | P43883 | NO | NO | NO | YES | YES | YES | P |
| ALDR Aldose reductase | P45376 | YES | YES | YES | YES | YES | YES | P/MQ |
| COF2 Cofilin-2 | P45591 | NO | YES | NO | NO | NO | NO | P |
| FKBP2 Peptidyl-prolyl cis-trans isomerase FKBP2 | P45878 | NO | YES | NO | NO | NO | NO | P |
| CDN1B Cyclin-dependent kinase inhibitor 1B | P46414 | NO | YES | NO | YES | YES | YES | P/MQ |
| NSF Vesicle-fusing ATPase | P46460 | YES | YES | YES | YES | YES | YES | P/MQ |
| VPS4B Vacuolar protein sorting-associated protein 4B | P46467 | NO | NO | NO | YES | YES | YES | P |
| PRS7 26S proteasome regulatory subunit 7 | P46471 | YES | YES | YES | YES | YES | YES | P/MQ |
| RB11B Ras-related protein Rab-11B | P46638 | YES | NO | NO | YES | YES | NO | P/MQ |
| ADX Adrenodoxin, mitochondrial | P46656 | NO | YES | NO | NO | NO | NO | P |
| PURA2 Adenylosuccinate synthetase isozyme 2 | P46664 | NO | NO | NO | YES | YES | NO | P |
| YAP1 Transcriptional coactivator YAP1 | P46938 | NO | YES | NO | YES | NO | NO | P |
| QOR Quinone oxidoreductase | P47199 | YES | YES | YES | YES | YES | YES | P/MQ |
| TES Testin | P47226 | NO | NO | NO | YES | NO | NO | P |
| PA24A Cytosolic phospholipase A2 | P47713 | NO | NO | NO | YES | NO | NO | P |
| ALDH2 Aldehyde dehydrogenase, mitochondrial | P47738 | YES | YES | YES | YES | YES | YES | P/MQ |
| AL3A2 Fatty aldehyde dehydrogenase | P47740 | YES | YES | NO | YES | NO | YES | P/MQ |
| CAZA1 F-actin-capping protein subunit alpha-1 | P47753 | NO | YES | YES | YES | YES | YES | P/MQ |
| CAZA2 F-actin-capping protein subunit alpha-2 | P47754 | YES | YES | YES | YES | YES | YES | P/MQ |
| CAPZB F-actin-capping protein subunit beta | P47757 | YES | YES | YES | YES | YES | YES | P/MQ |
| SRPRB Signal recognition particle receptor subunit beta | P47758 | NO | YES | NO | YES | NO | YES | P/MQ |
| RL6 60S ribosomal protein L6 | P47911 | YES | YES | YES | YES | YES | YES | P/MQ |
| RL29 60S ribosomal protein L29 | P47915 | YES | NO | YES | YES | YES | YES | P |
| CRAT Carnitine O-acetyltransferase | P47934 | YES | YES | NO | NO | NO | NO | P |
| CRKL Crk-like protein | P47941 | NO | YES | NO | YES | NO | NO | P/MQ |
| RLA1 60S acidic ribosomal protein P1 | P47955 | NO | NO | NO | YES | NO | NO | P |
| RL5 60S ribosomal protein L5 | P47962 | YES | YES | YES | YES | YES | YES | P/MQ |
| RL13 60S ribosomal protein L13 | P47963 | YES | YES | YES | YES | YES | YES | P/MQ |
| RL36 60S ribosomal protein L36 | P47964 | NO | YES | NO | NO | NO | NO | P |
| EIF1 Eukaryotic translation initiation factor 1 | P48024 | NO | NO | NO | YES | YES | NO | P/MQ |
| KSYK Tyrosine-protein kinase SYK | P48025 | NO | NO | NO | YES | YES | YES | P |
| ANXA5 Annexin A5 | P48036 | YES | YES | YES | YES | YES | YES | P/MQ |
| 41 Protein 4.1 | P48193 | YES | YES | YES | YES | YES | NO | P/MQ |
| TBCA Tubulin-specific chaperone A | P48428 | NO | YES | NO | YES | YES | YES | P |
| LMNA Prelamin-A/C | P48678 | YES | YES | YES | YES | YES | YES | P/MQ |
| HS74L Heat shock 70 kDa protein 4L | P48722 | YES | NO | NO | NO | YES | NO | P |
| CBR1 Carbonyl reductase [NADPH] 1 | P48758 | YES | YES | YES | YES | YES | YES | P/MQ |

| | | | | | | | | |
|---|---|---|---|---|---|---|---|---|
| CX7A2 Cytochrome c oxidase subunit 7A2, mitochondrial | P48771 | NO | YES | NO | NO | NO | NO | P |
| ADT1 ADP/ATP translocase 1 | P48962 | NO | YES | NO | NO | NO | NO | P |
| MAPK2 MAP kinase-activated protein kinase 2 | P49138 | NO | NO | NO | YES | NO | NO | P |
| HEP2 Heparin cofactor 2 | P49182 | NO | YES | NO | NO | NO | NO | P |
| ROA1 Heterogeneous nuclear ribonucleoprotein A1 | P49312 | YES | YES | YES | YES | YES | YES | P/MQ |
| INPP Inositol polyphosphate 1-phosphatase | P49442 | NO | NO | NO | YES | YES | NO | P |
| PPM1A Protein phosphatase 1A | P49443 | NO | NO | NO | YES | NO | NO | P |
| PCY1A Choline-phosphate cytidylyltransferase A | P49586 | NO | YES | NO | NO | YES | NO | P/MQ |
| HCLS1 Hematopoietic lineage cell-specific protein | P49710 | YES | YES | YES | YES | YES | YES | P/MQ |
| MCM5 DNA replication licensing factor MCM5 | P49718 | NO | NO | NO | YES | NO | NO | P |
| PSA2 Proteasome subunit alpha type-2 | P49722 | YES | YES | YES | YES | YES | YES | P/MQ |
| ODBA 2-oxoisovalerate dehydrogenase subunit alpha, mitochondrial | P50136 | NO | YES | NO | NO | NO | NO | P |
| DHB8 Estradiol 17-beta-dehydrogenase 8 | P50171 | YES | YES | NO | NO | NO | NO | P |
| SAHH Adenosylhomocysteinase | P50247 | YES | YES | YES | YES | YES | YES | P/MQ |
| FMO1 Dimethylaniline monooxygenase [N-oxide-forming] 1 | P50285 | NO | YES | NO | NO | NO | NO | P |
| GDIA Rab GDP dissociation inhibitor alpha | P50396 | NO | NO | NO | NO | YES | NO | P |
| ARSB Arylsulfatase B | P50429 | NO | YES | NO | NO | NO | NO | P |
| GLYC Serine hydroxymethyltransferase, cytosolic | P50431 | NO | YES | NO | NO | NO | NO | P |
| VATA V-type proton ATPase catalytic subunit A | P50516 | YES | YES | YES | YES | YES | YES | P/MQ |
| VATE1 V-type proton ATPase subunit E 1 | P50518 | YES | YES | YES | YES | YES | YES | P/MQ |
| S10AB Protein S100-A11 | P50543 | NO | YES | NO | YES | YES | YES | P |
| PA2G4 Proliferation-associated protein 2G4 | P50580 | YES | YES | YES | YES | YES | YES | P/MQ |
| ICAL Calpastatin | P51125 | NO | YES | YES | YES | YES | YES | P/MQ |
| RAB7A Ras-related protein Rab-7a | P51150 | YES | YES | YES | YES | YES | YES | P/MQ |
| RL9 60S ribosomal protein L9 | P51410 | YES | YES | YES | YES | YES | NO | P |
| CAMP Cathelicidin antimicrobial peptide | P51437 | NO | NO | NO | YES | NO | NO | P |
| DHB4 Peroxisomal multifunctional enzyme type 2 | P51660 | YES | YES | YES | YES | YES | YES | P/MQ |
| DYLT1 Dynein light chain Tctex-type 1 | P51807 | NO | NO | NO | YES | NO | NO | P |
| HDGF Hepatoma-derived growth factor | P51859 | YES | YES | YES | YES | YES | YES | P/MQ |
| VA0D1 V-type proton ATPase subunit d 1 | P51863 | YES | NO | NO | YES | YES | NO | P |
| FABP7 Fatty acid-binding protein, brain | P51880 | NO | YES | NO | NO | NO | NO | P |
| ADT2 ADP/ATP translocase 2 | P51881 | YES | YES | YES | YES | YES | YES | P/MQ |
| LUM Lumican | P51885 | NO | YES | NO | YES | YES | YES | P/MQ |
| THTR Thiosulfate sulfurtransferase | P52196 | YES | YES | NO | NO | NO | NO | P |
| NDUS6 NADH dehydrogenase [ubiquinone] iron-sulfur protein 6, mitochondrial | P52503 | YES | YES | YES | NO | NO | NO | P |
| UCK1 Uridine-cytidine kinase 1 | P52623 | NO | YES | NO | NO | NO | NO | P |
| RIDA 2-iminobutanoate/2-iminopropanoate deaminase | P52760 | YES | YES | YES | YES | YES | NO | P/MQ |

| | | | | | | | | |
|---|---|---|---|---|---|---|---|---|
| EFNB1 Ephrin-B1 | P52795 | NO | YES | NO | NO | NO | NO | P |
| CPT2 Carnitine O-palmitoyltransferase 2, mitochondrial | P52825 | YES | YES | NO | NO | NO | NO | P |
| RL10A 60S ribosomal protein L10a | P53026 | YES | YES | NO | YES | YES | YES | P/MQ |
| ODB2 Lipoamide acyltransferase component of branched-chain alpha-keto acid dehydrogenase complex, mitochondrial | P53395 | YES | YES | NO | NO | NO | NO | P |
| KPYR Pyruvate kinase PKLR | P53657 | YES | YES | NO | NO | NO | NO | P |
| PIPNB Phosphatidylinositol transfer protein beta isoform | P53811 | NO | YES | NO | NO | NO | NO | P |
| MOT1 Monocarboxylate transporter 1 | P53986 | NO | YES | NO | NO | NO | NO | P/MQ |
| RAB2A Ras-related protein Rab-2A | P53994 | YES | YES | NO | YES | YES | YES | P/MQ |
| STOM Erythrocyte band 7 integral membrane protein | P54116 | NO | NO | NO | NO | YES | NO | P/MQ |
| STMN1 Stathmin | P54227 | YES | YES | YES | YES | YES | YES | P/MQ |
| RD23A UV excision repair protein RAD23 homolog A | P54726 | NO | YES | NO | NO | NO | NO | P |
| RD23B UV excision repair protein RAD23 homolog B | P54728 | NO | YES | NO | YES | NO | YES | P/MQ |
| NUB1 NEDD8 ultimate buster 1 | P54729 | NO | YES | NO | NO | NO | NO | P |
| FAF1 FAS-associated factor 1 | P54731 | NO | YES | NO | NO | NO | NO | P |
| PRS6B 26S proteasome regulatory subunit 6B | P54775 | NO | YES | NO | YES | YES | YES | P/MQ |
| PUR8 Adenylosuccinate lyase | P54822 | NO | YES | NO | YES | NO | NO | P |
| DDX6 Probable ATP-dependent RNA helicase DDX6 | P54823 | NO | YES | YES | YES | YES | YES | P/MQ |
| ADPRH [Protein ADP-ribosylarginine] hydrolase | P54923 | NO | NO | NO | YES | NO | NO | P |
| IRG1 Cis-aconitate decarboxylase | P54987 | NO | NO | NO | YES | NO | NO | P |
| S12A1 Solute carrier family 12 member 1 | P55014 | NO | YES | NO | NO | NO | NO | P |
| 3BP1 SH3 domain-binding protein 1 | P55194 | NO | NO | NO | YES | YES | YES | P |
| RAB8A Ras-related protein Rab-8A | P55258 | NO | YES | NO | YES | NO | YES | P |
| Adenosine kinase | P55264 | YES | YES | YES | YES | YES | YES | MQ |
| AMRP Alpha-2-macroglobulin receptor-associated protein | P55302 | YES | YES | YES | YES | NO | YES | P |
| Stathmin-2 | P55821 | YES | YES | YES | YES | YES | YES | MQ |
| GOGA3 Golgin subfamily A member 3 | P55937 | NO | NO | NO | YES | YES | NO | P |
| ATPK ATP synthase subunit f, mitochondrial | P56135 | YES | YES | YES | YES | YES | YES | P/MQ |
| ACYP2 Acylphosphatase-2 | P56375 | NO | YES | NO | NO | NO | NO | P |
| ATP5E ATP synthase subunit epsilon, mitochondrial | P56382 | NO | YES | NO | NO | NO | NO | P |
| CDD Cytidine deaminase | P56389 | NO | YES | NO | NO | NO | NO | P/MQ |
| CX6B1 Cytochrome c oxidase subunit 6B1 | P56391 | YES | YES | NO | NO | NO | NO | P/MQ |
| COX17 Cytochrome c oxidase copper chaperone | P56394 | NO | NO | NO | YES | NO | YES | P |
| CYB5 Cytochrome b5 | P56395 | YES | YES | YES | YES | YES | YES | P/MQ |
| UBP5 Ubiquitin carboxyl-terminal hydrolase 5 | P56399 | YES | YES | YES | YES | YES | YES | P/MQ |
| ATP synthase subunit beta, mitochondrial | P56480 | YES | YES | YES | YES | YES | YES | MQ |
| PDCD5 Programmed cell death protein 5 | P56812 | YES | YES | YES | YES | YES | YES | P/MQ |
| FUS RNA-binding protein FUS | P56959 | YES | YES | NO | YES | YES | YES | P/MQ |

| | | | | | | | | |
|---|---|---|---|---|---|---|---|---|
| LAD1 Ladinin-1 | P57016 | YES | YES | YES | YES | YES | YES | P/MQ |
| PCBP3 Poly(rC)-binding protein 3 | P57722 | NO | YES | NO | NO | NO | NO | P |
| VATD V-type proton ATPase subunit D | P57746 | YES | YES | NO | NO | YES | NO | P |
| ERP29 Endoplasmic reticulum resident protein 29 | P57759 | YES | YES | YES | YES | YES | YES | P/MQ |
| EF1D Elongation factor 1-delta | P57776 | YES | YES | YES | YES | YES | YES | P/MQ |
| ACTN4 Alpha-actinin-4 | P57780 | YES | YES | NO | NO | NO | NO | P/MQ |
| RU2A U2 small nuclear ribonucleoprotein A' | P57784 | YES | YES | YES | YES | YES | YES | P/MQ |
| RT21 28S ribosomal protein S21, mitochondrial | P58059 | NO | YES | NO | NO | NO | NO | P |
| EF2 Elongation factor 2 | P58252 | YES | YES | YES | YES | YES | YES | P/MQ |
| OPA1 Dynamin-like 120 kDa protein, mitochondrial | P58281 | NO | YES | NO | NO | NO | NO | P |
| PTPA Serine/threonine-protein phosphatase 2A activator | P58389 | NO | YES | NO | YES | NO | NO | P |
| STRN4 Striatin-4 | P58404 | NO | YES | NO | NO | NO | NO | P |
| TPM1 Tropomyosin alpha-1 chain | P58771 | YES | YES | YES | YES | YES | YES | P |
| TPM2 Tropomyosin beta chain | P58774 | YES | YES | YES | YES | YES | YES | P |
| TB182 182 kDa tankyrase-1-binding protein | P58871 | YES | YES | YES | YES | YES | YES | P/MQ |
| B2L13 Bcl-2-like protein 13 | P59017 | NO | YES | NO | NO | NO | NO | P |
| S12A3 Solute carrier family 12 member 3 | P59158 | NO | YES | NO | NO | NO | NO | P |
| CING Cingulin | P59242 | NO | YES | NO | NO | NO | NO | P |
| IF5 Eukaryotic translation initiation factor 5 | P59325 | NO | NO | NO | YES | NO | YES | P |
| ANS1A Ankyrin repeat and SAM domain-containing protein 1A | P59672 | NO | YES | NO | NO | NO | NO | P |
| ARPC4 Actin-related protein 2/3 complex subunit 4 | P59999 | YES | YES | YES | YES | YES | YES | P/MQ |
| RUVB1 RuvB-like 1 | P60122 | NO | NO | YES | YES | YES | YES | P/MQ |
| EIF3E Eukaryotic translation initiation factor 3 subunit E | P60229 | NO | YES | NO | NO | NO | NO | P |
| PCBP1 Poly(rC)-binding protein 1 | P60335 | YES | YES | YES | YES | YES | YES | P/MQ |
| ACTB Actin, cytoplasmic 1 | P60710 | YES | YES | YES | YES | YES | YES | P |
| CDC42 Cell division control protein 42 homolog | P60766 | YES | YES | NO | YES | YES | YES | P/MQ |
| CIRBP Cold-inducible RNA-binding protein | P60824 | NO | YES | NO | NO | NO | NO | P |
| IF4A1 Eukaryotic initiation factor 4A-I | P60843 | YES | YES | YES | YES | YES | YES | P/MQ |
| RS20 40S ribosomal protein S20 | P60867 | YES | YES | NO | YES | YES | YES | P/MQ |
| RAB5B Ras-related protein Rab-5B | P61021 | YES | YES | YES | YES | YES | YES | P/MQ |
| CHP1 Calcineurin B homologous protein 1 | P61022 | NO | YES | NO | YES | NO | NO | P |
| RAB10 Ras-related protein Rab-10 | P61027 | YES | YES | YES | YES | YES | YES | P/MQ |
| RAB8B Ras-related protein Rab-8B | P61028 | NO | YES | NO | YES | YES | YES | P/MQ |
| UBC12 NEDD8-conjugating enzyme Ubc12 | P61082 | NO | YES | NO | NO | NO | NO | P |
| UBE2K Ubiquitin-conjugating enzyme E2 K | P61087 | NO | YES | NO | NO | YES | YES | P |
| UBE2N Ubiquitin-conjugating enzyme E2 N | P61089 | YES | YES | YES | YES | YES | YES | P/MQ |
| ARP2 Actin-related protein 2 | P61161 | YES | NO | YES | YES | YES | YES | P/MQ |
| ACTZ Alpha-centractin | P61164 | NO | YES | NO | YES | YES | NO | P |
| ABCE1 ATP-binding cassette sub-family E member 1 | P61222 | NO | NO | NO | NO | YES | NO | P |

| | | | | | | | | |
|---|---|---|---|---|---|---|---|---|
| RL26 60S ribosomal protein L26 | P61255 | YES | YES | YES | YES | YES | YES | P/MQ |
| PSME3 Proteasome activator complex subunit 3 | P61290 | NO | NO | NO | YES | YES | YES | P/MQ |
| RAB6B Ras-related protein Rab-6B | P61294 | YES | YES | NO | NO | NO | NO | P |
| RL27 60S ribosomal protein L27 | P61358 | YES | YES | YES | YES | YES | YES | P/MQ |
| PHS Pterin-4-alpha-carbinolamine dehydratase | P61458 | NO | YES | NO | NO | NO | NO | P |
| ARF4 ADP-ribosylation factor 4 | P61750 | NO | YES | NO | NO | NO | NO | P/MQ |
| GABT 4-aminobutyrate aminotransferase, mitochondrial | P61922 | NO | YES | NO | NO | NO | NO | P |
| COPZ1 Coatomer subunit zeta-1 | P61924 | NO | NO | NO | YES | YES | NO | P/MQ |
| SUMO2 Small ubiquitin-related modifier 2 | P61957 | YES | YES | NO | YES | NO | NO | P/MQ |
| UFM1 Ubiquitin-fold modifier 1 | P61961 | NO | YES | NO | YES | NO | NO | P/MQ |
| AP1S1 AP-1 complex subunit sigma-1A | P61967 | NO | YES | NO | NO | NO | NO | P/MQ |
| HNRPK Heterogeneous nuclear ribonucleoprotein K | P61979 | YES | YES | YES | YES | YES | YES | P/MQ |
| 1433G 14-3-3 protein gamma | P61982 | YES | YES | YES | YES | YES | YES | P/MQ |
| TIM13 Mitochondrial import inner membrane translocase subunit Tim13 | P62075 | YES | YES | NO | NO | YES | NO | P |
| Mitochondrial import inner membrane translocase subunit Tim8 B | P62077 | YES | YES | YES | YES | YES | YES | MQ |
| RS7 40S ribosomal protein S7 | P62082 | NO | YES | NO | YES | YES | NO | P |
| PP1A Serine/threonine-protein phosphatase PP1-alpha catalytic subunit | P62137 | YES | YES | YES | YES | YES | YES | P/MQ |
| PRS4 26S proteasome regulatory subunit 4 | P62192 | YES | YES | NO | YES | YES | YES | P/MQ |
| PRS8 26S proteasome regulatory subunit 8 | P62196 | NO | YES | YES | YES | YES | YES | P/MQ |
| RS8 40S ribosomal protein S8 | P62242 | YES | YES | YES | YES | YES | YES | P/MQ |
| RS15A 40S ribosomal protein S15a | P62245 | YES | YES | NO | YES | YES | YES | P/MQ |
| 1433E 14-3-3 protein epsilon | P62259 | YES | YES | YES | YES | YES | YES | P/MQ |
| RS14 40S ribosomal protein S14 | P62264 | YES | YES | YES | YES | YES | YES | P/MQ |
| RS23 40S ribosomal protein S23 | P62267 | YES | YES | YES | YES | YES | YES | P/MQ |
| RS18 40S ribosomal protein S18 | P62270 | YES | YES | YES | YES | YES | YES | P/MQ |
| RS29 40S ribosomal protein S29 | P62274 | NO | NO | NO | YES | NO | NO | P |
| RS11 40S ribosomal protein S11 | P62281 | NO | YES | YES | YES | YES | YES | P/MQ |
| RS13 40S ribosomal protein S13 | P62301 | YES | YES | YES | YES | YES | YES | P/MQ |
| LSM3 U6 snRNA-associated Sm-like protein LSm3 | P62311 | NO | YES | NO | NO | NO | YES | P |
| LSM6 U6 snRNA-associated Sm-like protein LSm6 | P62313 | NO | YES | NO | NO | NO | NO | P |
| SMD2 Small nuclear ribonucleoprotein Sm D2 | P62317 | YES | YES | YES | YES | YES | YES | P/MQ |
| PRS10 26S proteasome regulatory subunit 10B | P62334 | YES | YES | YES | YES | YES | YES | P/MQ |
| RB11A Ras-related protein Rab-11A | P62492 | NO | YES | NO | NO | NO | NO | P |
| RS4X 40S ribosomal protein S4, X isoform | P62500 | YES | YES | NO | NO | NO | NO | P/MQ |
| DLRB1 Dynein light chain roadblock-type 1 | P62627 | NO | YES | NO | NO | NO | NO | P/MQ |
| RS4X 40S ribosomal protein S4, X isoform | P62702 | YES | YES | YES | YES | YES | YES | P/MQ |
| RL18A 60S ribosomal protein L18a | P62717 | NO | NO | NO | YES | YES | NO | P |
| ACTA Actin, aortic smooth muscle | P62737 | NO | YES | NO | NO | NO | NO | P/MQ |

| | | | | | | | | |
|---|---|---|---|---|---|---|---|---|
| HPCL1 Hippocalcin-like protein 1 | P62748 | NO | NO | NO | NO | YES | NO | P |
| RL23A 60S ribosomal protein L23a | P62751 | YES | YES | YES | YES | YES | YES | P/MQ |
| RS6 40S ribosomal protein S6 | P62754 | YES | YES | YES | YES | YES | YES | P/MQ |
| MTPN Myotrophin | P62774 | YES | YES | NO | YES | YES | YES | P/MQ |
| H4 Histone H4 | P62806 | YES | YES | YES | YES | YES | YES | P/MQ |
| VATB2 V-type proton ATPase subunit B, brain isoform | P62814 | YES | YES | YES | YES | YES | YES | P/MQ |
| RAB1A Ras-related protein Rab-1A | P62821 | YES | YES | YES | YES | YES | YES | P/MQ |
| RAN GTP-binding nuclear protein Ran | P62827 | YES | YES | YES | YES | YES | YES | P/MQ |
| RL23 60S ribosomal protein L23 | P62830 | NO | YES | YES | YES | YES | YES | P |
| RS15 40S ribosomal protein S15 | P62843 | NO | YES | YES | YES | NO | NO | P |
| RS24 40S ribosomal protein S24 | P62849 | NO | NO | YES | NO | NO | NO | P |
| RS25 40S ribosomal protein S25 | P62852 | YES | YES | YES | YES | NO | YES | P/MQ |
| RS26 40S ribosomal protein S26 | P62855 | NO | YES | NO | YES | NO | YES | P |
| RS28 40S ribosomal protein S28 | P62858 | YES | YES | YES | YES | YES | YES | P/MQ |
| RS30 40S ribosomal protein S30 | P62862 | YES | YES | YES | YES | YES | YES | P/MQ |
| ELOB Elongin-B | P62869 | NO | YES | NO | YES | NO | YES | P |
| GBB1 Guanine nucleotide-binding protein G(I)/G(S)/G(T) subunit beta-1 | P62874 | NO | YES | NO | NO | NO | NO | P/MQ |
| GBB2 Guanine nucleotide-binding protein G(I)/G(S)/G(T) subunit beta-2 | P62880 | NO | YES | NO | NO | NO | NO | P/MQ |
| RL30 60S ribosomal protein L30 | P62889 | NO | YES | YES | YES | NO | YES | P |
| RL39 60S ribosomal protein L39 | P62892 | NO | NO | NO | YES | YES | NO | P |
| CYC Cytochrome c, somatic | P62897 | YES | YES | YES | YES | YES | YES | P/MQ |
| RL31 60S ribosomal protein L31 | P62900 | YES | YES | YES | YES | YES | YES | P/MQ |
| RS3 40S ribosomal protein S3 | P62908 | YES | YES | YES | YES | YES | YES | P/MQ |
| RL32 60S ribosomal protein L32 | P62911 | NO | NO | YES | YES | NO | NO | P |
| RL8 60S ribosomal protein L8 | P62918 | YES | YES | YES | YES | YES | YES | P/MQ |
| YBOX1 Nuclease-sensitive element-binding protein 1 | P62960 | YES | YES | YES | YES | YES | YES | P/MQ |
| PROF1 Profilin-1 | P62962 | YES | YES | YES | YES | YES | YES | P/MQ |
| Ubiquitin-40S ribosomal protein S27a | P62983 | YES | YES | YES | YES | YES | YES | MQ |
| TRA2B Transformer-2 protein homolog beta | P62996 | NO | YES | NO | YES | YES | YES | P |
| RAC1 Ras-related C3 botulinum toxin substrate 1 | P63001 | YES | YES | NO | YES | YES | YES | P |
| LIS1 Platelet-activating factor acetylhydrolase IB subunit alpha | P63005 | YES | YES | NO | YES | YES | YES | P |
| HSP7C Heat shock cognate 71 kDa protein | P63017 | YES | YES | YES | YES | YES | YES | P/MQ |
| VAMP3 Vesicle-associated membrane protein 3 | P63024 | NO | YES | YES | NO | YES | NO | P/MQ |
| TCTP Translationally-controlled tumor protein | P63028 | NO | NO | YES | YES | YES | YES | P |
| CH60 60 kDa heat shock protein, mitochondrial | P63038 | YES | YES | YES | YES | YES | YES | P/MQ |
| MAPK1 Mitogen-activated protein kinase 1 | P63085 | NO | YES | NO | NO | NO | NO | P |
| PP1G Serine/threonine-protein phosphatase PP1-gamma catalytic subunit | P63087 | NO | YES | YES | YES | YES | YES | P/MQ |
| 1433Z 14-3-3 protein zeta/delta | P63101 | YES | YES | YES | YES | YES | YES | P/MQ |

| | | | | | | | | |
|---|---|---|---|---|---|---|---|---|
| HMGB1 High mobility group protein B1 | P63158 | YES | YES | YES | YES | YES | YES | P/MQ |
| Small ubiquitin-related modifier 1 | P63166 | NO | YES | YES | YES | NO | YES | MQ |
| DYL1 Dynein light chain 1, cytoplasmic | P63168 | NO | NO | NO | YES | YES | NO | P |
| IF5A1 Eukaryotic translation initiation factor 5A-1 | P63242 | YES | YES | YES | YES | YES | YES | P/MQ |
| Actin, cytoplasmic 2 | P63260 | YES | YES | YES | YES | YES | YES | MQ |
| RALA Ras-related protein Ral-A | P63321 | NO | YES | NO | NO | NO | NO | P |
| RS12 40S ribosomal protein S12 | P63323 | NO | YES | YES | YES | NO | YES | P |
| RS10 40S ribosomal protein S10 | P63325 | NO | YES | YES | YES | YES | YES | P/MQ |
| PP2AA Serine/threonine-protein phosphatase 2A catalytic subunit alpha isoform | P63330 | NO | YES | NO | YES | YES | YES | P |
| PHB Prohibitin | P67778 | YES | YES | YES | YES | YES | YES | P/MQ |
| RL22 60S ribosomal protein L22 | P67984 | YES | YES | YES | YES | YES | YES | P/MQ |
| ACTC Actin, alpha cardiac muscle 1 | P68033 | NO | YES | YES | YES | YES | NO | P/MQ |
| UB2L3 Ubiquitin-conjugating enzyme E2 L3 | P68037 | YES | YES | YES | YES | YES | YES | P |
| RACK1 Receptor of activated protein C kinase 1 | P68040 | YES | YES | YES | YES | YES | YES | P/MQ |
| 1433T 14-3-3 protein theta | P68254 | YES | YES | YES | YES | YES | YES | P/MQ |
| TBA4A Tubulin alpha-4A chain | P68368 | YES | YES | YES | YES | YES | YES | P/MQ |
| TBB4B Tubulin beta-4B chain | P68372 | YES | YES | YES | YES | YES | YES | P/MQ |
| TBA1C Tubulin alpha-1C chain | P68373 | YES | YES | YES | YES | YES | YES | P/MQ |
| 1433F 14-3-3 protein eta | P68510 | YES | YES | YES | YES | YES | YES | P/MQ |
| IMB1 Importin subunit beta-1 | P70168 | NO | YES | NO | YES | YES | NO | P |
| PSB7 Proteasome subunit beta type-7 | P70195 | NO | YES | NO | NO | NO | NO | P |
| MYO1F Unconventional myosin-If | P70248 | NO | NO | NO | YES | NO | YES | P/MQ |
| PKN1 Serine/threonine-protein kinase N1 | P70268 | NO | YES | NO | YES | NO | NO | P |
| PDLI4 PDZ and LIM domain protein 4 | P70271 | NO | NO | NO | YES | NO | NO | P |
| EM55 55 kDa erythrocyte membrane protein | P70290 | YES | YES | YES | NO | YES | NO | P |
| PEBP1 Phosphatidylethanolamine-binding protein 1 | P70296 | YES | YES | YES | YES | YES | YES | P/MQ |
| STIM1 Stromal interaction molecule 1 | P70302 | NO | NO | NO | YES | NO | NO | P |
| WASP Wiskott-Aldrich syndrome protein homolog | P70315 | NO | YES | YES | YES | YES | YES | P |
| TIAR Nucleolysin TIAR | P70318 | NO | YES | NO | NO | NO | NO | P |
| HNRH2 Heterogeneous nuclear ribonucleoprotein H2 | P70333 | NO | YES | NO | NO | NO | NO | P |
| ROCK1 Rho-associated protein kinase 1 | P70335 | NO | NO | NO | YES | NO | NO | P |
| FRAT1 Proto-oncogene FRAT1 | P70339 | NO | YES | NO | NO | NO | NO | P |
| HINT1 Histidine triad nucleotide-binding protein 1 | P70349 | YES | YES | YES | YES | YES | YES | P/MQ |
| ELAV1 ELAV-like protein 1 | P70372 | YES | NO | YES | YES | YES | YES | P/MQ |
| TP53B TP53-binding protein 1 | P70399 | NO | NO | NO | YES | NO | NO | P |
| MYBPH Myosin-binding protein H | P70402 | NO | YES | NO | NO | NO | NO | P |
| NHRF1 Na(+)/H(+) exchange regulatory cofactor NHE-RF1 | P70441 | YES | YES | YES | YES | YES | YES | P/MQ |
| BID BH3-interacting domain death agonist | P70444 | NO | YES | NO | NO | NO | NO | P |
| STX4 Syntaxin-4 | P70452 | NO | NO | NO | YES | NO | NO | P |
| VASP Vasodilator-stimulated phosphoprotein | P70460 | NO | YES | YES | YES | YES | YES | P/MQ |

| | | | | | | | | |
|---|---|---|---|---|---|---|---|---|
| NACAM Nascent polypeptide-associated complex subunit alpha, muscle-specific form | P70670 | NO | YES | NO | YES | NO | NO | P/MQ |
| CASP3 Caspase-3 | P70677 | NO | YES | NO | NO | NO | NO | P |
| UD12 UDP-glucuronosyltransferase 1-2 | P70691 | NO | YES | NO | NO | NO | NO | P |
| GAA Lysosomal alpha-glucosidase | P70699 | YES | YES | NO | YES | NO | YES | P |
| TCPH T-complex protein 1 subunit eta | P80313 | YES | YES | YES | YES | YES | YES | P/MQ |
| TCPB T-complex protein 1 subunit beta | P80314 | YES | YES | YES | YES | YES | YES | P/MQ |
| TCPD T-complex protein 1 subunit delta | P80315 | YES | YES | YES | YES | YES | YES | P/MQ |
| TCPE T-complex protein 1 subunit epsilon | P80316 | YES | YES | YES | YES | YES | YES | P/MQ |
| TCPG T-complex protein 1 subunit gamma | P80317 | YES | YES | NO | NO | NO | NO | P/MQ |
| TCPG T-complex protein 1 subunit gamma | P80318 | YES | NO | YES | YES | YES | YES | P/MQ |
| NUCB2 Nucleobindin-2 | P81117 | NO | NO | NO | YES | NO | YES | P/MQ |
| BGH3 Transforming growth factor-beta-induced protein ig-h3 | P82198 | NO | NO | YES | YES | YES | YES | P/MQ |
| RENBP N-acylglucosamine 2-epimerase | P82343 | NO | YES | NO | NO | NO | NO | P |
| WNK1 Serine/threonine-protein kinase WNK1 | P83741 | NO | NO | NO | YES | NO | YES | P/MQ |
| RL36A 60S ribosomal protein L36a | P83882 | YES | NO | NO | NO | YES | YES | P |
| CBX1 Chromobox protein homolog 1 | P83917 | NO | NO | NO | YES | NO | YES | P |
| ELOC Elongin-C | P83940 | NO | NO | NO | YES | NO | NO | P/MQ |
| ARF1 ADP-ribosylation factor 1 | P84078 | YES | NO | YES | YES | YES | YES | P/MQ |
| ARF5 ADP-ribosylation factor 5 | P84084 | NO | YES | NO | NO | NO | NO | P/MQ |
| Enhancer of rudimentary homolog | P84089 | YES | YES | YES | YES | YES | YES | MQ |
| AP2M1 AP-2 complex subunit mu | P84091 | NO | NO | NO | NO | YES | NO | P |
| RHOG Rho-related GTP-binding protein RhoG | P84096 | NO | NO | YES | YES | YES | YES | P |
| RL19 60S ribosomal protein L19 | P84099 | YES | NO | YES | YES | YES | YES | P/MQ |
| SRSF3 Serine/arginine-rich splicing factor 3 | P84104 | YES | NO | YES | YES | YES | YES | P/MQ |
| H32 Histone H3.2 | P84228 | NO | NO | YES | YES | YES | YES | P/MQ |
| IC1 Plasma protease C1 inhibitor | P97290 | NO | NO | NO | NO | YES | NO | P |
| MCM2 DNA replication licensing factor MCM2 | P97310 | NO | NO | NO | YES | NO | NO | P |
| MCM6 DNA replication licensing factor MCM6 | P97311 | NO | NO | NO | NO | YES | YES | P |
| CSRP2 Cysteine and glycine-rich protein 2 | P97314 | NO | YES | NO | NO | NO | NO | P |
| CSRP1 Cysteine and glycine-rich protein 1 | P97315 | NO | NO | NO | YES | YES | YES | P/MQ |
| RS3A 40S ribosomal protein S3a | P97328 | YES | YES | NO | NO | NO | NO | P |
| RS3A 40S ribosomal protein S3a | P97351 | YES | NO | YES | YES | YES | YES | P/MQ |
| SPS2 Selenide, water dikinase 2 | P97364 | NO | YES | NO | NO | NO | NO | P |
| NCF4 Neutrophil cytosol factor 4 | P97369 | NO | NO | NO | YES | YES | YES | P |
| PSME1 Proteasome activator complex subunit 1 | P97371 | YES | NO | YES | YES | YES | YES | P/MQ |
| PSME2 Proteasome activator complex subunit 2 | P97372 | YES | NO | YES | YES | YES | YES | P/MQ |
| G3BP2 Ras GTPase-activating protein-binding protein 2 | P97379 | NO | NO | YES | YES | NO | YES | P |
| ANX11 Annexin A11 | P97384 | YES | NO | YES | YES | YES | YES | P/MQ |
| ANXA4 Annexin A4 | P97429 | YES | NO | YES | YES | YES | YES | P/MQ |

| | | | | | | | | |
|---|---|---|---|---|---|---|---|---|
| MPRIP Myosin phosphatase Rho-interacting protein | P97434 | NO | NO | NO | YES | YES | YES | P |
| AMPN Aminopeptidase N | P97449 | YES | NO | YES | YES | YES | YES | P/MQ |
| ATP5J ATP synthase-coupling factor 6, mitochondrial | P97450 | YES | NO | YES | YES | YES | YES | P/MQ |
| RS5 40S ribosomal protein S5 | P97461 | YES | NO | YES | YES | YES | YES | P/MQ |
| GSH1 Glutamate--cysteine ligase catalytic subunit | P97494 | YES | NO | YES | NO | NO | NO | P |
| PPIP1 Proline-serine-threonine phosphatase-interacting protein 1 | P97814 | NO | NO | NO | YES | NO | YES | P |
| S100G Protein S100-G | P97816 | YES | NO | NO | NO | YES | NO | P/MQ |
| CATC Dipeptidyl peptidase 1 | P97821 | NO | YES | NO | NO | NO | NO | P/MQ |
| LYPA1 Acyl-protein thioesterase 1 | P97823 | NO | YES | NO | NO | NO | NO | P |
| JUPI1 Jupiter microtubule associated homolog 1 | P97825 | NO | NO | NO | NO | YES | NO | P |
| G3BP1 Ras GTPase-activating protein-binding protein 1 | P97855 | NO | NO | YES | YES | YES | YES | P/MQ |
| CASP7 Caspase-7 | P97864 | NO | NO | NO | YES | NO | NO | P |
| BRCA2 Breast cancer type 2 susceptibility protein homolog | P97929 | NO | YES | NO | NO | NO | NO | P |
| DAB2 Disabled homolog 2 | P98078 | YES | NO | YES | YES | YES | YES | P/MQ |
| TBB5 Tubulin beta-5 chain | P99024 | YES | NO | YES | YES | YES | YES | P/MQ |
| PSB4 Proteasome subunit beta type-4 | P99026 | YES | NO | YES | YES | YES | YES | P/MQ |
| RLA2 60S acidic ribosomal protein P2 | P99027 | YES | NO | YES | YES | YES | YES | P/MQ |
| PRDX5 Peroxiredoxin-5, mitochondrial | P99029 | YES | NO | YES | YES | YES | YES | P/MQ |
| APOA1 Apolipoprotein A-I | Q00623 | YES | NO | YES | YES | YES | YES | P/MQ |
| A1AT4 Alpha-1-antitrypsin 1-4 | Q00897 | NO | NO | YES | YES | YES | YES | P/MQ |
| RET1 Retinol-binding protein 1 | Q00915 | NO | NO | NO | YES | NO | NO | P |
| HNRL2 Heterogeneous nuclear ribonucleoprotein U-like protein 2 | Q00PI9 | NO | NO | NO | YES | NO | NO | P/MQ |
| CO1A2 Collagen alpha-2(I) chain | Q01149 | NO | NO | NO | YES | YES | YES | P |
| TOP2A DNA topoisomerase 2-alpha | Q01320 | NO | NO | NO | NO | YES | NO | P |
| SC23A Protein transport protein Sec23A | Q01405 | NO | NO | NO | YES | NO | NO | P |
| RSU1 Ras suppressor protein 1 | Q01730 | YES | NO | YES | YES | YES | YES | P |
| NDKB Nucleoside diphosphate kinase B | Q01768 | YES | NO | YES | YES | YES | YES | P/MQ |
| TERA Transitional endoplasmic reticulum ATPase | Q01853 | YES | NO | YES | YES | YES | YES | P/MQ |
| UBA1 Ubiquitin-like modifier-activating enzyme 1 | Q02053 | YES | NO | YES | YES | YES | YES | P/MQ |
| C1QC Complement C1q subcomponent subunit C | Q02105 | NO | NO | NO | YES | NO | NO | P |
| CTNB1 Catenin beta-1 | Q02248 | NO | NO | NO | YES | NO | NO | P/MQ |
| CO6A2 Collagen alpha-2(VI) chain | Q02788 | YES | NO | YES | YES | YES | YES | P |
| NUCB1 Nucleobindin-1 | Q02819 | YES | NO | YES | YES | YES | YES | P/MQ |
| ATP synthase subunit alpha, mitochondrial | Q03265 | YES | YES | YES | YES | YES | YES | MQ |
| IKZF1 DNA-binding protein Ikaros | Q03267 | NO | NO | NO | YES | NO | NO | P |
| KCRB Creatine kinase B-type | Q04447 | NO | NO | YES | YES | YES | YES | P/MQ |
| ATNG Sodium/potassium-transporting ATPase subunit gamma | Q04646 | YES | YES | NO | NO | NO | NO | P/MQ |

| | | | | | | | | |
|---|---|---|---|---|---|---|---|---|
| DNA topoisomerase 1 | Q04750 | YES | YES | YES | YES | YES | YES | MQ |
| CO6A1 Collagen alpha-1(VI) chain | Q04857 | YES | NO | YES | YES | YES | YES | P |
| RAC2 Ras-related C3 botulinum toxin substrate 2 | Q05144 | NO | NO | YES | YES | YES | YES | P/MQ |
| RCN1 Reticulocalbin-1 | Q05186 | NO | NO | YES | YES | YES | YES | P/MQ |
| MARK2 Serine/threonine-protein kinase MARK2 | Q05512 | NO | NO | NO | NO | NO | YES | P/MQ |
| PGBM Basement membrane-specific heparan sulfate proteoglycan core protein | Q05793 | YES | NO | YES | YES | YES | YES | P/MQ |
| FABP5 Fatty acid-binding protein 5 | Q05816 | NO | NO | NO | YES | YES | YES | P |
| PC Pyruvate carboxylase, mitochondrial | Q05920 | YES | NO | YES | NO | YES | NO | P/MQ |
| IF2P Eukaryotic translation initiation factor 5B | Q05D44 | NO | NO | YES | YES | YES | YES | P/MQ |
| PTN2 Tyrosine-protein phosphatase non-receptor type 2 | Q06180 | NO | YES | NO | NO | NO | NO | P |
| ATP5I ATP synthase subunit e, mitochondrial | Q06185 | YES | NO | YES | YES | YES | YES | P/MQ |
| CLUS Clusterin | Q06890 | YES | NO | YES | YES | YES | YES | P/MQ |
| ANXA7 Annexin A7 | Q07076 | YES | NO | YES | YES | YES | NO | P/MQ |
| BAX Apoptosis regulator BAX | Q07813 | NO | NO | NO | YES | YES | YES | P |
| PEBB Core-binding factor subunit beta | Q08024 | NO | NO | NO | YES | NO | NO | P |
| CNN2 Calponin-2 | Q08093 | NO | NO | YES | YES | YES | YES | P/MQ |
| SSRP1 FACT complex subunit SSRP1 | Q08943 | NO | NO | NO | NO | YES | NO | P |
| NCF1 Neutrophil cytosol factor 1 | Q09014 | NO | NO | YES | YES | YES | YES | P |
| B4GN1 Beta-1,4 N-acetylgalactosaminyltransferase 1 | Q09200 | NO | NO | NO | YES | NO | NO | P |
| APRV1 Retroviral-like aspartic protease 1 | Q09PK2 | NO | NO | NO | NO | YES | NO | P |
| ZCH18 Zinc finger CCCH domain-containing protein 18 | Q0P678 | NO | YES | NO | NO | NO | NO | P |
| RBM15 RNA-binding protein 15 | Q0VBL3 | NO | NO | NO | YES | NO | NO | P |
| PP4R2 Serine/threonine-protein phosphatase 4 regulatory subunit 2 | Q0VGB7 | NO | YES | NO | NO | NO | NO | P |
| PSA Puromycin-sensitive aminopeptidase | Q11011 | NO | NO | NO | YES | YES | YES | P/MQ |
| PEPD Xaa-Pro dipeptidase | Q11136 | YES | NO | NO | YES | YES | YES | P/MQ |
| NSUN2 tRNA (cytosine(34)-C(5))-methyltransferase | Q1HFZ0 | NO | NO | NO | YES | YES | NO | P |
| ZFHX2 Zinc finger homeobox protein 2 | Q2MHN3 | NO | YES | NO | NO | NO | NO | P |
| GSK3A Glycogen synthase kinase-3 alpha | Q2NL51 | NO | YES | NO | NO | NO | NO | P |
| HSDL2 Hydroxysteroid dehydrogenase-like protein 2 | Q2TPA8 | NO | YES | NO | NO | NO | NO | P |
| HMHA1 Rho GTPase-activating protein 45 | Q3TBD2 | NO | NO | NO | YES | NO | YES | P |
| FAHD2 Fumarylacetoacetate hydrolase domain-containing protein 2A | Q3TC72 | NO | YES | NO | NO | NO | NO | P |
| H1BP3 HCLS1-binding protein 3 | Q3TC93 | NO | YES | NO | NO | NO | NO | P |
| PPR21 Protein phosphatase 1 regulatory subunit 21 | Q3TDD9 | NO | NO | NO | YES | NO | NO | P/MQ |
| FAS-associated factor 2 | Q3TDN2 | YES | YES | YES | YES | YES | YES | MQ |
| HP1B3 Heterochromatin protein 1-binding protein 3 | Q3TEA8 | YES | NO | YES | YES | YES | YES | P/MQ |
| F107B Protein FAM107B | Q3TGF2 | YES | NO | YES | YES | YES | YES | P/MQ |
| ML12B Myosin regulatory light chain 12B | Q3THE2 | YES | NO | YES | YES | YES | YES | P/MQ |
| T2FA General transcription factor IIF subunit 1 | Q3THK3 | NO | NO | NO | YES | YES | YES | P/MQ |

| | | | | | | | | |
|---|---|---|---|---|---|---|---|---|
| GUAA GMP synthase [glutamine-hydrolyzing] | Q3THK7 | NO | NO | NO | YES | YES | NO | P |
| ZC3HF Zinc finger CCCH domain-containing protein 15 | Q3TIV5 | NO | YES | NO | NO | NO | NO | P |
| SNUT2 U4/U6.U5 tri-snRNP-associated protein 2 | Q3TIX9 | NO | YES | NO | NO | NO | NO | P |
| PDLI7 PDZ and LIM domain protein 7 | Q3TJD7 | NO | NO | NO | YES | NO | NO | P |
| SMCA4 Transcription activator BRG1 | Q3TKT4 | NO | NO | YES | YES | NO | YES | P |
| PRC2C Protein PRRC2C | Q3TLH4 | NO | NO | YES | YES | YES | YES | P/MQ |
| ECHD2 Enoyl-CoA hydratase domain-containing protein 2, mitochondrial | Q3TLP5 | YES | YES | NO | NO | NO | NO | P/MQ |
| XYLB Xylulose kinase | Q3TNA1 | YES | YES | NO | NO | NO | NO | P |
| UAP1L UDP-N-acetylhexosamine pyrophosphorylase-like protein 1 | Q3TW96 | YES | NO | YES | YES | YES | YES | P/MQ |
| SRSF6 Serine/arginine-rich splicing factor 6 | Q3TWW8 | NO | NO | NO | YES | YES | YES | P/MQ |
| PSMD1 26S proteasome non-ATPase regulatory subunit 1 | Q3TXS7 | NO | NO | NO | YES | YES | NO | P |
| ESYT2 Extended synaptotagmin-2 | Q3TZZ7 | NO | NO | NO | YES | NO | NO | P |
| FUBP2 Far upstream element-binding protein 2 | Q3U0V1 | YES | NO | YES | YES | YES | YES | P/MQ |
| DDB1 DNA damage-binding protein 1 | Q3U1J4 | NO | NO | NO | YES | YES | NO | P |
| F16A2 FTS and Hook-interacting protein | Q3U2I3 | NO | YES | NO | NO | NO | NO | P |
| PYRD2 Pyridine nucleotide-disulfide oxidoreductase domain-containing protein 2 | Q3U4I7 | NO | YES | NO | NO | NO | NO | P |
| CMPK2 UMP-CMP kinase 2, mitochondrial | Q3U5Q7 | NO | NO | NO | NO | YES | NO | P |
| ESYT1 Extended synaptotagmin-1 | Q3U7R1 | NO | NO | NO | YES | YES | YES | P |
| FBX7 F-box only protein 7 | Q3U7U3 | NO | YES | NO | NO | NO | NO | P |
| CD44 antigen | Q3U8S1 | NO | YES | YES | YES | NO | YES | MQ |
| LBR Delta(14)-sterol reductase | Q3U9G9 | NO | NO | YES | YES | YES | YES | P/MQ |
| TM109 Transmembrane protein 109 | Q3UBX0 | NO | YES | NO | NO | NO | NO | P |
| PUF60 Poly(U)-binding-splicing factor PUF60 | Q3UEB3 | NO | NO | NO | NO | YES | NO | P |
| LYPL1 Lysophospholipase-like protein 1 | Q3UFF7 | NO | YES | NO | NO | NO | NO | P |
| EI3JA Eukaryotic translation initiation factor 3 subunit J-A | Q3UGC7 | NO | NO | YES | YES | YES | YES | P/MQ |
| NEK10 Serine/threonine-protein kinase Nek10 | Q3UGM2 | NO | YES | NO | NO | NO | NO | P |
| AAK1 AP2-associated protein kinase 1 | Q3UHJ0 | YES | NO | YES | YES | YES | YES | P/MQ |
| TEN4 Teneurin-4 | Q3UHK6 | NO | YES | NO | NO | NO | NO | P |
| Fibronectin | Q3UHL6 | YES | YES | YES | YES | YES | YES | MQ |
| HAP28 28 kDa heat- and acid-stable phosphoprotein | Q3UHX2 | YES | NO | YES | YES | YES | YES | P/MQ |
| RHG17 Rho GTPase-activating protein 17 | Q3UIA2 | NO | NO | NO | YES | NO | NO | P |
| PKHA7 Pleckstrin homology domain-containing family A member 7 | Q3UIL6 | NO | YES | NO | NO | NO | NO | P |
| NDUB6 NADH dehydrogenase [ubiquinone] 1 beta subcomplex subunit 6 | Q3UIU2 | YES | NO | NO | NO | NO | NO | P |
| Splicing factor 3b, subunit 2 | Q3UJB0 | YES | YES | YES | YES | YES | YES | MQ |
| EDC4 Enhancer of mRNA-decapping protein 4 | Q3UJB9 | NO | NO | NO | YES | YES | NO | P |
| RMD3 Regulator of microtubule dynamics protein 3 | Q3UJU9 | NO | YES | NO | NO | NO | NO | P |

| | | | | | | | | |
|---|---|---|---|---|---|---|---|---|
| MCCB Methylcrotonoyl-CoA carboxylase beta chain, mitochondrial | Q3ULD5 | YES | NO | YES | NO | YES | NO | P |
| PP1R7 Protein phosphatase 1 regulatory subunit 7 | Q3UM45 | NO | NO | NO | NO | YES | NO | P |
| COBL1 Cordon-bleu protein-like 1 | Q3UMF0 | YES | NO | NO | NO | YES | NO | P |
| PP12C Protein phosphatase 1 regulatory subunit 12C | Q3UMT1 | NO | NO | NO | YES | NO | NO | P |
| HDGR2 Hepatoma-derived growth factor-related protein 2 | Q3UMU9 | NO | NO | YES | YES | NO | NO | P/MQ |
| EMAL4 Echinoderm microtubule-associated protein-like 4 | Q3UMY5 | NO | NO | NO | YES | NO | NO | P |
| SKAP2 Src kinase-associated phosphoprotein 2 | Q3UND0 | NO | NO | NO | NO | NO | YES | P |
| QORL2 Quinone oxidoreductase-like protein 2 | Q3UNZ8 | YES | YES | NO | NO | NO | NO | P |
| ELANE Neutrophil elastase | Q3UP87 | NO | NO | NO | YES | YES | YES | P |
| ZCCHV Zinc finger CCCH-type antiviral protein 1 | Q3UPF5 | NO | NO | NO | NO | YES | NO | P |
| PRRC1 Protein PRRC1 | Q3UPH1 | NO | NO | NO | NO | YES | NO | P |
| SC31A Protein transport protein Sec31A | Q3UPL0 | YES | NO | YES | YES | YES | YES | P/MQ |
| IQGA2 Ras GTPase-activating-like protein IQGAP2 | Q3UQ44 | YES | NO | NO | YES | YES | NO | P/MQ |
| ACSF3 Acyl-CoA synthetase family member 3, mitochondrial | Q3URE1 | NO | YES | NO | NO | NO | NO | P |
| SRBS2 Sorbin and SH3 domain-containing protein 2 | Q3UTJ2 | YES | NO | YES | YES | YES | YES | P/MQ |
| NIBAN Protein Niban | Q3UW53 | NO | NO | YES | YES | YES | YES | P/MQ |
| LRRF1 Leucine-rich repeat flightless-interacting protein 1 | Q3UZ39 | NO | NO | YES | YES | YES | YES | P/MQ |
| CPZIP CapZ-interacting protein | Q3UZA1 | NO | NO | NO | YES | YES | YES | P |
| ST1D1 Sulfotransferase 1 family member D1 | Q3UZZ6 | YES | NO | NO | NO | YES | NO | P/MQ |
| PLSI Plastin-1 | Q3V0K9 | YES | YES | NO | NO | NO | NO | P |
| C1TM Monofunctional C1-tetrahydrofolate synthase, mitochondrial | Q3V3R1 | NO | NO | NO | YES | NO | NO | P |
| SC5AC Sodium-coupled monocarboxylate transporter 2 | Q49B93 | NO | YES | NO | NO | NO | NO | P |
| OXR1 Oxidation resistance protein 1 | Q4KMM3 | NO | YES | NO | NO | NO | NO | P |
| ARAP1 Arf-GAP with Rho-GAP domain, ANK repeat and PH domain-containing protein 1 | Q4LDD4 | NO | NO | NO | YES | NO | NO | P |
| CC88B Coiled-coil domain-containing protein 88B | Q4QRL3 | NO | NO | NO | YES | NO | YES | P |
| CDV3 Protein CDV3 | Q4VAA2 | NO | NO | NO | YES | NO | YES | P |
| DDX17 Probable ATP-dependent RNA helicase DDX17 | Q501J6 | YES | NO | YES | YES | YES | YES | P/MQ |
| PHAR4 Phosphatase and actin regulator 4 | Q501J7 | YES | NO | NO | YES | YES | NO | P |
| LRC47 Leucine-rich repeat-containing protein 47 | Q505F5 | NO | NO | YES | YES | YES | YES | P/MQ |
| SRRM1 Serine/arginine repetitive matrix protein 1 | Q52KI8 | NO | NO | NO | YES | NO | NO | P |
| DDX46 Probable ATP-dependent RNA helicase DDX46 | Q569Z5 | NO | NO | NO | YES | YES | YES | P |
| TR150 Thyroid hormone receptor-associated protein 3 | Q569Z6 | YES | NO | YES | YES | YES | YES | P/MQ |
| JIP4 C-Jun-amino-terminal kinase-interacting protein 4 | Q58A65 | NO | NO | NO | YES | YES | NO | P |
| NDUF2 NADH dehydrogenase [ubiquinone] 1 alpha subcomplex assembly factor 2 | Q59J78 | NO | YES | NO | NO | NO | NO | P |

| | | | | | | | | |
|---|---|---|---|---|---|---|---|---|
| BRE1A E3 ubiquitin-protein ligase BRE1A | Q5DTM8 | NO | NO | NO | NO | YES | NO | P |
| AAPK1 5'-AMP-activated protein kinase catalytic subunit alpha-1 | Q5EG47 | NO | NO | NO | YES | NO | YES | P |
| NUFP2 Nuclear fragile X mental retardation-interacting protein 2 | Q5F2E7 | NO | NO | NO | YES | NO | YES | P |
| RHG01 Rho GTPase-activating protein 1 | Q5FWK3 | YES | NO | YES | YES | YES | YES | P/MQ |
| LRC52 Leucine-rich repeat-containing protein 52 | Q5M8M9 | NO | YES | NO | NO | NO | NO | P |
| DIRA2 GTP-binding protein Di-Ras2 | Q5PR73 | NO | YES | NO | NO | NO | NO | P |
| HAVR1 Hepatitis A virus cellular receptor 1 homolog | Q5QNS5 | NO | NO | NO | NO | YES | NO | P/MQ |
| IF2B2 Insulin-like growth factor 2 mRNA-binding protein 2 | Q5SF07 | NO | YES | NO | NO | NO | NO | P |
| CYFP2 Cytoplasmic FMR1-interacting protein 2 | Q5SQX6 | NO | NO | NO | YES | NO | NO | P |
| TM1L2 TOM1-like protein 2 | Q5SRX1 | NO | YES | NO | NO | NO | NO | P |
| ABR Active breakpoint cluster region-related protein | Q5SSL4 | NO | NO | NO | YES | NO | NO | P |
| MYO1G Unconventional myosin-Ig | Q5SUA5 | NO | NO | NO | YES | NO | YES | P |
| LC7L3 Luc7-like protein 3 | Q5SUF2 | NO | YES | NO | NO | NO | NO | P |
| PUR4 Phosphoribosylformylglycinamidine synthase | Q5SUR0 | NO | NO | NO | YES | YES | NO | P/MQ |
| CLU Clustered mitochondria protein homolog | Q5SW19 | YES | NO | NO | NO | YES | NO | P |
| CYTSB Cytospin-B | Q5SXY1 | NO | NO | NO | YES | NO | NO | P |
| RABL6 Rab-like protein 6 | Q5U3K5 | NO | NO | NO | NO | YES | NO | P/MQ |
| GASP1 G-protein coupled receptor-associated sorting protein 1 | Q5U4C1 | NO | YES | NO | NO | NO | NO | P |
| HYKK Hydroxylysine kinase | Q5U5V2 | NO | YES | NO | NO | NO | NO | P |
| ACBD5 Acyl-CoA-binding domain-containing protein 5 | Q5XG73 | NO | YES | NO | NO | NO | NO | P |
| PARL Presenilins-associated rhomboid-like protein, mitochondrial | Q5XJY4 | NO | YES | NO | NO | NO | NO | P |
| COPD Coatomer subunit delta | Q5XJY5 | NO | NO | YES | YES | YES | NO | P/MQ |
| A1CF APOBEC1 complementation factor | Q5YD48 | NO | YES | NO | NO | NO | NO | P |
| A1AG1 Alpha-1-acid glycoprotein 1 | Q60590 | YES | NO | YES | YES | YES | YES | P/MQ |
| SRC8 Src substrate cortactin | Q60598 | YES | NO | YES | YES | YES | YES | P/MQ |
| ADSV Adseverin | Q60604 | NO | NO | YES | YES | YES | YES | P |
| MYL6 Myosin light polypeptide 6 | Q60605 | YES | NO | YES | YES | YES | YES | P/MQ |
| GRB2 Growth factor receptor-bound protein 2 | Q60631 | NO | NO | NO | YES | YES | YES | P |
| FLOT2 Flotillin-2 | Q60634 | NO | NO | YES | YES | YES | YES | P/MQ |
| SAP3 Ganglioside GM2 activator | Q60648 | NO | NO | NO | YES | NO | NO | P/MQ |
| HNRPD Heterogeneous nuclear ribonucleoprotein D0 | Q60668 | YES | NO | YES | YES | YES | YES | P/MQ |
| PSB6 Proteasome subunit beta type-6 | Q60692 | NO | NO | NO | YES | NO | NO | P |
| SAMH1 Deoxynucleoside triphosphate triphosphohydrolase SAMHD1 | Q60710 | NO | NO | YES | YES | YES | YES | P/MQ |
| P4HA1 Prolyl 4-hydroxylase subunit alpha-1 | Q60715 | NO | NO | NO | YES | YES | NO | P |
| KHDR1 KH domain-containing, RNA-binding, signal transduction-associated protein 1 | Q60749 | NO | NO | YES | YES | NO | YES | P |
| GCDH Glutaryl-CoA dehydrogenase, mitochondrial | Q60759 | NO | YES | NO | NO | NO | NO | P |

| | | | | | | | | |
|---|---|---|---|---|---|---|---|---|
| IRGM1 Immunity-related GTPase family M protein 1 | Q60766 | NO | NO | YES | YES | YES | NO | P/MQ |
| NACA Nascent polypeptide-associated complex subunit alpha | Q60817 | YES | NO | YES | NO | YES | YES | P/MQ |
| NPT2A Sodium-dependent phosphate transport protein 2A | Q60825 | NO | YES | NO | NO | NO | NO | P |
| COCA1 Collagen alpha-1(XII) chain | Q60847 | NO | NO | NO | YES | NO | YES | P |
| STIP1 Stress-induced-phosphoprotein 1 | Q60864 | YES | NO | YES | YES | YES | YES | P/MQ |
| CAPR1 Caprin-1 | Q60865 | YES | NO | YES | YES | YES | YES | P/MQ |
| PTER Phosphotriesterase-related protein | Q60866 | YES | NO | YES | NO | NO | NO | P/MQ |
| ARHG2 Rho guanine nucleotide exchange factor 2 | Q60875 | NO | NO | NO | YES | NO | NO | P |
| EP15R Epidermal growth factor receptor substrate 15-like 1 | Q60902 | YES | NO | YES | YES | YES | YES | P/MQ |
| VDAC2 Voltage-dependent anion-selective channel protein 2 | Q60930 | YES | NO | YES | YES | YES | YES | P/MQ |
| VDAC1 Voltage-dependent anion-selective channel protein 1 | Q60932 | YES | NO | YES | YES | YES | YES | P/MQ |
| COQ8A Atypical kinase COQ8A, mitochondrial | Q60936 | NO | YES | NO | NO | NO | NO | P |
| PAFA Platelet-activating factor acetylhydrolase | Q60963 | NO | NO | NO | NO | YES | NO | P |
| PAPS1 Bifunctional 3'-phosphoadenosine 5'-phosphosulfate synthase 1 | Q60967 | NO | YES | NO | NO | NO | NO | P |
| Histone-binding protein RBBP4 | Q60972 | YES | YES | YES | YES | YES | YES | MQ |
| NCOR1 Nuclear receptor corepressor 1 | Q60974 | NO | YES | NO | NO | NO | NO | P |
| LAMA5 Laminin subunit alpha-5 | Q61001 | NO | NO | YES | YES | NO | YES | P/MQ |
| CD6 T-cell differentiation antigen CD6 | Q61003 | NO | YES | NO | NO | NO | NO | P |
| LAP2B Lamina-associated polypeptide 2, isoforms beta/delta/epsilon/gamma | Q61029 | YES | NO | NO | YES | YES | YES | P/MQ |
| LAP2A Lamina-associated polypeptide 2, isoforms alpha/zeta | Q61033 | NO | NO | NO | YES | YES | YES | P |
| SYHC Histidine--tRNA ligase, cytoplasmic | Q61035 | YES | NO | NO | YES | YES | YES | P/MQ |
| PPM1G Protein phosphatase 1G | Q61074 | NO | NO | NO | YES | YES | YES | P/MQ |
| CDC37 Hsp90 co-chaperone Cdc37 | Q61081 | NO | NO | YES | YES | YES | YES | P/MQ |
| CY24B Cytochrome b-245 heavy chain | Q61093 | NO | NO | NO | YES | NO | YES | P |
| REQU Zinc finger protein ubi-d4 | Q61103 | NO | NO | NO | NO | YES | NO | P |
| CFAI Complement factor I | Q61129 | NO | YES | NO | NO | NO | NO | P/MQ |
| CERU Ceruloplasmin | Q61147 | YES | NO | YES | YES | YES | YES | P |
| 2A5E Serine/threonine-protein phosphatase 2A 56 kDa regulatory subunit epsilon isoform | Q61151 | NO | NO | NO | YES | NO | NO | P |
| PTN18 Tyrosine-protein phosphatase non-receptor type 18 | Q61152 | NO | NO | NO | YES | NO | NO | P |
| SL9A1 Sodium/hydrogen exchanger 1 | Q61165 | YES | YES | NO | NO | NO | NO | P |
| MARE1 Microtubule-associated protein RP/EB family member 1 | Q61166 | YES | NO | YES | YES | YES | YES | P/MQ |
| PRDX2 Peroxiredoxin-2 | Q61171 | YES | NO | YES | YES | YES | YES | P/MQ |
| ARG1 Arginase-1 | Q61176 | NO | NO | YES | YES | YES | YES | P/MQ |

| | | | | | | | | |
|---|---|---|---|---|---|---|---|---|
| PAPOA Poly(A) polymerase alpha | Q61183 | NO | YES | NO | NO | NO | NO | P |
| TS101 Tumor susceptibility gene 101 protein | Q61187 | NO | NO | NO | YES | NO | NO | P |
| ICLN Methylosome subunit pICln | Q61189 | NO | NO | NO | NO | YES | NO | P |
| HCFC1 Host cell factor 1 | Q61191 | NO | NO | YES | YES | YES | YES | P/MQ |
| PA1B2 Platelet-activating factor acetylhydrolase IB subunit beta | Q61206 | YES | NO | YES | NO | NO | NO | P/MQ |
| SAP Prosaposin | Q61207 | NO | YES | NO | NO | NO | NO | P |
| ARHG1 Rho guanine nucleotide exchange factor 1 | Q61210 | NO | NO | NO | YES | YES | YES | P |
| PLSL Plastin-2 | Q61233 | NO | NO | YES | YES | YES | YES | P/MQ |
| SNTB2 Beta-2-syntrophin | Q61235 | NO | YES | NO | NO | NO | NO | P |
| A2AP Alpha-2-antiplasmin | Q61247 | YES | NO | YES | YES | YES | YES | P/MQ |
| IGBP1 Immunoglobulin-binding protein 1 | Q61249 | NO | NO | YES | YES | YES | YES | P/MQ |
| LAMB2 Laminin subunit beta-2 | Q61292 | YES | NO | NO | NO | YES | NO | P |
| HSP74 Heat shock 70 kDa protein 4 | Q61316 | YES | NO | YES | YES | YES | YES | P/MQ |
| BAP31 B-cell receptor-associated protein 31 | Q61335 | NO | NO | YES | NO | YES | YES | P |
| CH3L1 Chitinase-3-like protein 1 | Q61362 | NO | NO | NO | YES | NO | NO | P |
| CY24A Cytochrome b-245 light chain | Q61462 | NO | NO | NO | YES | YES | YES | P |
| ECM1 Extracellular matrix protein 1 | Q61508 | NO | NO | YES | YES | YES | NO | P/MQ |
| TRI25 E3 ubiquitin/ISG15 ligase TRIM25 | Q61510 | NO | YES | NO | NO | NO | NO | P |
| EWS RNA-binding protein EWS | Q61545 | YES | NO | YES | YES | YES | YES | P/MQ |
| RAD21 Double-strand-break repair protein rad21 homolog | Q61550 | NO | NO | NO | NO | YES | NO | P |
| FSCN1 Fascin | Q61553 | NO | YES | NO | NO | NO | NO | P |
| FBN1 Fibrillin-1 | Q61554 | NO | NO | NO | YES | NO | NO | P |
| IBP7 Insulin-like growth factor-binding protein 7 | Q61581 | NO | YES | NO | NO | NO | NO | P |
| FXR1 Fragile X mental retardation syndrome-related protein 1 | Q61584 | NO | NO | NO | YES | NO | NO | P |
| KTN1 Kinectin | Q61595 | NO | NO | NO | NO | NO | YES | P |
| GDIB Rab GDP dissociation inhibitor beta | Q61598 | NO | NO | YES | YES | YES | YES | P/MQ |
| GDIR2 Rho GDP-dissociation inhibitor 2 | Q61599 | NO | NO | YES | YES | YES | YES | P/MQ |
| GTP-binding protein | Q61635 | NO | YES | YES | YES | NO | YES | MQ |
| HP Haptoglobin | Q61646 | YES | NO | YES | YES | YES | YES | P/MQ |
| DD19A ATP-dependent RNA helicase DDX19A | Q61655 | NO | NO | NO | YES | YES | YES | P/MQ |
| DDX5 Probable ATP-dependent RNA helicase DDX5 | Q61656 | YES | NO | YES | YES | YES | YES | P/MQ |
| ATRX Transcriptional regulator ATRX | Q61687 | NO | YES | NO | NO | NO | NO | P |
| HS71A Heat shock 70 kDa protein 1A | Q61696 | NO | NO | NO | YES | YES | YES | P/MQ |
| HS105 Heat shock protein 105 kDa | Q61699 | YES | NO | YES | YES | YES | YES | P/MQ |
| ITIH2 Inter-alpha-trypsin inhibitor heavy chain H2 | Q61703 | NO | YES | NO | NO | NO | NO | P |
| ITIH3 Inter-alpha-trypsin inhibitor heavy chain H3 | Q61704 | YES | NO | YES | YES | YES | YES | P/MQ |
| ITA6 Integrin alpha-6 | Q61739 | YES | NO | NO | NO | YES | NO | P/MQ |
| EI2BD Translation initiation factor eIF-2B subunit delta | Q61749 | NO | YES | NO | NO | NO | NO | P |
| SERA D-3-phosphoglycerate dehydrogenase | Q61753 | NO | NO | YES | YES | YES | YES | P/MQ |

| | | | | | | | | |
|---|---|---|---|---|---|---|---|---|
| 3BHS4 NADPH-dependent 3-keto-steroid reductase Hsd3b4 | Q61767 | YES | YES | NO | NO | NO | NO | P |
| KINH Kinesin-1 heavy chain | Q61768 | YES | NO | YES | YES | YES | YES | P/MQ |
| LASP1 LIM and SH3 domain protein 1 | Q61792 | YES | NO | YES | YES | YES | YES | P/MQ |
| PZP Pregnancy zone protein | Q61838 | YES | NO | YES | YES | YES | YES | P/MQ |
| MEP1B Meprin A subunit beta | Q61847 | NO | YES | NO | NO | NO | NO | P |
| MYH10 Myosin-10 | Q61879 | YES | NO | YES | YES | YES | YES | P/MQ |
| MCM7 DNA replication licensing factor MCM7 | Q61881 | NO | NO | NO | YES | NO | NO | P |
| NPM Nucleophosmin | Q61937 | YES | NO | YES | YES | YES | YES | P/MQ |
| PCBP2 Poly(rC)-binding protein 2 | Q61990 | NO | YES | NO | NO | NO | NO | P |
| POSTN Periostin | Q62009 | NO | NO | NO | YES | NO | NO | P |
| PEA15 Astrocytic phosphoprotein PEA-15 | Q62048 | NO | YES | NO | NO | NO | NO | P |
| SRSF2 Serine/arginine-rich splicing factor 2 | Q62093 | NO | NO | YES | YES | YES | YES | P/MQ |
| AL1A2 Retinal dehydrogenase 2 | Q62148 | NO | NO | YES | YES | YES | NO | P/MQ |
| DDX3X ATP-dependent RNA helicase DDX3X | Q62167 | YES | NO | YES | YES | YES | YES | P/MQ |
| SSRD Translocon-associated protein subunit delta | Q62186 | NO | NO | NO | YES | YES | NO | P |
| DPYL3 Dihydropyrimidinase-related protein 3 | Q62188 | YES | NO | YES | YES | YES | YES | P |
| SNRPA U1 small nuclear ribonucleoprotein A | Q62189 | NO | NO | NO | YES | NO | NO | P |
| SPTB2 Spectrin beta chain, non-erythrocytic 1 | Q62261 | YES | NO | YES | YES | YES | YES | P/MQ |
| TGTP1 T-cell-specific guanine nucleotide triphosphate-binding protein 1 | Q62293 | NO | NO | YES | YES | YES | YES | P |
| TIF1B Transcription intermediary factor 1-beta | Q62318 | NO | NO | NO | YES | YES | YES | P/MQ |
| RU17 U1 small nuclear ribonucleoprotein 70 kDa | Q62376 | YES | NO | YES | YES | YES | YES | P/MQ |
| TPD52 Tumor protein D52 | Q62393 | YES | NO | YES | YES | YES | YES | P/MQ |
| SRBS1 Sorbin and SH3 domain-containing protein 1 | Q62417 | YES | NO | YES | YES | YES | YES | P |
| Drebrin-like protein | Q62418 | YES | YES | YES | YES | YES | YES | MQ |
| SH3G1 Endophilin-A2 | Q62419 | NO | NO | NO | YES | NO | NO | P |
| OSTF1 Osteoclast-stimulating factor 1 | Q62422 | NO | NO | YES | YES | YES | YES | P |
| NDUA4 Cytochrome c oxidase subunit NDUFA4 | Q62425 | YES | YES | NO | NO | NO | NO | P |
| CYTB Cystatin-B | Q62426 | YES | NO | YES | YES | YES | YES | P/MQ |
| SMAD2 Mothers against decapentaplegic homolog 2 | Q62432 | NO | NO | NO | YES | NO | YES | P/MQ |
| NDRG1 Protein NDRG1 | Q62433 | YES | NO | YES | YES | YES | YES | P/MQ |
| FKBP3 Peptidyl-prolyl cis-trans isomerase FKBP3 | Q62446 | YES | NO | YES | YES | YES | YES | P/MQ |
| IF4G2 Eukaryotic translation initiation factor 4 gamma 2 | Q62448 | YES | NO | NO | YES | YES | YES | P/MQ |
| VAT1 Synaptic vesicle membrane protein VAT-1 homolog | Q62465 | YES | NO | YES | YES | YES | YES | P/MQ |
| VILI Villin-1 | Q62468 | YES | NO | YES | YES | YES | YES | P/MQ |
| ITA3 Integrin alpha-3 | Q62470 | NO | YES | NO | NO | NO | NO | P |
| ZYX Zyxin | Q62523 | NO | NO | YES | YES | YES | YES | P/MQ |
| SBP2 Selenium-binding protein 2 | Q63836 | NO | NO | NO | YES | NO | NO | P |
| MK03 Mitogen-activated protein kinase 3 | Q63844 | NO | NO | NO | YES | YES | YES | P |

| | | | | | | | | |
|---|---|---|---|---|---|---|---|---|
| CAVN2 Caveolae-associated protein 2 | Q63918 | NO | YES | NO | NO | NO | NO | P |
| MP2K2 Dual specificity mitogen-activated protein kinase kinase 2 | Q63932 | NO | NO | NO | YES | YES | NO | P |
| INADL InaD-like protein | Q63ZW7 | NO | YES | NO | NO | NO | NO | P |
| CRK Adapter molecule crk | Q64010 | YES | NO | NO | YES | YES | YES | P |
| RALY RNA-binding protein Raly | Q64012 | YES | NO | NO | YES | YES | YES | P/MQ |
| AEBP1 Adipocyte enhancer-binding protein 1 | Q640N1 | NO | NO | NO | YES | YES | NO | P/MQ |
| SPRE Sepiapterin reductase | Q64105 | YES | NO | YES | YES | YES | YES | P/MQ |
| SF01 Splicing factor 1 | Q64213 | YES | NO | NO | YES | YES | YES | P/MQ |
| STXB2 Syntaxin-binding protein 2 | Q64324 | NO | NO | NO | YES | YES | YES | P/MQ |
| MYO6 Unconventional myosin-VI | Q64331 | YES | YES | NO | NO | NO | NO | P |
| FKBP5 Peptidyl-prolyl cis-trans isomerase FKBP5 | Q64378 | NO | NO | NO | NO | YES | NO | P |
| CH10 10 kDa heat shock protein, mitochondrial | Q64433 | YES | NO | YES | YES | YES | YES | P/MQ |
| ATP4A Potassium-transporting ATPase alpha chain 1 | Q64436 | NO | YES | NO | NO | NO | NO | P |
| SORDp Sorbitol dehydrogenase | Q64442 | YES | NO | YES | YES | YES | NO | P/MQ |
| CP4B1 Cytochrome P450 4B1 | Q64462 | NO | YES | NO | NO | NO | NO | P |
| Histone H2B type 1-B | Q64475 | YES | YES | YES | YES | YES | YES | MQ |
| TPP2 Tripeptidyl-peptidase 2 | Q64514 | NO | NO | NO | YES | YES | NO | P/MQ |
| AT2A3 Sarcoplasmic/endoplasmic reticulum calcium ATPase 3 | Q64518 | NO | NO | NO | YES | NO | NO | P |
| H2A2C Histone H2A type 2-C | Q64523 | YES | NO | NO | YES | YES | YES | P/MQ |
| H2B2B Histone H2B type 2-B | Q64525 | NO | YES | NO | NO | NO | NO | P |
| NQO1 NAD(P)H dehydrogenase [quinone] 1 | Q64669 | NO | YES | NO | NO | NO | NO | P |
| SRM Spermidine synthase | Q64674 | NO | NO | NO | YES | YES | NO | P/MQ |
| VINC Vinculin | Q64727 | YES | NO | YES | YES | YES | YES | P/MQ |
| RAB3I Rab-3A-interacting protein | Q68EF0 | NO | YES | NO | NO | NO | NO | P |
| CLH1 Clathrin heavy chain 1 | Q68FD5 | YES | NO | YES | YES | YES | YES | P/MQ |
| PKP4 Plakophilin-4 | Q68FH0 | YES | YES | NO | NO | NO | NO | P |
| SAHH3 Putative adenosylhomocysteinase 3 | Q68FL4 | YES | NO | YES | YES | YES | YES | P/MQ |
| SYMC Methionine--tRNA ligase, cytoplasmic | Q68FL6 | NO | NO | NO | YES | NO | YES | P |
| MYOF Myoferlin | Q69ZN7 | NO | NO | YES | YES | YES | YES | P |
| PDS5A Sister chromatid cohesion protein PDS5 homolog A | Q6A026 | NO | NO | YES | YES | YES | YES | P/MQ |
| SWP70 Switch-associated protein 70 | Q6A028 | NO | NO | NO | YES | NO | NO | P |
| CE170 Centrosomal protein of 170 kDa | Q6A065 | NO | NO | NO | YES | NO | NO | P |
| CDC5L Cell division cycle 5-like protein | Q6A068 | NO | NO | NO | YES | YES | YES | P |
| LAR4B La-related protein 4B | Q6A0A2 | NO | NO | NO | YES | YES | NO | P |
| F120A Constitutive coactivator of PPAR-gamma-like protein 1 | Q6A0A9 | YES | NO | YES | YES | YES | YES | P/MQ |
| RFTN1 Raftlin | Q6A0D4 | NO | NO | NO | YES | NO | NO | P |
| CGNL1 Cingulin-like protein 1 | Q6AW69 | YES | NO | NO | YES | NO | NO | P/MQ |
| PRAM PML-RARA-regulated adapter molecule 1 | Q6BCL1 | NO | NO | NO | YES | YES | YES | P |

| | | | | | | | | |
|---|---|---|---|---|---|---|---|---|
| NOP58 Nucleolar protein 58 | Q6DFW4 | NO | NO | NO | YES | NO | NO | P/MQ |
| PFKFB4 6-phosphofructo-2-kinase/fructose-2,6-bisphosphatase 4 | Q6DTY7 | NO | NO | NO | YES | NO | NO | P |
| LEMD2 LEM domain-containing protein 2 | Q6DVA0 | NO | YES | NO | NO | NO | NO | P |
| ENPP3 Ectonucleotide pyrophosphatase/phosphodiesterase family member 3 | Q6DYE8 | NO | YES | NO | NO | NO | NO | P |
| A2MG Alpha-2-macroglobulin-P | Q6GQT1 | YES | NO | YES | NO | YES | NO | P |
| NOMO1 nodal modulator 1 | Q6GQT9 | NO | NO | YES | YES | YES | YES | P |
| RGPA1 Ral GTPase-activating protein subunit alpha-1 | Q6GYP7 | NO | YES | NO | NO | NO | NO | P |
| TPM4 Tropomyosin alpha-4 chain | Q6IRU2 | YES | NO | YES | YES | YES | YES | P/MQ |
| CLCB Clathrin light chain B | Q6IRU5 | YES | NO | YES | YES | YES | YES | P/MQ |
| PEPL1 Probable aminopeptidase NPEPL1 | Q6NSR8 | NO | YES | NO | NO | NO | NO | P |
| CPSF6 Cleavage and polyadenylation specificity factor subunit 6 | Q6NVF9 | NO | NO | NO | YES | NO | YES | P/MQ |
| DNJC8 DnaJ homolog subfamily C member 8 | Q6NZB0 | NO | NO | NO | YES | YES | YES | P/MQ |
| ZC11A Zinc finger CCCH domain-containing protein 11A | Q6NZF1 | NO | NO | NO | YES | NO | NO | P |
| IF4G1 Eukaryotic translation initiation factor 4 gamma 1 | Q6NZJ6 | YES | NO | YES | YES | YES | YES | P/MQ |
| SORCN Sorcin | Q6P069 | NO | NO | YES | YES | YES | YES | P |
| XPP1 Xaa-Pro aminopeptidase 1 | Q6P1B1 | YES | NO | NO | NO | YES | NO | P |
| ATG2A Autophagy-related protein 2 homolog A | Q6P4T0 | NO | YES | NO | NO | NO | NO | P |
| U520 U5 small nuclear ribonucleoprotein 200 kDa helicase | Q6P4T2 | NO | NO | NO | YES | YES | YES | P |
| ABCF1 ATP-binding cassette sub-family F member 1 | Q6P542 | NO | NO | NO | YES | YES | YES | P/MQ |
| UGGG1 UDP-glucose:glycoprotein glucosyltransferase 1 | Q6P5E4 | NO | NO | NO | YES | YES | NO | P |
| XPO1 Exportin-1 | Q6P5F9 | NO | YES | NO | NO | NO | NO | P/MQ |
| UBXN7 UBX domain-containing protein 7 | Q6P5G6 | NO | NO | NO | YES | YES | NO | P |
| NEST Nestin | Q6P5H2 | YES | NO | YES | YES | YES | NO | P/MQ |
| PCNP PEST proteolytic signal-containing nuclear protein | Q6P8I4 | YES | NO | YES | YES | YES | YES | P/MQ |
| SNX6 Sorting nexin-6 | Q6P8X1 | NO | NO | YES | YES | YES | YES | P/MQ |
| FHOD1 FH1/FH2 domain-containing protein 1 | Q6P9Q4 | NO | NO | NO | YES | NO | NO | P |
| FKB15 FK506-binding protein 15 | Q6P9Q6 | NO | NO | YES | YES | YES | YES | P/MQ |
| OXSR1 Serine/threonine-protein kinase OSR1 | Q6P9R2 | YES | NO | YES | YES | NO | YES | P |
| TXLNA Alpha-taxilin | Q6PAM1 | NO | YES | NO | NO | NO | NO | P/MQ |
| LPPRC Leucine-rich PPR motif-containing protein, mitochondrial | Q6PB66 | YES | NO | NO | YES | NO | NO | P |
| 2A5A Serine/threonine-protein phosphatase 2A 56 kDa regulatory subunit alpha isoform | Q6PD03 | NO | NO | NO | YES | NO | NO | P |
| SMRC2 SWI/SNF complex subunit SMARCC2 | Q6PDG5 | NO | NO | NO | YES | YES | YES | P |
| ECM29 Proteasome adapter and scaffold protein ECM29 | Q6PDI5 | NO | NO | NO | YES | YES | NO | P |

| | | | | | | | | |
|---|---|---|---|---|---|---|---|---|
| DC1L2 Cytoplasmic dynein 1 light intermediate chain 2 | Q6PDL0 | NO | NO | YES | YES | YES | YES | P |
| SRSF1 Serine/arginine-rich splicing factor 1 | Q6PDM2 | YES | NO | YES | YES | YES | YES | P/MQ |
| MYLK Myosin light chain kinase, smooth muscle | Q6PDN3 | YES | NO | YES | YES | YES | YES | P/MQ |
| CHD4 Chromodomain-helicase-DNA-binding protein 4 | Q6PDQ2 | NO | NO | NO | NO | YES | NO | P/MQ |
| AEDO 2-aminoethanethiol dioxygenase | Q6PDY2 | NO | NO | NO | YES | NO | NO | P |
| PRRT3 Proline-rich transmembrane protein 3 | Q6PE13 | NO | YES | NO | NO | NO | NO | P |
| EXOC8 Exocyst complex component 8 | Q6PGF7 | NO | YES | NO | NO | NO | NO | P |
| JUPI2 Jupiter microtubule associated homolog 2 | Q6PGH2 | NO | NO | YES | YES | YES | NO | P |
| WASC2 WASH complex subunit 2 | Q6PGL7 | YES | NO | NO | YES | YES | YES | P/MQ |
| RAB35 Ras-related protein Rab-35 | Q6PHN9 | NO | YES | NO | NO | NO | NO | P |
| YJ005 Uncharacterized protein FLJ45252 homolog | Q6PIU9 | NO | NO | NO | YES | NO | NO | P/MQ |
| AT2B4 Plasma membrane calcium-transporting ATPase 4 | Q6Q477 | NO | YES | NO | NO | NO | NO | P/MQ |
| DDX58 Probable ATP-dependent RNA helicase DDX58 | Q6Q899 | NO | NO | NO | YES | NO | NO | P |
| NEB2 Neurabin-2 | Q6R891 | NO | NO | NO | YES | NO | YES | P |
| UBR2 E3 ubiquitin-protein ligase UBR2 | Q6WKZ8 | NO | YES | NO | NO | NO | NO | P |
| KCD12 BTB/POZ domain-containing protein KCTD12 | Q6WVG3 | NO | NO | YES | YES | NO | YES | P/MQ |
| Calumenin | Q6XLQ8 | YES | YES | YES | YES | YES | YES | MQ |
| GGYF2 GRB10-interacting GYF protein 2 | Q6Y7W8 | NO | YES | NO | NO | NO | NO | P |
| PITM2 Membrane-associated phosphatidylinositol transfer protein 2 | Q6ZPQ6 | NO | YES | NO | NO | NO | NO | P |
| CAND1 Cullin-associated NEDD8-dissociated protein 1 | Q6ZQ38 | YES | NO | YES | YES | YES | YES | P/MQ |
| MLEC Malectin | Q6ZQI3 | NO | NO | YES | NO | NO | NO | P/MQ |
| UD17C UDP-glucuronosyltransferase 1-7C | Q6ZQM8 | NO | NO | NO | NO | YES | NO | P/MQ |
| RS9 40S ribosomal protein S9 | Q6ZWN5 | YES | NO | YES | YES | YES | YES | P/MQ |
| SPCS3 Signal peptidase complex subunit 3 | Q6ZWQ7 | NO | YES | NO | NO | NO | NO | P |
| RS27 40S ribosomal protein S27 | Q6ZWU9 | NO | NO | YES | YES | NO | YES | P/MQ |
| RL10 60S ribosomal protein L10 | Q6ZWV3 | NO | NO | NO | YES | YES | YES | P |
| RL35 60S ribosomal protein L35 | Q6ZWV7 | YES | NO | YES | YES | YES | YES | P/MQ |
| IF2A Eukaryotic translation initiation factor 2 subunit 1 | Q6ZWX6 | NO | NO | NO | NO | YES | YES | P/MQ |
| TYB10 Thymosin beta-10 | Q6ZWY8 | YES | NO | YES | YES | YES | YES | P |
| H2B1C Histone H2B type 1-C/E/G | Q6ZWY9 | NO | NO | YES | YES | YES | YES | P |
| TLN2 Talin-2 | Q71LX4 | NO | YES | NO | NO | NO | NO | P |
| MINY1 Ubiquitin carboxyl-terminal hydrolase MINDY-1 | Q76LS9 | NO | YES | NO | NO | NO | NO | P |
| 2AAA Serine/threonine-protein phosphatase 2A 65 kDa regulatory subunit A alpha isoform | Q76MZ3 | YES | NO | NO | YES | YES | YES | P/MQ |
| USMG5 Up-regulated during skeletal muscle growth protein 5 | Q78IK2 | NO | NO | NO | NO | YES | NO | P/MQ |
| MIC27 MICOS complex subunit Mic27 | Q78IK4 | NO | YES | NO | NO | NO | NO | P |
| 3HAO 3-hydroxyanthranilate 3,4-dioxygenase | Q78JT3 | YES | YES | NO | NO | NO | NO | P |
| SND1 Staphylococcal nuclease domain-containing protein 1 | Q78PY7 | YES | NO | YES | YES | YES | YES | P/MQ |

| | | | | | | | | |
|---|---|---|---|---|---|---|---|---|
| NP1L4 Nucleosome assembly protein 1-like 4 | Q78ZA7 | NO | NO | YES | YES | YES | YES | P/MQ |
| MTCH2 Mitochondrial carrier homolog 2 | Q791V5 | NO | YES | NO | NO | NO | NO | P |
| PICAL Phosphatidylinositol-binding clathrin assembly protein | Q7M6Y3 | YES | NO | YES | YES | YES | YES | P/MQ |
| Nuclear pore complex-associated intranuclear coiled-coil protein TPR | Q7M739 | YES | YES | YES | YES | YES | YES | MQ |
| CYFP1 Cytoplasmic FMR1-interacting protein 1 | Q7TMB8 | NO | NO | NO | YES | YES | YES | P/MQ |
| NDUAC NADH dehydrogenase [ubiquinone] 1 alpha subcomplex subunit 12 | Q7TMF3 | NO | YES | NO | NO | NO | NO | P |
| HNRPQ Heterogeneous nuclear ribonucleoprotein Q | Q7TMK9 | YES | NO | YES | YES | YES | YES | P/MQ |
| TBB2A Tubulin beta-2A chain | Q7TMM9 | YES | NO | NO | YES | YES | NO | P/MQ |
| ABCG2 ATP-binding cassette sub-family G member 2 | Q7TMS5 | NO | YES | NO | NO | NO | NO | P |
| HUWE1 E3 ubiquitin-protein ligase HUWE1 | Q7TMY8 | NO | NO | NO | YES | NO | NO | P |
| INAVA Innate immunity activator protein | Q7TN12 | NO | YES | NO | NO | NO | NO | P |
| SMAP2 Stromal membrane-associated protein 2 | Q7TN29 | NO | NO | NO | YES | NO | NO | P |
| LC7L2 Putative RNA-binding protein Luc7-like 2 | Q7TNC4 | NO | NO | NO | YES | YES | YES | P/MQ |
| DOC2A Double C2-like domain-containing protein alpha | Q7TNF0 | NO | YES | NO | NO | NO | NO | P |
| EMAL2 Echinoderm microtubule-associated protein-like 2 | Q7TNG5 | NO | NO | NO | YES | YES | YES | P |
| LDHD Probable D-lactate dehydrogenase, mitochondrial | Q7TNG8 | YES | NO | YES | NO | YES | YES | P |
| DEK Protein DEK | Q7TNV0 | NO | NO | YES | YES | YES | YES | P/MQ |
| ACTN1 Alpha-actinin-1 | Q7TPR4 | YES | NO | YES | YES | YES | YES | P/MQ |
| MBB1A Myb-binding protein 1A | Q7TPV4 | YES | NO | YES | YES | YES | YES | P/MQ |
| PKHA6 Pleckstrin homology domain-containing family A member 6 | Q7TQG1 | NO | YES | NO | NO | NO | NO | P |
| ATX2L Ataxin-2-like protein | Q7TQH0 | NO | NO | NO | YES | NO | YES | P |
| OTUB1 Ubiquitin thioesterase OTUB1 | Q7TQI3 | NO | YES | NO | NO | NO | NO | P |
| PRC2A Protein PRRC2A | Q7TSC1 | NO | NO | NO | YES | YES | YES | P |
| PP6R1 Serine/threonine-protein phosphatase 6 regulatory subunit 1 | Q7TSI3 | NO | NO | NO | YES | NO | NO | P |
| PGM2 Phosphoglucomutase-2 | Q7TSV4 | NO | NO | NO | YES | YES | YES | P/MQ |
| ENKD1 Enkurin domain-containing protein 1 | Q7TSV9 | NO | YES | NO | NO | NO | NO | P |
| MRCKB Serine/threonine-protein kinase MRCK beta | Q7TT50 | NO | YES | NO | NO | NO | NO | P |
| GVIN1 Interferon-induced very large GTPase 1 | Q80SU7 | NO | NO | YES | YES | YES | YES | P |
| SAHH2 S-adenosylhomocysteine hydrolase-like protein 1 | Q80SW1 | NO | YES | NO | NO | NO | NO | P |
| FNBP1 Formin-binding protein 1 | Q80TY0 | NO | NO | NO | YES | NO | NO | P |
| SNPH Syntaphilin | Q80U23 | NO | YES | NO | NO | NO | NO | P |
| ARHGH Rho guanine nucleotide exchange factor 17 | Q80U35 | NO | YES | NO | NO | NO | NO | P |
| UBP8 Ubiquitin carboxyl-terminal hydrolase 8 | Q80U87 | NO | NO | NO | YES | NO | NO | P |
| Band 4.1-like protein 2 | Q80UE5 | NO | YES | NO | YES | YES | YES | MQ |

| | | | | | | | | |
|---|---|---|---|---|---|---|---|---|
| SEPT9 Septin-9 | Q80UG5 | YES | NO | YES | YES | YES | YES | P/MQ |
| MOGS Mannosyl-oligosaccharide glucosidase | Q80UM7 | NO | NO | NO | YES | NO | NO | P |
| PGRC2 Membrane-associated progesterone receptor component 2 | Q80UU9 | NO | NO | NO | YES | YES | NO | P |
| FA98B Protein FAM98B | Q80VD1 | YES | YES | NO | NO | NO | NO | P |
| SRA1 Steroid receptor RNA activator 1 | Q80VJ2 | NO | NO | NO | YES | NO | NO | P |
| DNPH1 2'-deoxynucleoside 5'-phosphate N-hydrolase 1 | Q80VJ3 | NO | YES | NO | NO | NO | NO | P |
| KASH5 Protein KASH5 | Q80VJ8 | NO | YES | NO | NO | NO | NO | P |
| AL3B1 Aldehyde dehydrogenase family 3 member B1 | Q80VQ0 | NO | NO | NO | YES | YES | YES | P |
| THNS2 Threonine synthase-like 2 | Q80W22 | NO | YES | NO | NO | NO | NO | P |
| KIRR1 Kin of IRRE-like protein 1 | Q80W68 | NO | YES | NO | NO | NO | NO | P |
| AGFG2 Arf-GAP domain and FG repeat-containing protein 2 | Q80WC7 | NO | YES | NO | NO | NO | NO | P |
| LYRIC Protein LYRIC | Q80WJ7 | NO | NO | NO | YES | YES | YES | P/MQ |
| Protein VAC14 homolog | Q80WQ2 | YES | YES | YES | YES | YES | YES | MQ |
| COEA1 Collagen alpha-1(XIV) chain | Q80X19 | YES | NO | NO | YES | YES | NO | P |
| VRK1 Serine/threonine-protein kinase VRK1 | Q80X41 | NO | NO | NO | NO | NO | YES | P |
| UBP2L Ubiquitin-associated protein 2-like | Q80X50 | YES | NO | YES | YES | YES | YES | P/MQ |
| C2C2L Phospholipid transfer protein C2CD2L | Q80X80 | NO | YES | NO | NO | NO | NO | P |
| FLNB Filamin-B | Q80X90 | YES | NO | YES | YES | YES | YES | P/MQ |
| FBX34 F-box only protein 34 | Q80XI1 | NO | YES | NO | NO | NO | NO | P |
| IF4G3 Eukaryotic translation initiation factor 4 gamma 3 | Q80XI3 | NO | NO | NO | YES | NO | NO | P |
| PI42B Phosphatidylinositol 5-phosphate 4-kinase type-2 beta | Q80XI4 | NO | NO | NO | YES | NO | NO | P |
| BDH1 D-beta-hydroxybutyrate dehydrogenase, mitochondrial | Q80XN0 | NO | YES | NO | NO | NO | NO | P |
| NUCKS Nuclear ubiquitous casein and cyclin-dependent kinase substrate 1 | Q80XU3 | NO | NO | NO | YES | NO | YES | P/MQ |
| GLRX5 Glutaredoxin-related protein 5, mitochondrial | Q80Y14 | YES | NO | NO | NO | YES | NO | P |
| SPT20 Spermatogenesis-associated protein 20 | Q80YT5 | NO | YES | NO | NO | NO | NO | P |
| TENA Tenascin | Q80YX1 | NO | NO | NO | NO | YES | YES | P |
| DLG1 Disks large homolog 1 | Q811D0 | YES | NO | NO | YES | NO | NO | P |
| GALD1 Glutamine amidotransferase-like class 1 domain-containing protein 1 | Q8BFQ8 | NO | NO | NO | YES | YES | YES | P/MQ |
| GNS N-acetylglucosamine-6-sulfatase | Q8BFR4 | NO | NO | NO | YES | YES | NO | P |
| EFTU Elongation factor Tu, mitochondrial | Q8BFR5 | YES | NO | YES | YES | YES | YES | P/MQ |
| LPP Lipoma-preferred partner homolog | Q8BFW7 | YES | NO | YES | YES | YES | YES | P/MQ |
| TNPO1 Transportin-1 | Q8BFY9 | NO | NO | NO | NO | YES | NO | P |
| ACTBL Beta-actin-like protein 2 | Q8BFZ3 | NO | NO | NO | YES | YES | YES | P/MQ |
| ERLN2 Erlin-2 | Q8BFZ9 | NO | NO | NO | NO | YES | NO | P |
| ROA3 Heterogeneous nuclear ribonucleoprotein A3 | Q8BG05 | YES | NO | YES | YES | YES | YES | P/MQ |

| | | | | | | | | |
|---|---|---|---|---|---|---|---|---|
| PSD11 26S proteasome non-ATPase regulatory subunit 11 | Q8BG32 | YES | NO | YES | YES | YES | YES | P/MQ |
| PDIP3 Polymerase delta-interacting protein 3 | Q8BG81 | YES | NO | NO | YES | YES | YES | P/MQ |
| COMD7 COMM domain-containing protein 7 | Q8BG94 | NO | YES | NO | NO | NO | NO | P |
| ACSM5 Acyl-coenzyme A synthetase ACSM5, mitochondrial | Q8BGA8 | YES | NO | NO | NO | NO | NO | P |
| HTSF1 HIV Tat-specific factor 1 homolog | Q8BGC0 | NO | YES | NO | NO | NO | NO | P/MQ |
| PTGR3 Prostaglandin reductase-3 | Q8BGC4 | NO | YES | NO | NO | NO | NO | P |
| IF4B Eukaryotic translation initiation factor 4B | Q8BGD9 | YES | NO | YES | YES | YES | YES | P/MQ |
| SAM50 Sorting and assembly machinery component 50 homolog | Q8BGH2 | NO | YES | NO | NO | NO | NO | P |
| SYAC Alanine--tRNA ligase, cytoplasmic | Q8BGQ7 | NO | NO | NO | YES | YES | NO | P |
| E41L5 Band 4.1-like protein 5 | Q8BGS1 | NO | YES | NO | NO | NO | NO | P |
| TACD2 Tumor-associated calcium signal transducer 2 | Q8BGV3 | NO | YES | NO | NO | NO | NO | P |
| AL8A1 2-aminomuconic semialdehyde dehydrogenase | Q8BH00 | YES | NO | NO | NO | YES | NO | P |
| PCK2 Phosphoenolpyruvate carboxykinase [GTP], mitochondrial | Q8BH04 | NO | NO | NO | YES | NO | NO | P |
| WASF2 Wiskott-Aldrich syndrome protein family member 2 | Q8BH43 | NO | NO | NO | YES | YES | YES | P/MQ |
| CMC1 Calcium-binding mitochondrial carrier protein Aralar1 | Q8BH59 | NO | YES | NO | NO | NO | NO | P |
| F13A Coagulation factor XIII A chain | Q8BH61 | NO | NO | YES | YES | YES | YES | P/MQ |
| GLUCM D-glutamate cyclase, mitochondrial | Q8BH86 | YES | NO | NO | NO | YES | NO | P |
| RCN3 Reticulocalbin-3 | Q8BH97 | NO | NO | NO | YES | NO | NO | P |
| PTBP3 Polypyrimidine tract-binding protein 3 | Q8BHD7 | NO | NO | NO | YES | NO | YES | P |
| NRDC Nardilysin | Q8BHG1 | NO | NO | NO | NO | NO | YES | P |
| CZIB CXXC motif containing zinc binding protein | Q8BHG2 | NO | NO | NO | YES | NO | NO | P |
| TB10B TBC1 domain family member 10B | Q8BHL3 | NO | NO | NO | YES | NO | NO | P |
| PSMF1 Proteasome inhibitor PI31 subunit | Q8BHL8 | NO | YES | NO | NO | NO | NO | P |
| GANAB Neutral alpha-glucosidase AB | Q8BHN3 | YES | NO | YES | YES | YES | YES | P/MQ |
| CEMIP Cell migration-inducing and hyaluronan-binding protein | Q8BI06 | NO | YES | NO | NO | NO | NO | P |
| MAL2 Protein MAL2 | Q8BI08 | NO | YES | NO | NO | NO | NO | P/MQ |
| CARF CDKN2A-interacting protein | Q8BI72 | NO | NO | NO | YES | YES | YES | P/MQ |
| TGO1 Transport and Golgi organization protein 1 homolog | Q8BI84 | NO | NO | NO | YES | NO | NO | P |
| SYIM Isoleucine--tRNA ligase, mitochondrial | Q8BIJ6 | NO | YES | NO | NO | NO | NO | P |
| CSTF2 Cleavage stimulation factor subunit 2 | Q8BIQ5 | NO | NO | NO | YES | NO | NO | P |
| CHDH Choline dehydrogenase, mitochondrial | Q8BJ64 | YES | NO | NO | NO | YES | NO | P/MQ |
| NUP93 Nuclear pore complex protein Nup93 | Q8BJ71 | NO | NO | NO | YES | NO | NO | P |
| CHM2B Charged multivesicular body protein 2b | Q8BJF9 | NO | NO | NO | YES | NO | NO | P |
| SMAL1 SWI/SNF-related matrix-associated actin-dependent regulator of chromatin subfamily A-like protein 1 | Q8BJL0 | NO | YES | NO | NO | NO | NO | P |

| | | | | | | | | |
|---|---|---|---|---|---|---|---|---|
| SUN2 SUN domain-containing protein 2 | Q8BJS4 | NO | NO | YES | YES | YES | YES | P |
| SGTA Small glutamine-rich tetratricopeptide repeat-containing protein alpha | Q8BJU0 | NO | NO | NO | NO | YES | NO | P |
| EIF2A Eukaryotic translation initiation factor 2A | Q8BJW6 | NO | NO | YES | YES | YES | YES | P/MQ |
| PSMD5 26S proteasome non-ATPase regulatory subunit 5 | Q8BJY1 | NO | YES | NO | NO | NO | NO | P |
| RT35 28S ribosomal protein S35, mitochondrial | Q8BJZ4 | NO | YES | NO | NO | NO | NO | P |
| NDUV3 NADH dehydrogenase [ubiquinone] flavoprotein 3, mitochondrial | Q8BK30 | YES | YES | NO | NO | NO | NO | P/MQ |
| AHSA1 Activator of 90 kDa heat shock protein ATPase homolog 1 | Q8BK64 | NO | NO | NO | NO | YES | NO | P |
| RCC2 Protein RCC2 | Q8BK67 | NO | NO | YES | YES | YES | YES | P/MQ |
| IPO5 Importin-5 | Q8BKC5 | NO | NO | NO | YES | NO | NO | P |
| BAIP2 Brain-specific angiogenesis inhibitor 1-associated protein 2 | Q8BKX1 | NO | YES | NO | NO | NO | NO | P |
| PDHX Pyruvate dehydrogenase protein X component, mitochondrial | Q8BKZ9 | NO | YES | NO | NO | NO | NO | P |
| EEA1 Early endosome antigen 1 | Q8BL66 | YES | NO | NO | YES | YES | YES | P/MQ |
| SRSF7 Serine/arginine-rich splicing factor 7 | Q8BL97 | NO | NO | YES | YES | YES | YES | P/MQ |
| NCEH1 Neutral cholesterol ester hydrolase 1 | Q8BLF1 | NO | NO | NO | NO | YES | NO | P |
| SRP68 Signal recognition particle subunit SRP68 | Q8BMA6 | NO | NO | NO | YES | YES | NO | P/MQ |
| VATG3 V-type proton ATPase subunit G 3 | Q8BMC1 | YES | NO | NO | NO | YES | YES | P/MQ |
| ODP2 Dihydrolipoyllysine-residue acetyltransferase component of pyruvate dehydrogenase complex, mitochondrial | Q8BMF4 | YES | NO | YES | YES | YES | NO | P/MQ |
| SYLC Leucine--tRNA ligase, cytoplasmic | Q8BMJ2 | NO | NO | NO | YES | NO | YES | P |
| IF1AX Eukaryotic translation initiation factor 1A, X-chromosomal | Q8BMJ3 | NO | NO | NO | YES | YES | NO | P |
| CKAP4 Cytoskeleton-associated protein 4 | Q8BMK4 | YES | NO | YES | YES | YES | YES | P/MQ |
| SYQ Glutamine--tRNA ligase | Q8BML9 | NO | NO | NO | YES | NO | YES | P |
| GCP60 Golgi resident protein GCP60 | Q8BMP6 | NO | NO | YES | YES | YES | YES | P/MQ |
| ECHA Trifunctional enzyme subunit alpha, mitochondrial | Q8BMS1 | YES | NO | YES | YES | YES | NO | P/MQ |
| SFR1 Swi5-dependent recombination DNA repair protein 1 homolog | Q8BP27 | NO | NO | NO | YES | NO | NO | P |
| PPA6 Lysophosphatidic acid phosphatase type 6 | Q8BP40 | YES | NO | NO | NO | NO | NO | P |
| SYNC Asparagine--tRNA ligase, cytoplasmic | Q8BP47 | NO | NO | NO | NO | YES | NO | P |
| RL24 60S ribosomal protein L24 | Q8BP67 | YES | NO | YES | YES | YES | YES | P/MQ |
| PPR18 Phostensin | Q8BQ30 | NO | NO | NO | YES | YES | YES | P |
| CPNE3 Copine-3 | Q8BT60 | YES | NO | YES | YES | YES | YES | P |
| Serine/arginine repetitive matrix protein 2 | Q8BTI8 | YES | YES | YES | YES | YES | YES | MQ |
| FLNA Filamin-A | Q8BTM8 | YES | NO | YES | YES | YES | YES | P/MQ |
| NUP54 Nuclear pore complex protein Nup54 | Q8BTS4 | NO | NO | NO | NO | YES | NO | P |

| | | | | | | | | |
|---|---|---|---|---|---|---|---|---|
| CPSF7 Cleavage and polyadenylation specificity factor subunit 7 | Q8BTV2 | NO | NO | NO | YES | NO | YES | P |
| GMPPB Mannose-1-phosphate guanyltransferase beta | Q8BTZ7 | NO | YES | NO | NO | NO | NO | P |
| SYIC Isoleucine--tRNA ligase, cytoplasmic | Q8BU30 | NO | NO | NO | NO | YES | NO | P |
| HOOK3 Protein Hook homolog 3 | Q8BUK6 | NO | YES | NO | NO | NO | NO | P |
| GEPH Gephyrin | Q8BUV3 | NO | YES | NO | NO | NO | NO | P |
| TIDC1 Complex I assembly factor TIMMDC1, mitochondrial | Q8BUY5 | NO | YES | NO | NO | NO | NO | P |
| IFIX Pyrin and HIN domain-containing protein 1 | Q8BV49 | NO | NO | YES | YES | YES | YES | P/MQ |
| VATH V-type proton ATPase subunit H | Q8BVE3 | YES | NO | NO | YES | YES | YES | P |
| DHPR Dihydropteridine reductase | Q8BVI4 | YES | NO | NO | NO | YES | NO | P/MQ |
| PPME1 Protein phosphatase methylesterase 1 | Q8BVQ5 | NO | NO | NO | YES | NO | NO | P |
| RL1D1 Ribosomal L1 domain-containing protein 1 | Q8BVY0 | NO | NO | NO | YES | YES | NO | P/MQ |
| SSDH Succinate-semialdehyde dehydrogenase, mitochondrial | Q8BWF0 | YES | NO | NO | NO | YES | NO | P |
| ACOT4 Acyl-coenzyme A thioesterase 4 | Q8BWN8 | NO | YES | NO | NO | NO | NO | P |
| CP062 UPF0505 protein C16orf62 homolog | Q8BWQ6 | NO | YES | NO | NO | NO | NO | P |
| OSGEP Probable tRNA N6-ade0sine threonylcarbamoyltransferase | Q8BWU5 | NO | YES | NO | NO | NO | NO | P |
| LARP4 La-related protein 4 | Q8BWW4 | NO | NO | NO | YES | NO | NO | P |
| ERF1 Eukaryotic peptide chain release factor subunit 1 | Q8BWY3 | NO | NO | YES | YES | NO | YES | P |
| KANK2 KN motif and ankyrin repeat domain-containing protein 2 | Q8BX02 | NO | YES | NO | NO | NO | NO | P |
| GOLI4 Golgi integral membrane protein 4 | Q8BXA1 | NO | NO | NO | NO | NO | YES | P |
| CLIC5 Chloride intracellular channel protein 5 | Q8BXK9 | NO | YES | NO | NO | NO | NO | P |
| UBP47 Ubiquitin carboxyl-terminal hydrolase 47 | Q8BY87 | YES | NO | NO | NO | YES | NO | P/MQ |
| TBCD Tubulin-specific chaperone D | Q8BYA0 | NO | YES | NO | NO | NO | NO | P |
| PCAT2 Lysophosphatidylcholine acyltransferase 2 | Q8BYI6 | NO | NO | NO | YES | NO | NO | P |
| YTHD3 YTH domain-containing family protein 3 | Q8BYK6 | NO | NO | NO | YES | NO | YES | P/MQ |
| RHG25 Rho GTPase-activating protein 25 | Q8BYW1 | NO | NO | NO | YES | NO | YES | P |
| SWAHB Ankyrin repeat domain-containing protein SOWAHB | Q8BZW2 | NO | YES | NO | NO | NO | NO | P |
| NHLC2 NHL repeat-containing protein 2 | Q8BZW8 | NO | YES | NO | NO | NO | NO | P |
| CHPT1 Cholinephosphotransferase 1 | Q8C025 | NO | YES | NO | NO | NO | NO | P |
| SYFA Phenylalanine--tRNA ligase alpha subunit | Q8C0C7 | NO | NO | NO | NO | YES | NO | P |
| VP26B Vacuolar protein sorting-associated protein 26B | Q8C0E2 | NO | NO | NO | NO | NO | YES | P |
| ADAS Alkyldihydroxyacetonephosphate synthase, peroxisomal | Q8C0I1 | NO | NO | NO | YES | NO | NO | P |
| TMX4 Thioredoxin-related transmembrane protein 4 | Q8C0L0 | NO | NO | NO | YES | YES | YES | P |
| LCAP Leucyl-cystinyl aminopeptidase | Q8C129 | NO | NO | NO | YES | NO | YES | P |
| DOCK8 Dedicator of cytokinesis protein 8 | Q8C147 | NO | NO | NO | YES | NO | NO | P |
| CPNE1 Copine-1 | Q8C166 | NO | NO | YES | YES | YES | YES | P |
| SEP11 Septin-11 | Q8C1B7 | YES | NO | YES | YES | YES | YES | P/MQ |

| | | | | | | | | |
|---|---|---|---|---|---|---|---|---|
| RBM14 RNA-binding protein 14 | Q8C2Q3 | NO | NO | NO | YES | YES | YES | P |
| DOCK2 Dedicator of cytokinesis protein 2 | Q8C3J5 | NO | NO | NO | YES | NO | YES | P/MQ |
| RAE1L mRNA export factor | Q8C570 | NO | YES | NO | NO | NO | NO | P |
| NAKD2 NAD kinase 2, mitochondrial | Q8C5H8 | YES | NO | NO | NO | NO | NO | P |
| C0T2 CCR4-0T transcription complex subunit 2 | Q8C5L3 | NO | YES | NO | NO | NO | NO | P |
| GRSF1 G-rich sequence factor 1 | Q8C5Q4 | NO | YES | NO | NO | NO | NO | P |
| CLMN Calmin | Q8C5W0 | NO | YES | NO | NO | NO | NO | P |
| UBA3 NEDD8-activating enzyme E1 catalytic subunit | Q8C878 | NO | YES | NO | NO | NO | NO | P |
| CAF17 Putative transferase CAF17 homolog, mitochondrial | Q8CAK1 | NO | YES | NO | NO | NO | NO | P |
| MIC60 MICOS complex subunit Mic60 | Q8CAQ8 | YES | NO | YES | YES | YES | YES | P/MQ |
| PARP9 Protein mono-ADP-ribosyltransferase PARP9 | Q8CAS9 | NO | YES | NO | NO | NO | NO | P |
| THIC Acetyl-CoA acetyltransferase, cytosolic | Q8CAY6 | YES | NO | NO | NO | NO | NO | P |
| ABI1 Abl interactor 1 | Q8CBW3 | NO | NO | NO | YES | NO | NO | P |
| GATC Glutamyl-tRNA(Gln) amidotransferase subunit C, mitochondrial | Q8CBY0 | NO | YES | NO | NO | NO | NO | P |
| SYNPO Synaptopodin | Q8CC35 | NO | YES | NO | NO | NO | NO | P/MQ |
| VWA8 von Willebrand factor A domain-containing protein 8 | Q8CC88 | YES | YES | NO | NO | NO | NO | P |
| PRP31 U4/U6 small nuclear ribonucleoprotein Prp31 | Q8CCF0 | NO | NO | NO | YES | NO | NO | P |
| RYBP RING1 and YY1-binding protein | Q8CCI5 | NO | NO | NO | YES | NO | NO | P |
| H2AW Core histone macro-H2A.2 | Q8CCK0 | NO | YES | NO | NO | NO | NO | P |
| NEMF Nuclear export mediator factor Nemf | Q8CCP0 | NO | YES | NO | NO | NO | NO | P |
| PABP2 Polyadenylate-binding protein 2 | Q8CCS6 | NO | NO | NO | YES | NO | NO | P/MQ |
| PIWL2 Piwi-like protein 2 | Q8CDG1 | NO | YES | NO | NO | NO | NO | P |
| TXNL1 Thioredoxin-like protein 1 | Q8CDN6 | YES | NO | YES | YES | YES | YES | P/MQ |
| NUP88 Nuclear pore complex protein Nup88 | Q8CEC0 | NO | YES | NO | NO | NO | NO | P |
| NED4L E3 ubiquitin-protein ligase NEDD4-like | Q8CFI0 | NO | YES | NO | NO | NO | NO | P |
| SYPM Probable proline--tRNA ligase, mitochondrial | Q8CFI5 | NO | YES | NO | NO | NO | NO | P |
| RPB2 DNA-directed RNA polymerase II subunit RPB2 | Q8CFI7 | NO | NO | NO | NO | YES | NO | P/MQ |
| G6PE GDH/6PGL endoplasmic bifunctional protein | Q8CFX1 | YES | NO | NO | YES | YES | YES | P/MQ |
| S22AC Solute carrier family 22 member 12 | Q8CFZ5 | NO | YES | NO | NO | NO | NO | P |
| ARK72 Aflatoxin B1 aldehyde reductase member 2 | Q8CG76 | YES | NO | YES | NO | YES | NO | P |
| SYEP Bifunctional glutamate/proline--tRNA ligase | Q8CGC7 | YES | NO | YES | YES | YES | YES | P/MQ |
| LONM Lon protease homolog, mitochondrial | Q8CGK3 | YES | NO | NO | YES | NO | NO | P |
| CCAR1 Cell division cycle and apoptosis regulator protein 1 | Q8CH18 | NO | YES | NO | NO | NO | NO | P |
| SLTM SAFB-like transcription modulator | Q8CH25 | NO | NO | NO | YES | NO | NO | P |
| SYNJ1 Synaptojanin-1 | Q8CHC4 | NO | NO | NO | YES | NO | NO | P |
| RFIP3 Rab11 family-interacting protein 3 | Q8CHD8 | NO | YES | NO | NO | NO | NO | P |
| PYM1 Partner of Y14 and mago | Q8CHP5 | NO | NO | NO | NO | YES | NO | P |
| PGP Glycerol-3-phosphate phosphatase | Q8CHP8 | NO | YES | NO | NO | NO | NO | P |

| | | | | | | | | |
|---|---|---|---|---|---|---|---|---|
| DPYD Dihydropyrimidine dehydrogenase [NADP(+)] | Q8CHR6 | NO | YES | NO | NO | NO | NO | P |
| PLCX1 PI-PLC X domain-containing protein 1 | Q8CHS4 | NO | YES | NO | NO | NO | NO | P |
| Delta-1-pyrroline-5-carboxylate dehydrogenase, mitochondrial | Q8CHT0 | YES | YES | YES | YES | YES | YES | MQ |
| P66A Transcriptional repressor p66 alpha | Q8CHY6 | NO | NO | NO | YES | NO | NO | P |
| PDLI5 PDZ and LIM domain protein 5 | Q8CI51 | YES | NO | YES | YES | YES | YES | P/MQ |
| PYGB Glycogen phosphorylase, brain form | Q8CI94 | YES | NO | NO | YES | NO | YES | P |
| FERM2 Fermitin family homolog 2 | Q8CIB5 | NO | YES | NO | NO | NO | NO | P |
| COPA Coatomer subunit alpha | Q8CIE6 | YES | NO | YES | YES | YES | YES | P/MQ |
| BTD Biotinidase | Q8CIF4 | NO | YES | NO | NO | NO | NO | P |
| SIDT2 SID1 transmembrane family member 2 | Q8CIF6 | NO | YES | NO | NO | NO | NO | P |
| PLCG2 1-phosphatidylinositol 4,5-bisphosphate phosphodiesterase gamma-2 | Q8CIH5 | NO | NO | NO | YES | YES | YES | P |
| PAK2 Serine/threonine-protein kinase PAK 2 | Q8CIN4 | YES | NO | YES | YES | YES | YES | P/MQ |
| ASPM Abnormal spindle-like microcephaly-associated protein homolog | Q8CJ27 | NO | YES | NO | NO | NO | NO | P |
| CROCC Rootletin | Q8CJ40 | NO | YES | NO | NO | NO | NO | P |
| ELYS Protein ELYS | Q8CJF7 | NO | NO | NO | YES | NO | NO | P |
| AGO3 Protein argonaute-3 | Q8CJF9 | NO | YES | NO | NO | NO | NO | P |
| HMGCS1 Hydroxymethylglutaryl-CoA synthase, cytoplasmic | Q8JZK9 | NO | YES | NO | NO | NO | NO | P |
| AFG32 AFG3-like protein 2 | Q8JZQ2 | NO | YES | NO | NO | NO | NO | P |
| EIF3B Eukaryotic translation initiation factor 3 subunit B | Q8JZQ9 | YES | NO | YES | YES | YES | YES | P/MQ |
| NAGA N-acetylglucosamine-6-phosphate deacetylase | Q8JZV7 | NO | NO | NO | YES | NO | NO | P |
| BDH2 3-hydroxybutyrate dehydrogenase type 2 | Q8JZV9 | YES | NO | YES | NO | YES | NO | P |
| Splicing factor 45 | Q8JZX4 | NO | YES | NO | YES | NO | YES | MQ |
| UD3A2 UDP-glucuronosyltransferase 3A2 | Q8JZZ0 | NO | YES | NO | NO | NO | NO | P |
| TMA7 Translation machinery-associated protein 7 | Q8K003 | NO | NO | NO | YES | NO | NO | P |
| OPLA 5-oxoprolinase | Q8K010 | YES | NO | NO | NO | YES | NO | P |
| BCLF1 Bcl-2-associated transcription factor 1 | Q8K019 | NO | NO | NO | YES | NO | NO | P/MQ |
| SCAM1 Secretory carrier-associated membrane protein 1 | Q8K021 | NO | NO | NO | YES | YES | YES | P |
| EFGM Elongation factor G, mitochondrial | Q8K0D5 | NO | YES | NO | NO | NO | NO | P |
| FIBB Fibrinogen beta chain | Q8K0E8 | YES | NO | YES | YES | YES | YES | P/MQ |
| ACSM2 Acyl-coenzyme A synthetase ACSM2, mitochondrial | Q8K0L3 | YES | NO | YES | NO | YES | NO | P/MQ |
| HS12A Heat shock 70 kDa protein 12A | Q8K0U4 | NO | YES | NO | NO | NO | NO | P/MQ |
| PKHO2 Pleckstrin homology domain-containing family O member 2 | Q8K124 | NO | NO | YES | YES | YES | YES | P/MQ |
| GALM Aldose 1-epimerase | Q8K157 | YES | YES | NO | NO | NO | NO | P/MQ |
| PDXK Pyridoxal kinase | Q8K183 | NO | NO | YES | NO | YES | NO | P |
| URP2 Fermitin family homolog 3 | Q8K1B8 | NO | NO | YES | YES | YES | YES | P |

| | | | | | | | | |
|---|---|---|---|---|---|---|---|---|
| STX5 Syntaxin-5 | Q8K1E0 | NO | NO | NO | YES | NO | NO | P |
| WIPF1 WAS/WASL-interacting protein family member 1 | Q8K1I7 | NO | NO | NO | YES | YES | YES | P |
| DNM1L Dynamin-1-like protein | Q8K1M6 | YES | NO | YES | YES | YES | YES | P/MQ |
| SPAS2 Spermatogenesis-associated serine-rich protein 2 | Q8K1N4 | NO | NO | NO | YES | NO | NO | P |
| WDR11 WD repeat-containing protein 11 | Q8K1X1 | NO | YES | NO | NO | NO | NO | P |
| COQ9 Ubiquinone biosynthesis protein COQ9, mitochondrial | Q8K1Z0 | YES | YES | NO | NO | NO | NO | P |
| ARFP2 Arfaptin-2 | Q8K221 | NO | YES | NO | NO | NO | NO | P |
| GT251 Procollagen galactosyltransferase 1 | Q8K297 | NO | NO | NO | YES | YES | YES | P |
| SDHA Succinate dehydrogenase [ubiquinone] flavoprotein subunit, mitochondrial | Q8K2B3 | YES | NO | YES | YES | YES | YES | P/MQ |
| OTU6B Deubiquitinase OTUD6B | Q8K2H2 | NO | YES | NO | NO | NO | NO | P |
| MANBA Beta-mannosidase | Q8K2I4 | NO | YES | NO | NO | NO | NO | P |
| AGFG1 Arf-GAP domain and FG repeat-containing protein 1 | Q8K2K6 | NO | NO | NO | YES | NO | NO | P |
| SHOT1 Shootin-1 | Q8K2Q9 | NO | NO | NO | NO | NO | YES | P |
| PP4R1 Serine/threonine-protein phosphatase 4 regulatory subunit 1 | Q8K2V1 | NO | NO | NO | YES | NO | NO | P |
| EVA1B Protein eva-1 homolog B | Q8K2Y3 | NO | YES | NO | NO | NO | NO | P/MQ |
| MATR3 Matrin-3 | Q8K310 | YES | NO | YES | YES | YES | YES | P/MQ |
| SASH3 SAM and SH3 domain-containing protein 3 | Q8K352 | NO | NO | NO | YES | NO | NO | P |
| CBR3 Carbonyl reductase [NADPH] 3 | Q8K354 | NO | NO | NO | YES | NO | NO | P/MQ |
| DCC-interacting protein 13-alpha | Q8K3H0 | NO | YES | YES | YES | NO | YES | MQ |
| NDUS8 NADH dehydrogenase [ubiquinone] iron-sulfur protein 8, mitochondrial | Q8K3J1 | NO | YES | NO | NO | NO | NO | P |
| GPC5C G-protein coupled receptor family C group 5 member C | Q8K3J9 | NO | YES | NO | NO | NO | NO | P |
| PREP Presequence protease, mitochondrial | Q8K411 | NO | YES | NO | NO | NO | NO | P |
| EMIL2 EMILIN-2 | Q8K482 | NO | NO | NO | YES | NO | NO | P |
| IRAK3 Interleukin-1 receptor-associated kinase 3 | Q8K4B2 | NO | NO | NO | NO | YES | NO | P |
| ABLM1 Actin-binding LIM protein 1 | Q8K4G5 | NO | YES | NO | NO | NO | NO | P |
| ARHG6 Rho guanine nucleotide exchange factor 6 | Q8K4I3 | NO | NO | NO | YES | NO | NO | P/MQ |
| PANK1 Pantothenate kinase 1 | Q8K4K6 | NO | YES | NO | NO | NO | NO | P |
| SVIL Supervillin | Q8K4L3 | NO | NO | YES | YES | YES | YES | P/MQ |
| NNRE NAD(P)H-hydrate epimerase | Q8K4Z3 | NO | YES | NO | NO | NO | NO | P |
| SF3A1 Splicing factor 3A subunit 1 | Q8K4Z5 | YES | NO | YES | YES | YES | YES | P/MQ |
| HIBCH 3-hydroxyisobutyryl-CoA hydrolase, mitochondrial | Q8QZS1 | NO | YES | NO | NO | NO | NO | P |
| THIL Acetyl-CoA acetyltransferase, mitochondrial | Q8QZT1 | YES | NO | YES | YES | YES | YES | P/MQ |
| F151A Protein FAM151A | Q8QZW3 | NO | YES | NO | NO | NO | NO | P |
| EIF3L Eukaryotic translation initiation factor 3 subunit L | Q8QZY1 | NO | NO | NO | YES | YES | YES | P/MQ |

| | | | | | | | | |
|---|---|---|---|---|---|---|---|---|
| GLCTK Glycerate kinase | Q8QZY2 | NO | YES | NO | NO | NO | NO | P |
| SF3B4 Splicing factor 3B subunit 4 | Q8QZY9 | NO | YES | NO | NO | NO | NO | P/MQ |
| MARE2 Microtubule-associated protein RP/EB family member 2 | Q8R001 | NO | NO | NO | YES | NO | NO | P |
| AIMP2 Aminoacyl tRNA synthase complex-interacting multifunctional protein 2 | Q8R010 | NO | YES | NO | NO | NO | NO | P |
| ICT1 Peptidyl-tRNA hydrolase ICT1, mitochondrial | Q8R035 | NO | YES | NO | NO | NO | NO | P |
| ERF3A Eukaryotic peptide chain release factor GTP-binding subunit ERF3A | Q8R050 | NO | NO | NO | YES | NO | NO | P |
| HNRPL Heterogeneous nuclear ribonucleoprotein L | Q8R081 | NO | NO | YES | YES | YES | YES | P/MQ |
| SUOX Sulfite oxidase, mitochondrial | Q8R086 | NO | YES | NO | NO | NO | NO | P |
| General transcription factor IIF subunit 2 | Q8R0A0 | YES | YES | YES | YES | YES | YES | MQ |
| FAHD1 Acylpyruvase FAHD1, mitochondrial | Q8R0F8 | NO | YES | NO | NO | NO | NO | P |
| GGA1 ADP-ribosylation factor-binding protein GGA1 | Q8R0H9 | NO | NO | NO | NO | YES | NO | P |
| HOT Hydroxyacid-oxoacid transhydrogenase, mitochondrial | Q8R0N6 | NO | YES | NO | NO | NO | NO | P |
| SGPL1 Sphingosine-1-phosphate lyase 1 | Q8R0X7 | NO | NO | NO | YES | YES | YES | P |
| APEH Acylamino-acid-releasing enzyme | Q8R146 | YES | NO | NO | YES | NO | NO | P |
| BPHL Valacyclovir hydrolase | Q8R164 | YES | NO | NO | YES | YES | NO | P/MQ |
| ERO1A ERO1-like protein alpha | Q8R180 | NO | NO | NO | YES | YES | NO | P |
| DOCK7 Dedicator of cytokinesis protein 7 | Q8R1A4 | NO | YES | NO | NO | NO | NO | P |
| EIF3C Eukaryotic translation initiation factor 3 subunit C | Q8R1B4 | YES | NO | YES | YES | YES | YES | P/MQ |
| PDLI2 PDZ and LIM domain protein 2 | Q8R1G6 | NO | NO | NO | YES | YES | NO | P |
| QCR9 Cytochrome b-c1 complex subunit 9 | Q8R1I1 | YES | YES | NO | NO | NO | NO | P/MQ |
| H2AJ Histone H2A.J | Q8R1M2 | YES | NO | YES | YES | YES | YES | P |
| DC1L1 Cytoplasmic dynein 1 light intermediate chain 1 | Q8R1Q8 | YES | NO | YES | YES | YES | YES | P/MQ |
| CGRE1 Cell growth regulator with EF hand domain protein 1 | Q8R1U2 | NO | YES | NO | NO | NO | NO | P |
| TMED4 Transmembrane emp24 domain-containing protein 4 | Q8R1V4 | NO | NO | NO | YES | NO | NO | P |
| CD177 CD177 antigen | Q8R2S8 | NO | NO | YES | YES | YES | YES | P/MQ |
| NUDT4 Diphosphoinositol polyphosphate phosphohydrolase 2 | Q8R2U6 | NO | YES | NO | NO | NO | NO | P |
| UBQL1 Ubiquilin-1 | Q8R317 | NO | NO | NO | YES | YES | YES | P/MQ |
| RFIP5 Rab11 family-interacting protein 5 | Q8R361 | YES | NO | YES | YES | YES | NO | P/MQ |
| ACY2 Aspartoacylase | Q8R3P0 | NO | YES | NO | NO | NO | NO | P |
| CCD58 Coiled-coil domain-containing protein 58 | Q8R3Q6 | NO | YES | NO | NO | NO | NO | P |
| CS047 Uncharacterized protein C19orf47 homolog | Q8R3Y5 | NO | YES | NO | NO | NO | NO | P |
| HEXI1 Protein HEXIM1 | Q8R409 | NO | NO | NO | YES | NO | NO | P |
| ABCA3 ATP-binding cassette sub-family A member 3 | Q8R420 | NO | YES | NO | NO | NO | NO | P |
| IL36G Interleukin-36 gamma | Q8R460 | NO | NO | NO | YES | YES | YES | P |
| IRAK4 Interleukin-1 receptor-associated kinase 4 | Q8R4K2 | NO | NO | NO | YES | NO | NO | P |

| | | | | | | | | |
|---|---|---|---|---|---|---|---|---|
| CLYBL Citramalyl-CoA lyase, mitochondrial | Q8R4N0 | YES | NO | YES | YES | YES | NO | P |
| NUP35 Nucleoporin NUP35 | Q8R4R6 | NO | NO | NO | YES | NO | YES | P |
| LUZP1 Leucine zipper protein 1 | Q8R4U7 | NO | NO | NO | YES | NO | NO | P |
| SH3K1 SH3 domain-containing kinase-binding protein 1 | Q8R550 | NO | NO | NO | YES | YES | YES | P/MQ |
| AB1IP Amyloid beta A4 precursor protein-binding family B member 1-interacting protein | Q8R5A3 | NO | NO | NO | YES | NO | NO | P |
| ACTY Beta-centractin | Q8R5C5 | NO | YES | NO | NO | NO | NO | P |
| TMX1 Thioredoxin-related transmembrane protein 1 | Q8VBT0 | NO | NO | NO | NO | YES | NO | P |
| APOBR Apolipoprotein B receptor | Q8VBT6 | NO | NO | YES | YES | YES | YES | P/MQ |
| ASPC1 Tether containing UBX domain for GLUT4 | Q8VBT9 | NO | NO | NO | YES | NO | NO | P |
| CSN8 COP9 signalosome complex subunit 8 | Q8VBV7 | NO | NO | NO | NO | YES | NO | P |
| Cleft lip and palate transmembrane protein 1 homolog | Q8VBZ3 | YES | YES | YES | YES | YES | YES | MQ |
| HUTU Urocanate hydratase | Q8VC12 | NO | YES | NO | NO | NO | NO | P |
| TKFC Triokinase/FMN cyclase | Q8VC30 | YES | NO | YES | YES | YES | NO | P/MQ |
| S22A6 Solute carrier family 22 member 6 | Q8VC69 | NO | YES | NO | NO | NO | NO | P |
| SCRN2 Secernin-2 | Q8VCA8 | NO | YES | NO | NO | NO | NO | P |
| RPGF3 Rap guanine nucleotide exchange factor 3 | Q8VCC8 | NO | YES | NO | NO | NO | NO | P |
| MAVS Mitochondrial antiviral-signaling protein | Q8VCF0 | NO | YES | NO | NO | NO | NO | P |
| FIBG Fibrinogen gamma chain | Q8VCM7 | YES | NO | YES | YES | YES | YES | P/MQ |
| CGL Cystathionine gamma-lyase | Q8VCN5 | YES | NO | NO | NO | YES | NO | P |
| TBCC Tubulin-specific chaperone C | Q8VCN9 | NO | YES | NO | NO | NO | NO | P |
| Caldesmon 1 | Q8VCQ8 | YES | YES | YES | YES | YES | YES | MQ |
| ABHEB Protein ABHD14B | Q8VCR7 | YES | NO | YES | NO | YES | NO | P |
| AMPB Aminopeptidase B | Q8VCT3 | NO | NO | NO | YES | YES | YES | P/MQ |
| CES1D Carboxylesterase 1D | Q8VCT4 | YES | YES | NO | NO | NO | NO | P |
| HIP1 Huntingtin-interacting protein 1 | Q8VD75 | NO | NO | NO | YES | NO | NO | P |
| MYH9 Myosin-9 | Q8VDD5 | YES | NO | YES | YES | YES | YES | P/MQ |
| VIGLN Vigilin | Q8VDJ3 | YES | NO | YES | YES | YES | YES | P/MQ |
| ADPGK ADP-dependent glucokinase | Q8VDL4 | NO | NO | NO | YES | NO | YES | P |
| PSMD2 26S proteasome non-ATPase regulatory subunit 2 | Q8VDM4 | NO | NO | YES | YES | YES | YES | P/MQ |
| HNRL1 Heterogeneous nuclear ribonucleoprotein U-like protein 1 | Q8VDM6 | NO | NO | NO | YES | NO | NO | P |
| AT1A1 Sodium/potassium-transporting ATPase subunit alpha-1 | Q8VDN2 | YES | NO | YES | YES | YES | YES | P/MQ |
| MICA1 [F-actin]-monooxygenase MICAL1 | Q8VDP3 | NO | NO | NO | YES | NO | NO | P |
| CCAR2 Cell cycle and apoptosis regulator protein 2 | Q8VDP4 | NO | NO | NO | YES | NO | YES | P |
| SIR2 NAD-dependent protein deacetylase sirtuin-2 | Q8VDQ8 | NO | NO | NO | YES | YES | NO | P |
| RCC1 Regulator of chromosome condensation | Q8VE37 | YES | NO | YES | YES | YES | YES | P/MQ |
| PDC10 Programmed cell death protein 10 | Q8VE70 | NO | YES | NO | NO | NO | NO | P |
| SRSF4 Serine/arginine-rich splicing factor 4 | Q8VE97 | YES | NO | NO | YES | YES | YES | P/MQ |

| | | | | | | | | |
|---|---|---|---|---|---|---|---|---|
| LMCD1 LIM and cysteine-rich domains protein 1 | Q8VEE1 | NO | YES | NO | NO | NO | NO | P |
| VPS4A Vacuolar protein sorting-associated protein 4A | Q8VEJ9 | NO | YES | NO | NO | NO | NO | P |
| HNRPU Heterogeneous nuclear ribonucleoprotein U | Q8VEK3 | YES | NO | YES | YES | YES | YES | P/MQ |
| MPCP Phosphate carrier protein, mitochondrial | Q8VEM8 | YES | NO | YES | NO | YES | YES | P |
| RBM39 RNA-binding protein 39 | Q8VH51 | NO | NO | YES | YES | YES | YES | P/MQ |
| SEC63 Translocation protein SEC63 homolog | Q8VHE0 | NO | NO | NO | NO | YES | NO | P |
| CARTF Calcium-responsive transcription factor | Q8VHI4 | NO | YES | NO | NO | NO | NO | P |
| P66B Transcriptional repressor p66-beta | Q8VHR5 | NO | NO | NO | YES | NO | NO | P |
| PAXI Paxillin | Q8VI36 | NO | NO | NO | YES | YES | NO | P |
| SFPQ Splicing factor, proline- and glutamine-rich | Q8VIJ6 | YES | NO | YES | YES | YES | YES | P/MQ |
| BSND Barttin | Q8VIM4 | NO | YES | NO | NO | NO | NO | P |
| TRAM1 Translocating chain-associated membrane protein 1 | Q91V04 | NO | NO | NO | YES | NO | NO | P |
| RAB14 Ras-related protein Rab-14 | Q91V41 | YES | NO | YES | YES | YES | YES | P/MQ |
| SFXN3 Sideroflexin-3 | Q91V61 | YES | NO | NO | YES | YES | NO | P/MQ |
| ISOC1 Isochorismatase domain-containing protein 1 | Q91V64 | YES | NO | NO | YES | YES | YES | P/MQ |
| CK054 Ester hydrolase C11orf54 homolog | Q91V76 | YES | NO | YES | NO | YES | NO | P |
| ACLY ATP-citrate synthase | Q91V92 | NO | NO | YES | YES | YES | YES | P/MQ |
| ACSM1 Acyl-coenzyme A synthetase ACSM1, mitochondrial | Q91VA0 | NO | YES | NO | NO | NO | NO | P |
| PDIP2 Polymerase delta-interacting protein 2 | Q91VA6 | NO | YES | NO | NO | NO | NO | P |
| IF4A3 Eukaryotic initiation factor 4A-III | Q91VC3 | NO | NO | NO | YES | YES | NO | P |
| PLVAP Plasmalemma vesicle-associated protein | Q91VC4 | YES | YES | NO | NO | NO | NO | P |
| NDUS1 NADH-ubiquinone oxidoreductase 75 kDa subunit, mitochondrial | Q91VD9 | YES | NO | YES | YES | YES | YES | P/MQ |
| SNX9 Sorting nexin-9 | Q91VH2 | NO | NO | NO | YES | YES | NO | P |
| RINI Ribonuclease inhibitor | Q91VI7 | YES | NO | YES | YES | YES | YES | P/MQ |
| PQBP1 Polyglutamine-binding protein 1 | Q91VJ5 | NO | NO | NO | YES | NO | NO | P |
| IPYR2 Inorganic pyrophosphatase 2, mitochondrial | Q91VM9 | YES | NO | NO | NO | YES | NO | P/MQ |
| MIC25 MICOS complex subunit Mic25 | Q91VN4 | NO | YES | NO | NO | NO | NO | P |
| ATPG ATP synthase subunit gamma, mitochondrial | Q91VR2 | YES | NO | YES | YES | YES | NO | P/MQ |
| DDX1 ATP-dependent RNA helicase DDX1 | Q91VR5 | YES | NO | YES | YES | YES | NO | P/MQ |
| CBR4 Carbonyl reductase family member 4 | Q91VT4 | NO | YES | NO | NO | NO | NO | P |
| SH3L3 SH3 domain-binding glutamic acid-rich-like protein 3 | Q91VW3 | NO | NO | NO | YES | YES | YES | P/MQ |
| UBAP2 Ubiquitin-associated protein 2 | Q91VX2 | NO | YES | NO | NO | NO | NO | P |
| GCSP Glycine dehydrogenase (decarboxylating), mitochondrial | Q91W43 | NO | YES | NO | NO | NO | NO | P |
| CSDE1 Cold shock domain-containing protein E1 | Q91W50 | NO | NO | YES | YES | YES | NO | P/MQ |
| RBMS1 RNA-binding motif, single-stranded-interacting protein 1 | Q91W59 | NO | NO | NO | YES | NO | NO | P |
| NRK1 Nicotinamide riboside kinase 1 | Q91W63 | NO | YES | NO | NO | NO | NO | P |
| TXND5 Thioredoxin domain-containing protein 5 | Q91W90 | NO | NO | YES | YES | YES | YES | P |

| | | | | | | | | |
|---|---|---|---|---|---|---|---|---|
| SETD3 Actin-histidine N-methyltransferase | Q91WC0 | NO | NO | NO | NO | YES | NO | P |
| NDUS2 NADH dehydrogenase [ubiquinone] iron-sulfur protein 2, mitochondrial | Q91WD5 | YES | YES | NO | NO | NO | NO | P |
| CES2C Acylcarnitine hydrolase | Q91WG0 | YES | YES | NO | NO | NO | NO | P |
| RABE2 Rab GTPase-binding effector protein 2 | Q91WG2 | NO | YES | NO | NO | NO | NO | P |
| FUBP1 Far upstream element-binding protein 1 | Q91WJ8 | YES | NO | YES | YES | YES | YES | P/MQ |
| LRRF2 Leucine-rich repeat flightless-interacting protein 2 | Q91WK0 | YES | NO | NO | NO | YES | NO | P/MQ |
| EIF3H Eukaryotic translation initiation factor 3 subunit H | Q91WK2 | NO | NO | NO | YES | YES | NO | P/MQ |
| EVA1A Protein eva-1 homolog A | Q91WM6 | NO | YES | NO | NO | NO | NO | P |
| KMO Kynurenine 3-monooxygenase | Q91WN4 | NO | YES | NO | NO | NO | NO | P |
| SPA3N Serine protease inhibitor A3N | Q91WP6 | YES | NO | YES | YES | YES | YES | P/MQ |
| SYYC Tyrosine--tRNA ligase, cytoplasmic | Q91WQ3 | NO | NO | YES | YES | YES | NO | P |
| AK1CL Aldo-keto reductase family 1 member C21 | Q91WR5 | YES | NO | NO | NO | YES | NO | P |
| CISD1 CDGSH iron-sulfur domain-containing protein 1 | Q91WS0 | YES | YES | NO | NO | NO | NO | P/MQ |
| RBM47 RNA-binding protein 47 | Q91WT8 | NO | YES | NO | NO | NO | NO | P |
| AS3MT Arsenite methyltransferase | Q91WU5 | YES | YES | NO | NO | NO | NO | P |
| NC2B Protein Dr1 | Q91WV0 | NO | NO | YES | NO | NO | NO | P |
| PODO Podocin | Q91X05 | NO | YES | NO | NO | NO | NO | P |
| UROM Uromodulin | Q91X17 | YES | YES | NO | NO | NO | NO | P |
| DCXR L-xylulose reductase | Q91X52 | NO | NO | NO | NO | YES | NO | P |
| HPX Hemopexin | Q91X72 | YES | NO | YES | YES | YES | YES | P/MQ |
| ERLN1 Erlin-1 | Q91X78 | NO | YES | NO | NO | NO | NO | P |
| GOLM1 Golgi membrane protein 1 | Q91XA2 | NO | NO | NO | YES | NO | NO | P |
| DAP1 Death-associated protein 1 | Q91XC8 | NO | NO | YES | YES | NO | NO | P |
| VPS36 Vacuolar protein-sorting-associated protein 36 | Q91XD6 | NO | YES | NO | NO | NO | NO | P |
| GLYAT Glycine N-acyltransferase | Q91XE0 | YES | NO | NO | NO | YES | NO | P |
| ACY3 N-acyl-aromatic-L-amino acid amidohydrolase (carboxylate-forming) | Q91XE4 | YES | NO | YES | NO | YES | NO | P |
| PNPO Pyridoxine-5'-phosphate oxidase | Q91XF0 | NO | YES | NO | NO | NO | NO | P |
| Leucine-rich HEV glycoprotein | Q91XL1 | YES | YES | YES | YES | YES | YES | MQ |
| OSBL1 Oxysterol-binding protein-related protein 1 | Q91XL9 | NO | YES | NO | NO | NO | NO | P |
| BASP1 Brain acid soluble protein 1 | Q91XV3 | NO | NO | YES | YES | YES | YES | P/MQ |
| S13A3 Solute carrier family 13 member 3 | Q91Y63 | NO | YES | NO | NO | NO | NO | P |
| ARLY Argininosuccinate lyase | Q91YI0 | YES | NO | YES | YES | YES | NO | P/MQ |
| ARRB2 Beta-arrestin-2 | Q91YI4 | NO | NO | NO | YES | YES | NO | P |
| SNX4 Sorting nexin-4 | Q91YJ2 | NO | YES | NO | NO | NO | NO | P/MQ |
| UAP1 UDP-N-acetylhexosamine pyrophosphorylase | Q91YN5 | NO | NO | NO | NO | YES | NO | P |
| L2HDH L-2-hydroxyglutarate dehydrogenase, mitochondrial | Q91YP0 | NO | YES | NO | NO | NO | NO | P |
| NEUL Neurolysin, mitochondrial | Q91YP2 | NO | YES | NO | NO | NO | NO | P |

| | | | | | | | | |
|---|---|---|---|---|---|---|---|---|
| RPN1 Dolichyl-diphosphooligosaccharide--protein glycosyltransferase subunit 1 | Q91YQ5 | YES | NO | YES | YES | YES | YES | P/MQ |
| TWF1 Twinfilin-1 | Q91YR1 | YES | NO | YES | YES | YES | YES | P/MQ |
| PRP6 Pre-mRNA-processing factor 6 | Q91YR7 | NO | YES | NO | NO | NO | NO | P |
| PTGR1 Prostaglandin reductase 1 | Q91YR9 | NO | NO | NO | YES | NO | NO | P |
| Kinesin light chain 2 | Q91YS4 | YES | YES | YES | YES | YES | YES | MQ |
| NDUV1 NADH dehydrogenase [ubiquinone] flavoprotein 1, mitochondrial | Q91YT0 | YES | YES | NO | NO | NO | NO | P |
| YTH domain-containing family protein 2 | Q91YT7 | YES | YES | YES | YES | NO | YES | MQ |
| DNJC3 DnaJ homolog subfamily C member 3 | Q91YW3 | NO | NO | NO | YES | NO | NO | P |
| GRHPR Glyoxylate reductase/hydroxypyruvate reductase | Q91Z53 | YES | NO | YES | NO | YES | YES | P/MQ |
| BMP2K BMP-2-inducible protein kinase | Q91Z96 | NO | YES | NO | NO | NO | NO | P |
| PCCA Propionyl-CoA carboxylase alpha chain, mitochondrial | Q91ZA3 | YES | NO | YES | NO | YES | NO | P/MQ |
| UGPA UTP--glucose-1-phosphate uridylyltransferase | Q91ZJ5 | YES | NO | YES | YES | YES | YES | P/MQ |
| LPIN1 Phosphatidate phosphatase LPIN1 | Q91ZP3 | NO | YES | NO | NO | NO | NO | P |
| SNX18 Sorting nexin-18 | Q91ZR2 | NO | NO | NO | YES | NO | YES | P |
| MIA2 Melanoma inhibitory activity protein 2 | Q91ZV0 | NO | NO | NO | YES | NO | NO | P |
| SMCA5 SWI/SNF-related matrix-associated actin-dependent regulator of chromatin subfamily A member 5 | Q91ZW3 | NO | NO | YES | YES | YES | YES | P/MQ |
| LRP1 Prolow-density lipoprotein receptor-related protein 1 | Q91ZX7 | NO | NO | YES | YES | YES | YES | P/MQ |
| RISC Retinoid-inducible serine carboxypeptidase | Q920A5 | NO | YES | NO | NO | NO | NO | P |
| SP16H FACT complex subunit SPT16 | Q920B9 | NO | NO | NO | YES | YES | YES | P/MQ |
| FPPS Farnesyl pyrophosphate synthase | Q920E5 | NO | NO | NO | YES | NO | NO | P |
| VPP4 V-type proton ATPase 116 kDa subunit a isoform 4 | Q920R6 | YES | YES | NO | NO | NO | NO | P |
| RAB31 Ras-related protein Rab-31 | Q921E2 | NO | NO | NO | YES | YES | NO | P |
| TADBP TAR DNA-binding protein 43 | Q921F2 | YES | NO | YES | YES | YES | YES | P/MQ |
| HNRLL Heterogeneous nuclear ribonucleoprotein L-like | Q921F4 | NO | NO | NO | YES | NO | NO | P |
| LRCH4 Leucine-rich repeat and calponin homology domain-containing protein 4 | Q921G6 | NO | NO | NO | YES | YES | NO | P |
| ETFD Electron transfer flavoprotein-ubiquinone oxidoreductase, mitochondrial | Q921G7 | YES | NO | YES | NO | YES | NO | P/MQ |
| THIKA 3-ketoacyl-CoA thiolase A, peroxisomal | Q921H8 | YES | NO | YES | YES | YES | YES | P/MQ |
| TRFE Serotransferrin | Q921I1 | YES | NO | YES | YES | YES | YES | P/MQ |
| TMCO1 Calcium load-activated calcium channel | Q921L3 | NO | NO | NO | NO | YES | NO | P |
| SF3B3 Splicing factor 3B subunit 3 | Q921M3 | NO | NO | YES | YES | YES | YES | P/MQ |
| FA49B Protein FAM49B | Q921M7 | NO | NO | NO | YES | YES | YES | P/MQ |
| RM37 39S ribosomal protein L37, mitochondrial | Q921S7 | NO | YES | NO | NO | NO | NO | P |
| TOIP1 Torsin-1A-interacting protein 1 | Q921T2 | NO | YES | NO | NO | NO | NO | P |
| SMTN Smoothelin | Q921U8 | YES | NO | NO | NO | NO | NO | P |

| | | | | | | | | |
|---|---|---|---|---|---|---|---|---|
| MOB1A MOB kinase activator 1A | Q921Y0 | YES | NO | NO | YES | NO | YES | P |
| TFIP8 Tumor necrosis factor alpha-induced protein 8 | Q921Z5 | NO | YES | NO | NO | NO | NO | P |
| SYDC Aspartate--tRNA ligase, cytoplasmic | Q922B2 | YES | NO | YES | YES | YES | YES | P/MQ |
| ITPI2 Protein ITPRID2 | Q922B9 | NO | YES | NO | NO | NO | NO | P |
| C1TC C-1-tetrahydrofolate synthase, cytoplasmic | Q922D8 | YES | NO | YES | YES | YES | YES | P/MQ |
| PCY2 Ethanolamine-phosphate cytidylyltransferase | Q922E4 | NO | YES | NO | NO | NO | NO | P |
| GMPPA Mannose-1-phosphate guanyltransferase alpha | Q922H4 | NO | NO | NO | NO | YES | NO | P |
| CLIP1 CAP-Gly domain-containing linker protein 1 | Q922J3 | NO | NO | YES | YES | YES | YES | P/MQ |
| GLYR1 Putative oxidoreductase GLYR1 | Q922P9 | NO | NO | NO | NO | YES | NO | P |
| MARC2 Mitochondrial amidoxime reducing component 2 | Q922Q1 | YES | NO | NO | NO | NO | NO | P |
| LRC59 Leucine-rich repeat-containing protein 59 | Q922Q8 | NO | NO | YES | YES | YES | NO | P/MQ |
| PDIA6 Protein disulfide-isomerase A6 | Q922R8 | YES | NO | YES | YES | YES | YES | P/MQ |
| UBXN1 UBX domain-containing protein 1 | Q922Y1 | NO | NO | NO | YES | YES | YES | P |
| BLVRB Flavin reductase (NADPH) | Q923D2 | NO | NO | NO | YES | YES | YES | P/MQ |
| WBP11 WW domain-binding protein 11 | Q923D5 | NO | NO | NO | YES | NO | NO | P |
| SC5A2 Sodium/glucose cotransporter 2 | Q923I7 | NO | YES | NO | NO | NO | NO | P |
| ST5 Suppression of tumorigenicity 5 protein | Q924W7 | NO | NO | NO | YES | NO | NO | P |
| PAWR PRKC apoptosis WT1 regulator protein | Q925B0 | YES | NO | YES | YES | YES | YES | P/MQ |
| ATAD3 ATPase family AAA domain-containing protein 3 | Q925I1 | YES | NO | NO | NO | NO | NO | P |
| TALDO1 Transaldolase | Q93092 | YES | NO | YES | YES | YES | YES | P/MQ |
| ROAA Heterogeneous nuclear ribonucleoprotein A/B | Q99020 | NO | NO | YES | YES | YES | YES | P/MQ |
| MLYCD Malonyl-CoA decarboxylase, mitochondrial | Q99J39 | NO | YES | NO | NO | NO | NO | P |
| SIAS Sialic acid synthase | Q99J77 | NO | NO | YES | YES | YES | YES | P/MQ |
| THTM 3-mercaptopyruvate sulfurtransferase | Q99J99 | YES | NO | NO | YES | NO | NO | P |
| STML2 Stomatin-like protein 2, mitochondrial | Q99JB2 | NO | NO | NO | YES | YES | NO | P |
| AMNLS Protein amnionless | Q99JB7 | NO | YES | NO | NO | NO | NO | P |
| PSIP1 PC4 and SFRS1-interacting protein | Q99JF8 | NO | NO | NO | YES | NO | NO | P |
| Musculoskeletal embryonic nuclear protein 1 | Q99JI1 | YES | YES | YES | YES | YES | YES | MQ |
| PSMD6 26S proteasome non-ATPase regulatory subunit 6 | Q99JI4 | YES | NO | YES | YES | YES | YES | P/MQ |
| RAP1B Ras-related protein Rap-1b | Q99JI6 | NO | NO | YES | YES | YES | YES | P/MQ |
| SFXN1 Sideroflexin-1 | Q99JR1 | YES | NO | YES | YES | YES | NO | P/MQ |
| TINAL Tubulointerstitial nephritis antigen-like | Q99JR5 | NO | YES | NO | NO | NO | NO | P |
| STK26 Serine/threonine-protein kinase 26 | Q99JT2 | NO | NO | NO | NO | NO | YES | P/MQ |
| ACY1 Aminoacylase-1 | Q99JW2 | NO | YES | NO | NO | NO | NO | P |
| GORS2 Golgi reassembly-stacking protein 2 | Q99JX3 | NO | NO | NO | YES | YES | NO | P/MQ |
| ECHB Trifunctional enzyme subunit beta, mitochondrial | Q99JY0 | YES | NO | NO | YES | YES | NO | P |
| GIMA4 GTPase IMAP family member 4 | Q99JY3 | YES | NO | YES | YES | YES | YES | P |

| | | | | | | | | |
|---|---|---|---|---|---|---|---|---|
| PLPP3 Phospholipid phosphatase 3 | Q99JY8 | NO | YES | NO | NO | NO | NO | P |
| ARP3 Actin-related protein 3 | Q99JY9 | YES | NO | YES | YES | YES | YES | P/MQ |
| PDXD1 Pyridoxal-dependent decarboxylase domain-containing protein 1 | Q99K01 | NO | YES | NO | NO | NO | NO | P |
| ADP-ribosylation factor GTPase-activating protein 2 | Q99K28 | YES | YES | YES | YES | YES | YES | MQ |
| ES8L2 Epidermal growth factor receptor kinase substrate 8-like protein 2 | Q99K30 | YES | YES | NO | NO | NO | NO | P/MQ |
| EMIL1 EMILIN-1 | Q99K41 | YES | NO | YES | YES | YES | YES | P/MQ |
| 00 non-POU domain-containing octamer-binding protein | Q99K48 | NO | NO | YES | YES | YES | YES | P |
| PLST Plastin-3 | Q99K51 | YES | NO | YES | NO | YES | NO | P/MQ |
| AASS Alpha-aminoadipic semialdehyde synthase, mitochondrial | Q99K67 | YES | NO | NO | NO | YES | NO | P |
| SERC Phosphoserine aminotransferase | Q99K85 | YES | NO | NO | NO | NO | NO | P |
| GLO2 Hydroxyacylglutathione hydrolase, mitochondrial | Q99KB8 | NO | YES | NO | NO | NO | NO | P/MQ |
| VMA5A von Willebrand factor A domain-containing protein 5A | Q99KC8 | YES | NO | YES | YES | YES | YES | P/MQ |
| ME2 NAD-dependent malic enzyme, mitochondrial | Q99KE1 | NO | YES | NO | NO | NO | NO | P |
| TMED9 Transmembrane emp24 domain-containing protein 9 | Q99KF1 | NO | NO | YES | YES | YES | YES | P/MQ |
| RBM10 RNA-binding protein 10 | Q99KG3 | NO | NO | NO | YES | NO | NO | P |
| STK24 Serine/threonine-protein kinase 24 | Q99KH8 | NO | NO | YES | YES | YES | NO | P/MQ |
| DCTN2 Dynactin subunit 2 | Q99KJ8 | YES | NO | YES | YES | YES | YES | P/MQ |
| DPP3 Dipeptidyl peptidase 3 | Q99KK7 | NO | NO | NO | NO | YES | NO | P |
| EPN4 Clathrin interactor 1 | Q99KN9 | YES | NO | YES | YES | YES | YES | P/MQ |
| CRYL1 Lambda-crystallin homolog | Q99KP3 | YES | NO | NO | NO | YES | NO | P/MQ |
| PRP19 Pre-mRNA-processing factor 19 | Q99KP6 | NO | NO | YES | YES | NO | NO | P/MQ |
| NAMPT Nicotinamide phosphoribosyltransferase | Q99KQ4 | YES | NO | YES | YES | YES | YES | P/MQ |
| LACB2 Endoribonuclease LACTB2 | Q99KR3 | NO | YES | NO | NO | NO | NO | P |
| FUCO2 Plasma alpha-L-fucosidase | Q99KR8 | NO | YES | NO | NO | NO | NO | P |
| VMP1 Vacuole membrane protein 1 | Q99KU0 | NO | YES | NO | NO | NO | NO | P |
| DJB11 DnaJ homolog subfamily B member 11 | Q99KV1 | YES | NO | YES | YES | YES | YES | P/MQ |
| GAK Cyclin-G-associated kinase | Q99KY4 | NO | NO | NO | YES | NO | NO | P |
| DHRS1 Dehydrogenase/reductase SDR family member 1 | Q99L04 | NO | NO | NO | YES | YES | NO | P |
| 3HIDH 3-hydroxyisobutyrate dehydrogenase, mitochondrial | Q99L13 | YES | NO | YES | YES | YES | YES | P/MQ |
| CDS2 Phosphatidate cytidylyltransferase 2 | Q99L43 | NO | YES | NO | NO | NO | NO | P/MQ |
| IF2B Eukaryotic translation initiation factor 2 subunit 2 | Q99L45 | NO | NO | YES | NO | NO | YES | P/MQ |
| F10A1 Hsc70-interacting protein | Q99L47 | NO | NO | YES | YES | YES | YES | P/MQ |
| VATC2 V-type proton ATPase subunit C 2 | Q99L60 | NO | YES | NO | NO | NO | NO | P |
| DHRS4 Dehydrogenase/reductase SDR family member 4 | Q99LB2 | YES | YES | NO | NO | NO | NO | P |

| | | | | | | | | |
|---|---|---|---|---|---|---|---|---|
| SARDH Sarcosine dehydrogenase, mitochondrial | Q99LB7 | YES | NO | NO | NO | YES | NO | P/MQ |
| NDUAA NADH dehydrogenase [ubiquinone] 1 alpha subcomplex subunit 10, mitochondrial | Q99LC3 | YES | YES | NO | NO | NO | NO | P |
| ETFA Electron transfer flavoprotein subunit alpha, mitochondrial | Q99LC5 | YES | NO | YES | YES | YES | YES | P/MQ |
| CSN1 COP9 signalosome complex subunit 1 | Q99LD4 | NO | YES | NO | NO | NO | NO | P |
| DDAH2 N(G),N(G)-dimethylarginine dimethylaminohydrolase 2 | Q99LD8 | NO | NO | NO | NO | NO | YES | P/MQ |
| RTCB tRNA-splicing ligase RtcB homolog | Q99LF4 | NO | NO | YES | YES | YES | NO | P |
| HGS Hepatocyte growth factor-regulated tyrosine kinase substrate | Q99LI8 | NO | YES | NO | NO | NO | NO | P |
| CT2NL CTTNBP2 N-terminal-like protein | Q99LJ0 | NO | NO | NO | YES | NO | NO | P |
| DPY30 Protein dpy-30 homolog | Q99LT0 | NO | NO | NO | NO | NO | YES | P |
| PARK7 Protein/nucleic acid deglycase DJ-1 | Q99LX0 | YES | NO | YES | YES | YES | NO | P |
| PPIP2 Proline-serine-threonine phosphatase-interacting protein 2 | Q99M15 | NO | YES | NO | NO | NO | NO | P |
| RNPS1 RNA-binding protein with serine-rich domain 1 | Q99M28 | NO | YES | NO | NO | NO | NO | P/MQ |
| NCK1 Cytoplasmic protein NCK1 | Q99M51 | NO | NO | NO | YES | NO | NO | P |
| DNJA3 DnaJ homolog subfamily A member 3, mitochondrial | Q99M87 | NO | YES | NO | NO | NO | NO | P |
| ARBK1 Beta-adrenergic receptor kinase 1 | Q99MK8 | NO | NO | NO | YES | NO | NO | P |
| SYK Lysine--tRNA ligase | Q99MN1 | YES | NO | YES | YES | YES | YES | P/MQ |
| PCCB Propionyl-CoA carboxylase beta chain, mitochondrial | Q99MN9 | YES | NO | YES | NO | YES | NO | P/MQ |
| SRRT Serrate RNA effector molecule homolog | Q99MR6 | NO | NO | YES | YES | NO | NO | P/MQ |
| MCCA Methylcrotonoyl-CoA carboxylase subunit alpha, mitochondrial | Q99MR8 | YES | YES | NO | NO | NO | NO | P |
| MYO7B Unconventional myosin-VIIb | Q99MZ6 | NO | YES | NO | NO | NO | NO | P |
| CAH15 Carbonic anhydrase 15 | Q99N23 | NO | YES | NO | NO | NO | NO | P |
| RM27 39S ribosomal protein L27, mitochondrial | Q99N92 | NO | YES | NO | NO | NO | NO | P |
| RM16 39S ribosomal protein L16, mitochondrial | Q99N93 | NO | YES | NO | NO | NO | NO | P |
| ACSS1 Acetyl-coenzyme A synthetase 2-like, mitochondrial | Q99NB1 | YES | YES | NO | NO | NO | NO | P |
| SF3B1 Splicing factor 3B subunit 1 | Q99NB9 | YES | NO | YES | YES | YES | YES | P/MQ |
| RTN4 Reticulon-4 | Q99P72 | NO | NO | YES | YES | YES | YES | P/MQ |
| OGFR Opioid growth factor receptor | Q99PG2 | NO | NO | NO | YES | YES | NO | P |
| TS1R1 Taste receptor type 1 member 1 | Q99PG6 | NO | YES | NO | NO | NO | NO | P |
| GDIR1 Rho GDP-dissociation inhibitor 1 | Q99PT1 | YES | NO | YES | YES | YES | YES | P/MQ |
| PRP8 Pre-mRNA-processing-splicing factor 8 | Q99PV0 | NO | NO | YES | YES | YES | YES | P/MQ |
| COX6C Cytochrome c oxidase subunit 6C | Q9CPQ1 | NO | NO | YES | NO | NO | NO | P/MQ |
| ATP5L ATP synthase subunit g, mitochondrial | Q9CPQ8 | NO | NO | NO | NO | YES | NO | P/MQ |
| RL17 60S ribosomal protein L17 | Q9CPR4 | YES | NO | YES | YES | YES | YES | P/MQ |
| MYDGF Myeloid-derived growth factor | Q9CPT4 | NO | NO | NO | NO | YES | NO | P |

| | | | | | | | | |
|---|---|---|---|---|---|---|---|---|
| GLOD4 Glyoxalase domain-containing protein 4 | Q9CPV4 | YES | NO | YES | YES | YES | YES | P/MQ |
| RM54 39S ribosomal protein L54, mitochondrial | Q9CPW3 | NO | YES | NO | NO | NO | NO | P |
| ARPC5 Actin-related protein 2/3 complex subunit 5 | Q9CPW4 | NO | NO | YES | YES | YES | YES | P/MQ |
| ATG3 Ubiquitin-like-conjugating enzyme ATG3 | Q9CPX6 | NO | NO | NO | YES | YES | NO | P |
| AMPL Cytosol aminopeptidase | Q9CPY7 | YES | NO | YES | YES | YES | YES | P/MQ |
| MYL9 Myosin regulatory light polypeptide 9 | Q9CQ19 | NO | YES | NO | NO | NO | NO | P |
| LTOR1 Ragulator complex protein LAMTOR1 | Q9CQ22 | NO | YES | NO | NO | NO | NO | P |
| RM49 39S ribosomal protein L49, mitochondrial | Q9CQ40 | NO | YES | NO | NO | NO | NO | P |
| NDUC2 NADH dehydrogenase [ubiquinone] 1 subunit C2 | Q9CQ54 | NO | YES | NO | NO | NO | NO | P |
| MTAP S-methyl-5'-thioadenosine phosphorylase | Q9CQ65 | NO | NO | NO | YES | YES | NO | P |
| QCR8 Cytochrome b-c1 complex subunit 8 | Q9CQ69 | YES | NO | YES | YES | NO | NO | P |
| NDUA2 NADH dehydrogenase [ubiquinone] 1 alpha subcomplex subunit 2 | Q9CQ75 | YES | NO | YES | NO | YES | NO | P/MQ |
| FIS1 Mitochondrial fission 1 protein | Q9CQ92 | NO | NO | NO | YES | NO | NO | P/MQ |
| SDHB Succinate dehydrogenase [ubiquinone] iron-sulfur subunit, mitochondrial | Q9CQA3 | YES | NO | YES | NO | YES | NO | P |
| BZW1 Basic leucine zipper and W2 domain-containing protein 1 | Q9CQC6 | NO | NO | NO | YES | NO | NO | P |
| NDUB4 NADH dehydrogenase [ubiquinone] 1 beta subcomplex subunit 4 | Q9CQC7 | YES | NO | YES | NO | YES | NO | P/MQ |
| RAB5A Ras-related protein Rab-5A | Q9CQD1 | NO | YES | NO | NO | NO | NO | P |
| RGS10 Regulator of G-protein signaling 10 | Q9CQE5 | NO | NO | NO | YES | NO | NO | P |
| RTRAF RNA transcription, translation and transport factor protein | Q9CQE8 | YES | NO | NO | YES | YES | YES | P |
| RM11 39S ribosomal protein L11, mitochondrial | Q9CQF0 | NO | YES | NO | NO | NO | NO | P |
| CPSF5 Cleavage and polyadenylation specificity factor subunit 5 | Q9CQF3 | NO | NO | YES | YES | YES | YES | P/MQ |
| PDZ1I PDZK1-interacting protein 1 | Q9CQH0 | NO | YES | NO | NO | NO | NO | P/MQ |
| NDUB5 NADH dehydrogenase [ubiquinone] 1 beta subcomplex subunit 5, mitochondrial | Q9CQH3 | YES | YES | NO | NO | NO | NO | P |
| BT3L4 Transcription factor BTF3 homolog 4 | Q9CQH7 | NO | YES | NO | NO | NO | NO | P |
| GMFB Glia maturation factor beta | Q9CQI3 | NO | NO | NO | YES | NO | NO | P |
| COTL1 Coactosin-like protein | Q9CQI6 | YES | NO | YES | YES | YES | YES | P/MQ |
| DENR Density-regulated protein | Q9CQJ6 | NO | NO | NO | NO | NO | YES | P |
| RM18 39S ribosomal protein L18, mitochondrial | Q9CQL5 | NO | YES | NO | NO | NO | NO | P |
| MOFA1 MORF4 family-associated protein 1 | Q9CQL7 | NO | YES | NO | NO | NO | NO | P |
| TXD17 Thioredoxin domain-containing protein 17 | Q9CQM5 | NO | YES | NO | NO | NO | NO | P |
| GLRX3 Glutaredoxin-3 | Q9CQM9 | NO | NO | YES | YES | YES | NO | P |
| TRAP1 Heat shock protein 75 kDa, mitochondrial | Q9CQN1 | YES | NO | YES | YES | YES | NO | P/MQ |
| AT5F1 ATP synthase F(0) complex subunit B1, mitochondrial | Q9CQQ7 | YES | NO | NO | YES | YES | YES | P |
| RS21 40S ribosomal protein S21 | Q9CQR2 | YES | NO | YES | YES | YES | YES | P/MQ |

| | | | | | | | | |
|---|---|---|---|---|---|---|---|---|
| ACOT13 Acyl-coenzyme A thioesterase 13 | Q9CQR4 | NO | YES | NO | NO | NO | NO | P |
| SC61B Protein transport protein Sec61 subunit beta | Q9CQS8 | NO | NO | NO | YES | YES | YES | P/MQ |
| MTNA Methylthioribose-1-phosphate isomerase | Q9CQT1 | NO | YES | NO | NO | NO | NO | P |
| TXD12 Thioredoxin domain-containing protein 12 | Q9CQU0 | NO | YES | NO | NO | NO | NO | P |
| RER1 Protein RER1 | Q9CQU3 | NO | YES | NO | NO | NO | NO | P |
| TIM14 Mitochondrial import inner membrane translocase subunit TIM14 | Q9CQV7 | NO | YES | NO | NO | NO | NO | P |
| 1433B 14-3-3 protein beta/alpha | Q9CQV8 | YES | NO | YES | YES | YES | YES | P/MQ |
| YKT6 Synaptobrevin homolog YKT6 | Q9CQW1 | NO | NO | NO | NO | YES | NO | P |
| ARL8B ADP-ribosylation factor-like protein 8B | Q9CQW2 | NO | YES | NO | NO | NO | NO | P |
| RT36 28S ribosomal protein S36, mitochondrial | Q9CQX8 | YES | NO | NO | NO | NO | NO | P |
| NDUA6 NADH dehydrogenase [ubiquinone] 1 alpha subcomplex subunit 6 | Q9CQZ5 | NO | YES | NO | NO | NO | NO | P |
| NDUB3 NADH dehydrogenase [ubiquinone] 1 beta subcomplex subunit 3 | Q9CQZ6 | NO | NO | NO | YES | YES | YES | P/MQ |
| PSMD9 26S proteasome non-ATPase regulatory subunit 9 | Q9CR00 | NO | NO | NO | NO | YES | NO | P |
| UFC1 Ubiquitin-fold modifier-conjugating enzyme 1 | Q9CR09 | NO | YES | NO | NO | NO | NO | P |
| PPID Peptidyl-prolyl cis-trans isomerase D | Q9CR16 | NO | NO | NO | YES | YES | YES | P/MQ |
| VTA1 Vacuolar protein sorting-associated protein VTA1 homolog | Q9CR26 | NO | NO | NO | NO | YES | NO | P |
| VATG1 V-type proton ATPase subunit G 1 | Q9CR51 | YES | NO | YES | YES | YES | YES | P/MQ |
| RL14 60S ribosomal protein L14 | Q9CR57 | YES | NO | YES | YES | YES | YES | P |
| NADH dehydrogenase [ubiquinone] 1 beta subcomplex subunit 7 | Q9CR61 | YES | YES | NO | YES | YES | YES | MQ |
| M2OM Mitochondrial 2-oxoglutarate/malate carrier protein | Q9CR62 | NO | YES | NO | NO | NO | NO | P |
| UCRI Cytochrome b-c1 complex subunit Rieske, mitochondrial | Q9CR68 | YES | NO | YES | YES | YES | YES | P |
| CHSP1 Calcium-regulated heat stable protein 1 | Q9CR86 | NO | NO | NO | NO | YES | NO | P |
| NHP2 H/ACA ribonucleoprotein complex subunit 2 | Q9CRB2 | NO | YES | NO | NO | NO | NO | P |
| Tubulin polymerization-promoting protein family member 3 | Q9CRB6 | YES | YES | YES | YES | YES | YES | MQ |
| MIC19 MICOS complex subunit Mic19 | Q9CRB9 | YES | NO | YES | YES | YES | YES | P/MQ |
| OCAD1 OCIA domain-containing protein 1 | Q9CRD0 | YES | NO | NO | YES | YES | NO | P/MQ |
| EMC2 ER membrane protein complex subunit 2 | Q9CRD2 | NO | YES | NO | NO | NO | NO | P |
| PRPS2 Ribose-phosphate pyrophosphokinase 2 | Q9CS42 | NO | YES | NO | NO | NO | NO | P |
| SNW1 SNW domain-containing protein 1 | Q9CSN1 | NO | NO | YES | YES | YES | YES | P |
| RPR1B Regulation of nuclear pre-mRNA domain-containing protein 1B | Q9CSU0 | NO | NO | NO | YES | YES | NO | P/MQ |
| RANB3 Ran-binding protein 3 | Q9CT10 | NO | NO | NO | YES | YES | YES | P/MQ |
| SMC1A Structural maintenance of chromosomes protein 1A | Q9CU62 | NO | NO | NO | YES | YES | YES | P/MQ |
| ARPC2 Actin-related protein 2/3 complex subunit 2 | Q9CVB6 | NO | NO | YES | YES | YES | YES | P/MQ |

| | | | | | | | | |
|---|---|---|---|---|---|---|---|---|
| SMC3 Structural maintenance of chromosomes protein 3 | Q9CW03 | NO | NO | YES | YES | YES | YES | P/MQ |
| RAVR1 Ribonucleoprotein PTB-binding 1 | Q9CW46 | NO | NO | NO | YES | YES | NO | P/MQ |
| TBB2B Tubulin beta-2B chain | Q9CWF2 | NO | YES | NO | NO | NO | NO | P/MQ |
| PUR9 Bifunctional purine biosynthesis protein PURH | Q9CWJ9 | YES | NO | YES | YES | YES | YES | P/MQ |
| SNX2 Sorting nexin-2 | Q9CWK8 | YES | NO | YES | YES | YES | YES | P/MQ |
| PFD1 Prefoldin subunit 1 | Q9CWM4 | NO | NO | NO | YES | NO | NO | P |
| DDAH1 N(G),N(G)-dimethylarginine dimethylaminohydrolase 1 | Q9CWS0 | YES | NO | YES | YES | YES | NO | P/MQ |
| RBM8A RNA-binding protein 8A | Q9CWZ3 | NO | YES | NO | NO | NO | NO | P |
| IST1 IST1 homolog | Q9CX00 | NO | YES | NO | NO | NO | NO | P |
| SGT1 Protein SGT1 homolog | Q9CX34 | NO | NO | NO | NO | NO | YES | P |
| CYGB Cytoglobin | Q9CX80 | NO | YES | NO | NO | NO | NO | P |
| ROA0 Heterogeneous nuclear ribonucleoprotein A0 | Q9CX86 | NO | NO | YES | YES | YES | YES | P/MQ |
| T3HPD Trans-L-3-hydroxyproline dehydratase | Q9CXA2 | NO | YES | NO | NO | NO | NO | P |
| TBC15 TBC1 domain family member 15 | Q9CXF4 | NO | NO | NO | NO | NO | YES | P |
| ABCB8 ATP-binding cassette sub-family B member 8, mitochondrial | Q9CXJ4 | NO | YES | NO | NO | NO | NO | P |
| CG050 Uncharacterized protein C7orf50 homolog | Q9CXL3 | NO | NO | NO | YES | NO | YES | P |
| DHRS7 Dehydrogenase/reductase SDR family member 7 | Q9CXR1 | NO | NO | YES | YES | YES | YES | P |
| CENPV Centromere protein V | Q9CXS4 | YES | NO | YES | YES | NO | NO | P/MQ |
| MPPB Mitochondrial-processing peptidase subunit beta | Q9CXT8 | NO | YES | NO | NO | NO | NO | P |
| CYBP Calcyclin-binding protein | Q9CXW3 | NO | NO | NO | YES | NO | NO | P |
| RL11 60S ribosomal protein L11 | Q9CXW4 | YES | NO | YES | YES | YES | YES | P/MQ |
| ILF2 Interleukin enhancer-binding factor 2 | Q9CXY6 | NO | YES | NO | NO | NO | NO | P/MQ |
| NDUS4 NADH dehydrogenase [ubiquinone] iron-sulfur protein 4, mitochondrial | Q9CXZ1 | YES | NO | YES | NO | YES | NO | P/MQ |
| RT28 28S ribosomal protein S28, mitochondrial | Q9CY16 | NO | YES | NO | NO | NO | NO | P |
| SNX7 Sorting nexin-7 | Q9CY18 | NO | YES | NO | NO | NO | NO | P |
| SSRA Translocon-associated protein subunit alpha | Q9CY50 | NO | NO | NO | NO | YES | NO | P |
| PAIRB Plasminogen activator inhibitor 1 RNA-binding protein | Q9CY58 | YES | NO | YES | YES | YES | YES | P/MQ |
| BIEA Biliverdin reductase A | Q9CY64 | YES | NO | YES | YES | YES | YES | P |
| TOM34 Mitochondrial import receptor subunit TOM34 | Q9CYG7 | YES | NO | YES | YES | YES | YES | P/MQ |
| PXL2A Peroxiredoxin-like 2A | Q9CYH2 | NO | YES | NO | NO | NO | NO | P |
| LUC7L Putative RNA-binding protein Luc7-like 1 | Q9CYI4 | NO | YES | NO | NO | NO | NO | P/MQ |
| GAPR1 Golgi-associated plant pathogenesis-related protein 1 | Q9CYL5 | NO | NO | NO | YES | NO | NO | P |
| SPCS2 Signal peptidase complex subunit 2 | Q9CYN2 | NO | YES | NO | NO | NO | NO | P |
| TPD54 Tumor protein D54 | Q9CYZ2 | YES | NO | YES | YES | YES | YES | P/MQ |
| CSN7A COP9 signalosome complex subunit 7a | Q9CZ04 | NO | NO | NO | YES | NO | NO | P |

| | | | | | | | | |
|---|---|---|---|---|---|---|---|---|
| QCR1 Cytochrome b-c1 complex subunit 1, mitochondrial | Q9CZ13 | YES | NO | YES | YES | YES | YES | P/MQ |
| SNF8 Vacuolar-sorting protein SNF8 | Q9CZ28 | NO | YES | NO | NO | NO | NO | P |
| OLA1 Obg-like ATPase 1 | Q9CZ30 | NO | NO | NO | YES | NO | YES | P |
| NNRD ATP-dependent (S)-NAD(P)H-hydrate dehydratase | Q9CZ42 | NO | YES | NO | NO | NO | NO | P |
| NSF1C NSFL1 cofactor p47 | Q9CZ44 | YES | NO | YES | YES | YES | YES | P/MQ |
| MMAC Methylmalonic aciduria and homocystinuria type C protein homolog | Q9CZD0 | NO | YES | NO | NO | NO | NO | P |
| GARS Glycine--tRNA ligase | Q9CZD3 | NO | NO | YES | YES | YES | YES | P |
| RAB32 Ras-related protein Rab-32 | Q9CZE3 | NO | NO | YES | YES | YES | YES | P |
| MXRA7 Matrix-remodeling-associated protein 7 | Q9CZH7 | NO | NO | NO | NO | NO | YES | P |
| RL15 60S ribosomal protein L15 | Q9CZM2 | YES | NO | YES | YES | YES | YES | P/MQ |
| GLYM Serine hydroxymethyltransferase, mitochondrial | Q9CZN7 | YES | NO | YES | YES | YES | NO | P/MQ |
| MTREX Exosome RNA helicase MTR4 | Q9CZU3 | NO | NO | NO | NO | YES | NO | P |
| TOM70 Mitochondrial import receptor subunit TOM70 | Q9CZW5 | NO | NO | NO | YES | NO | NO | P |
| RS19 40S ribosomal protein S19 | Q9CZX8 | YES | NO | YES | YES | YES | YES | P/MQ |
| UB2V1 Ubiquitin-conjugating enzyme E2 variant 1 | Q9CZY3 | NO | NO | YES | YES | YES | NO | P/MQ |
| 5NT3A Cytosolic 5'-nucleotidase 3A | Q9D020 | NO | NO | NO | NO | YES | NO | P/MQ |
| PDHB Pyruvate dehydrogenase E1 component subunit beta, mitochondrial | Q9D051 | YES | NO | YES | YES | YES | YES | P/MQ |
| SRSF9 Serine/arginine-rich splicing factor 9 | Q9D0B0 | NO | NO | NO | YES | NO | NO | P |
| HNRPM Heterogeneous nuclear ribonucleoprotein M | Q9D0E1 | YES | NO | YES | YES | YES | YES | P/MQ |
| LMAN1 Protein ERGIC-53 | Q9D0F3 | NO | NO | NO | YES | YES | YES | P/MQ |
| PGM1 Phosphoglucomutase-1 | Q9D0F9 | YES | NO | YES | YES | YES | YES | P/MQ |
| SYRC Arginine--tRNA ligase, cytoplasmic | Q9D0I9 | NO | NO | YES | YES | YES | YES | P/MQ |
| PTMS Parathymosin | Q9D0J8 | YES | NO | YES | YES | YES | YES | P/MQ |
| MCES mRNA cap guanine-N7 methyltransferase | Q9D0L8 | NO | NO | NO | YES | NO | NO | P |
| CY1 Cytochrome c1, heme protein, mitochondrial | Q9D0M3 | YES | NO | YES | YES | NO | NO | P |
| SYTC Threonine--tRNA ligase, cytoplasmic | Q9D0R2 | NO | NO | NO | YES | NO | YES | P |
| HINT2 Histidine triad nucleotide-binding protein 2, mitochondrial | Q9D0S9 | YES | NO | YES | NO | YES | NO | P |
| NH2L1 NHP2-like protein 1 | Q9D0T1 | NO | NO | YES | YES | YES | YES | P |
| ILEUA Leukocyte elastase inhibitor A | Q9D154 | NO | NO | YES | YES | YES | YES | P/MQ |
| GAL3A Glutamine amidotransferase-like class 1 domain-containing protein 3A, mitochondrial | Q9D172 | YES | NO | YES | YES | NO | NO | P/MQ |
| CNDP2 Cytosolic non-specific dipeptidase | Q9D1A2 | YES | NO | YES | YES | YES | YES | P/MQ |
| RM28 39S ribosomal protein L28, mitochondrial | Q9D1B9 | NO | YES | NO | NO | NO | NO | P |
| TMEDA Transmembrane emp24 domain-containing protein 10 | Q9D1D4 | NO | NO | YES | YES | NO | YES | P/MQ |
| TBCB Tubulin-folding cofactor B | Q9D1E6 | NO | NO | NO | YES | YES | NO | P |
| RAB1B Ras-related protein Rab-1B | Q9D1G1 | NO | NO | NO | YES | YES | NO | P |
| RM14 39S ribosomal protein L14, mitochondrial | Q9D1I6 | NO | YES | NO | NO | NO | NO | P |

| | | | | | | | | |
|---|---|---|---|---|---|---|---|---|
| NECP2 Adaptin ear-binding coat-associated protein 2 | Q9D1J1 | NO | NO | NO | YES | NO | NO | P/MQ |
| SARNP SAP domain-containing ribonucleoprotein | Q9D1J3 | YES | NO | NO | YES | YES | YES | P/MQ |
| VATF V-type proton ATPase subunit F | Q9D1K2 | NO | NO | NO | YES | YES | YES | P |
| FKB11 Peptidyl-prolyl cis-trans isomerase FKBP11 | Q9D1M7 | NO | NO | NO | NO | YES | NO | P |
| RM13 39S ribosomal protein L13, mitochondrial | Q9D1P0 | NO | YES | NO | NO | NO | NO | P |
| CHRD1 Cysteine and histidine-rich domain-containing protein 1 | Q9D1P4 | NO | NO | NO | YES | NO | NO | P |
| ERP44 Endoplasmic reticulum resident protein 44 | Q9D1Q6 | NO | NO | NO | YES | NO | YES | P/MQ |
| RL34 60S ribosomal protein L34 | Q9D1R9 | NO | NO | NO | NO | YES | NO | P |
| noL3 Nucleolar protein 3 | Q9D1X0 | NO | YES | NO | NO | NO | NO | P |
| SPF27 Pre-mRNA-splicing factor SPF27 | Q9D287 | NO | NO | NO | YES | NO | NO | P |
| DLST Dihydrolipoyllysine-residue succinyltransferase component of 2-oxoglutarate dehydrogenase complex, mitochondrial | Q9D2G2 | YES | NO | YES | YES | YES | NO | P/MQ |
| UB2V2 Ubiquitin-conjugating enzyme E2 variant 2 | Q9D2M8 | NO | YES | NO | NO | NO | NO | P |
| AACS Acetoacetyl-CoA synthetase | Q9D2R0 | NO | NO | NO | YES | NO | NO | P |
| COA3 Cytochrome c oxidase assembly factor 3 homolog, mitochondrial | Q9D2R6 | NO | YES | NO | NO | NO | NO | P |
| CORO7 Coronin-7 | Q9D2V7 | NO | NO | YES | YES | YES | YES | P/MQ |
| K1C20 Keratin, type I cytoskeletal 20 | Q9D312 | NO | YES | NO | NO | NO | NO | P |
| RM19 39S ribosomal protein L19, mitochondrial | Q9D338 | NO | YES | NO | NO | NO | NO | P |
| PPAC Low molecular weight phosphotyrosine protein phosphatase | Q9D358 | NO | NO | NO | NO | YES | NO | P |
| HYEP Epoxide hydrolase 1 | Q9D379 | NO | YES | NO | NO | NO | NO | P |
| ATPD ATP synthase subunit delta, mitochondrial | Q9D3D9 | YES | NO | YES | YES | YES | YES | P/MQ |
| SF3A3 Splicing factor 3A subunit 3 | Q9D554 | NO | NO | YES | YES | YES | YES | P/MQ |
| SYAP1 Synapse-associated protein 1 | Q9D5V6 | NO | NO | NO | YES | NO | NO | P/MQ |
| RFIP1 Rab11 family-interacting protein 1 | Q9D620 | NO | NO | NO | YES | NO | NO | P |
| SC23B Protein transport protein Sec23B | Q9D662 | NO | NO | NO | YES | NO | NO | P |
| S6A19 Sodium-dependent neutral amino acid transporter B(0)AT1 | Q9D687 | NO | YES | NO | NO | NO | NO | P |
| NDUB8 NADH dehydrogenase [ubiquinone] 1 beta subcomplex subunit 8, mitochondrial | Q9D6J5 | NO | NO | NO | NO | YES | NO | P |
| NDUV2 NADH dehydrogenase [ubiquinone] flavoprotein 2, mitochondrial | Q9D6J6 | YES | NO | YES | YES | YES | NO | P/MQ |
| IDH3A Isocitrate dehydrogenase [NAD] subunit alpha, mitochondrial | Q9D6R2 | YES | NO | YES | YES | YES | YES | P/MQ |
| RRFM Ribosome-recycling factor, mitochondrial | Q9D6S7 | NO | YES | NO | NO | NO | NO | P |
| MSRA Mitochondrial peptide methionine sulfoxide reductase | Q9D6Y7 | YES | NO | YES | YES | YES | YES | P/MQ |
| noP56 Nucleolar protein 56 | Q9D6Z1 | YES | NO | YES | YES | YES | YES | P |
| TMX2 Thioredoxin-related transmembrane protein 2 | Q9D710 | NO | YES | NO | NO | NO | NO | P |
| TRIR Telomerase RNA component interacting RNase | Q9D735 | NO | NO | NO | YES | NO | NO | P |
| PPIL2 RING-type E3 ubiquitin-protein ligase PPIL2 | Q9D787 | NO | YES | NO | NO | NO | NO | P |

| | | | | | | | | |
|---|---|---|---|---|---|---|---|---|
| ACAD8 Isobutyryl-CoA dehydrogenase, mitochondrial | Q9D7B6 | NO | YES | NO | NO | NO | NO | P |
| RTCA RNA 3'-terminal phosphate cyclase | Q9D7H3 | NO | YES | NO | NO | NO | NO | P |
| APMAP Adipocyte plasma membrane-associated protein | Q9D7N9 | NO | YES | NO | NO | NO | NO | P |
| CHMP5 Charged multivesicular body protein 5 | Q9D7S9 | NO | NO | NO | YES | NO | NO | P |
| DUS3 Dual specificity protein phosphatase 3 | Q9D7X3 | NO | NO | NO | YES | NO | NO | P |
| IPYR Inorganic pyrophosphatase | Q9D819 | NO | NO | YES | NO | YES | YES | P/MQ |
| PRXD1 Prolyl-tRNA synthetase associated domain-containing protein 1 | Q9D820 | NO | YES | NO | NO | NO | NO | P |
| FIP1 Pre-mRNA 3'-end-processing factor FIP1 | Q9D824 | NO | NO | NO | YES | NO | NO | P/MQ |
| SOX Peroxisomal sarcosine oxidase | Q9D826 | YES | NO | NO | NO | YES | NO | P |
| QCR7 Cytochrome b-c1 complex subunit 7 | Q9D855 | YES | NO | YES | YES | YES | YES | P/MQ |
| TIM50 Mitochondrial import inner membrane translocase subunit TIM50 | Q9D880 | NO | NO | NO | NO | YES | NO | P |
| U2AF1 Splicing factor U2AF 35 kDa subunit | Q9D883 | NO | NO | NO | NO | NO | YES | P |
| ITPA Inosine triphosphate pyrophosphatase | Q9D892 | NO | YES | NO | NO | NO | NO | P/MQ |
| ARP5L Actin-related protein 2/3 complex subunit 5-like protein | Q9D898 | NO | YES | NO | NO | NO | NO | P |
| CHM4B Charged multivesicular body protein 4b | Q9D8B3 | YES | NO | YES | YES | YES | YES | P/MQ |
| RL4 60S ribosomal protein L4 | Q9D8E6 | YES | NO | YES | YES | YES | YES | P/MQ |
| GLOD5 Glyoxalase domain-containing protein 5 | Q9D8I3 | NO | YES | NO | NO | NO | NO | P |
| EF1G Elongation factor 1-gamma | Q9D8N0 | YES | NO | YES | YES | YES | YES | P/MQ |
| ORN Oligoribonuclease, mitochondrial | Q9D8S4 | NO | YES | NO | NO | NO | NO | P |
| GSDMD Gasdermin-D | Q9D8T2 | NO | NO | NO | YES | NO | NO | P |
| SNX5 Sorting nexin-5 | Q9D8U8 | YES | NO | YES | YES | YES | YES | P/MQ |
| PSD12 26S proteasome non-ATPase regulatory subunit 12 | Q9D8W5 | NO | NO | NO | YES | YES | YES | P |
| CC124 Coiled-coil domain-containing protein 124 | Q9D8X2 | NO | NO | YES | YES | YES | NO | P |
| EFHD2 EF-hand domain-containing protein D2 | Q9D8Y0 | YES | NO | YES | YES | YES | YES | P/MQ |
| T126A Transmembrane protein 126A | Q9D8Y1 | NO | YES | NO | NO | NO | NO | P |
| ST1C2 Sulfotransferase 1C2 | Q9D939 | NO | YES | NO | NO | NO | NO | P |
| DCPS m7GpppX diphosphatase | Q9DAR7 | NO | NO | YES | YES | NO | YES | P/MQ |
| MTU1 Mitochondrial tRNA-specific 2-thiouridylase 1 | Q9DAT5 | NO | YES | NO | NO | NO | NO | P |
| CNPY3 Protein canopy homolog 3 | Q9DAU1 | NO | NO | NO | YES | YES | YES | P/MQ |
| CNN3 Calponin-3 | Q9DAW9 | NO | NO | NO | NO | YES | NO | P/MQ |
| SNAA Alpha-soluble NSF attachment protein | Q9DB05 | NO | NO | NO | YES | YES | YES | P/MQ |
| RM12 39S ribosomal protein L12, mitochondrial | Q9DB15 | YES | NO | NO | YES | NO | YES | P/MQ |
| ATPO ATP synthase subunit O, mitochondrial | Q9DB20 | YES | NO | YES | YES | YES | YES | P/MQ |
| IAH1 Isoamyl acetate-hydrolyzing esterase 1 homolog | Q9DB29 | YES | YES | NO | NO | NO | NO | P |
| CHM2A Charged multivesicular body protein 2a | Q9DB34 | NO | NO | NO | NO | NO | YES | P |
| QCR2 Cytochrome b-c1 complex subunit 2, mitochondrial | Q9DB77 | YES | NO | YES | YES | YES | YES | P/MQ |

| | | | | | | | | |
|---|---|---|---|---|---|---|---|---|
| KAP0 cAMP-dependent protein kinase type I-alpha regulatory subunit | Q9DBC7 | NO | NO | NO | YES | NO | NO | P |
| CSAD Cysteine sulfinic acid decarboxylase | Q9DBE0 | NO | YES | NO | NO | NO | NO | P |
| AL7A1 Alpha-aminoadipic semialdehyde dehydrogenase | Q9DBF1 | YES | NO | YES | NO | YES | NO | P |
| PLIN3 Perilipin-3 | Q9DBG5 | NO | NO | YES | YES | YES | YES | P |
| RPN2 Dolichyl-diphosphooligosaccharide--protein glycosyltransferase subunit 2 | Q9DBG6 | NO | NO | YES | YES | YES | YES | P/MQ |
| TX1B3 Tax1-binding protein 3 | Q9DBG9 | YES | NO | NO | YES | NO | YES | P/MQ |
| LMAN2 Vesicular integral-membrane protein VIP36 | Q9DBH5 | NO | NO | YES | YES | YES | YES | P |
| GDAP2 Ganglioside-induced differentiation-associated protein 2 | Q9DBL2 | NO | YES | NO | NO | NO | NO | P |
| COASY Bifunctional coenzyme A synthase | Q9DBL7 | NO | YES | NO | NO | NO | NO | P |
| ECHP Peroxisomal bifunctional enzyme | Q9DBM2 | YES | NO | YES | NO | YES | NO | P/MQ |
| KCY UMP-CMP kinase | Q9DBP5 | YES | NO | YES | YES | YES | YES | P/MQ |
| XRN2 5'-3' exoribonuclease 2 | Q9DBR1 | NO | NO | NO | YES | NO | YES | P/MQ |
| MYPT1 Protein phosphatase 1 regulatory subunit 12A | Q9DBR7 | YES | NO | YES | YES | YES | YES | P/MQ |
| TMM43 Transmembrane protein 43 | Q9DBS1 | NO | NO | NO | NO | YES | NO | P |
| KLC4 Kinesin light chain 4 | Q9DBS5 | NO | YES | NO | NO | NO | NO | P |
| M2GD Dimethylglycine dehydrogenase, mitochondrial | Q9DBT9 | YES | NO | YES | NO | YES | YES | P |
| SUSD2 Sushi domain-containing protein 2 | Q9DBX3 | NO | YES | NO | NO | NO | NO | P |
| KC1D Casein kinase I isoform delta | Q9DC28 | NO | YES | NO | NO | NO | NO | P |
| GNAI3 Guanine nucleotide-binding protein G(k) subunit alpha | Q9DC51 | NO | YES | NO | NO | NO | NO | P |
| MPPA Mitochondrial-processing peptidase subunit alpha | Q9DC61 | NO | YES | NO | NO | NO | NO | P |
| NDUA9 NADH dehydrogenase [ubiquinone] 1 alpha subcomplex subunit 9, mitochondrial | Q9DC69 | NO | YES | NO | NO | NO | NO | P/MQ |
| NDUS7 NADH dehydrogenase [ubiquinone] iron-sulfur protein 7, mitochondrial | Q9DC70 | NO | YES | NO | NO | NO | NO | P |
| Pyrroline-5-carboxylate reductase 3 | Q9DCC4 | YES | YES | YES | YES | YES | YES | MQ |
| Interferon gamma-induced GTPase | Q9DCE9 | YES | YES | YES | YES | YES | YES | MQ |
| SSRG Translocon-associated protein subunit gamma | Q9DCF9 | NO | NO | NO | YES | YES | YES | P |
| PBLD1 Phenazine biosynthesis-like domain-containing protein 1 | Q9DCG6 | YES | NO | NO | NO | YES | NO | P |
| ZFAN6 AN1-type zinc finger protein 6 | Q9DCH6 | NO | YES | NO | NO | NO | NO | P |
| NDUA8 NADH dehydrogenase [ubiquinone] 1 alpha subcomplex subunit 8 | Q9DCJ5 | YES | NO | YES | YES | YES | NO | P/MQ |
| NPL N-acetylneuraminate lyase | Q9DCJ9 | YES | YES | NO | NO | NO | NO | P |
| IPP2 Protein phosphatase inhibitor 2 | Q9DCL8 | NO | NO | NO | YES | NO | NO | P/MQ |
| PUR6 Multifunctional protein ADE2 | Q9DCL9 | NO | NO | NO | NO | NO | YES | P/MQ |
| ETHE1 Persulfide dioxygenase ETHE1, mitochondrial | Q9DCM0 | YES | NO | NO | YES | NO | NO | P |
| AP3S1 AP-3 complex subunit sigma-1 | Q9DCR2 | NO | NO | NO | YES | YES | YES | P |
| MTL26 Methyltransferase-like 26 | Q9DCS2 | YES | YES | NO | NO | NO | NO | P |

| | | | | | | | | |
|---|---|---|---|---|---|---|---|---|
| NDUBA NADH dehydrogenase [ubiquinone] 1 beta subcomplex subunit 10 | Q9DCS9 | YES | NO | YES | NO | YES | NO | P/MQ |
| AKCL2 1,5-anhydro-D-fructose reductase | Q9DCT1 | NO | NO | NO | YES | NO | NO | P |
| NDUS3 NADH dehydrogenase [ubiquinone] iron-sulfur protein 3, mitochondrial | Q9DCT2 | YES | NO | YES | NO | YES | NO | P |
| Cysteine-rich protein 2 | Q9DCT8 | YES | YES | NO | YES | YES | YES | MQ |
| K2C7 Keratin, type II cytoskeletal 7 | Q9DCV7 | YES | NO | YES | YES | YES | YES | P |
| ETFB Electron transfer flavoprotein subunit beta | Q9DCW4 | YES | NO | YES | YES | YES | YES | P/MQ |
| ATP5H ATP synthase subunit d, mitochondrial | Q9DCX2 | YES | NO | YES | YES | YES | YES | P/MQ |
| KEG1 Glycine N-acyltransferase-like protein Keg1 | Q9DCY0 | YES | NO | YES | NO | YES | NO | P |
| MIC26 MICOS complex subunit Mic26 | Q9DCZ4 | YES | YES | NO | NO | NO | NO | P/MQ |
| DTD1 D-aminoacyl-tRNA deacylase 1 | Q9DD18 | NO | YES | NO | NO | NO | NO | P/MQ |
| SAC1 Phosphatidylinositide phosphatase SAC1 | Q9EP69 | NO | YES | NO | NO | NO | NO | P |
| RAI14 Ankycorbin | Q9EP71 | NO | NO | NO | YES | YES | YES | P |
| ASC Apoptosis-associated speck-like protein containing a CARD | Q9EPB4 | NO | NO | NO | YES | NO | NO | P |
| PARVA Alpha-parvin | Q9EPC1 | YES | NO | YES | YES | YES | YES | P/MQ |
| ARFG1 ADP-ribosylation factor GTPase-activating protein 1 | Q9EPJ9 | NO | NO | NO | NO | YES | NO | P |
| IPO7 Importin-7 | Q9EPL8 | NO | YES | NO | NO | NO | NO | P |
| RENT1 Regulator of nonsense transcripts 1 | Q9EPU0 | NO | NO | NO | YES | YES | YES | P/MQ |
| DHB11 Estradiol 17-beta-dehydrogenase 11 | Q9EQ06 | NO | YES | NO | NO | NO | NO | P/MQ |
| MMSA Methylmalonate-semialdehyde dehydrogenase [acylating], mitochondrial | Q9EQ20 | YES | NO | YES | YES | YES | NO | P/MQ |
| VPS35 Vacuolar protein sorting-associated protein 35 | Q9EQH3 | YES | NO | YES | YES | YES | YES | P/MQ |
| MVP Major vault protein | Q9EQK5 | YES | NO | YES | YES | YES | YES | P/MQ |
| T22D4 TSC22 domain family protein 4 | Q9EQN3 | NO | NO | NO | YES | NO | NO | P |
| EHD4 EH domain-containing protein 4 | Q9EQP2 | YES | NO | YES | YES | YES | YES | P/MQ |
| SET Protein SET | Q9EQU5 | NO | NO | YES | YES | YES | YES | P/MQ |
| AIF1L Allograft inflammatory factor 1-like | Q9EQX4 | YES | YES | NO | NO | NO | NO | P |
| STX12 Syntaxin-12 | Q9ER00 | NO | NO | NO | YES | NO | YES | P |
| SYCC Cysteine--tRNA ligase, cytoplasmic | Q9ER72 | NO | YES | NO | NO | NO | NO | P |
| SNP29 Synaptosomal-associated protein 29 | Q9ERB0 | NO | NO | NO | YES | YES | YES | P |
| PARVG Gamma-parvin | Q9ERD8 | NO | NO | NO | NO | YES | NO | P |
| MESD LRP chaperone MESD | Q9ERE7 | NO | NO | NO | YES | YES | NO | P |
| LIMA1 LIM domain and actin-binding protein 1 | Q9ERG0 | YES | NO | YES | YES | YES | YES | P/MQ |
| STRN3 Striatin-3 | Q9ERG2 | NO | YES | NO | NO | NO | NO | P |
| XPO2 Exportin-2 | Q9ERK4 | NO | NO | NO | YES | NO | NO | P |
| GMFG Glia maturation factor gamma | Q9ERL7 | NO | NO | NO | YES | NO | NO | P |
| SCAM2 Secretory carrier-associated membrane protein 2 | Q9ERN0 | NO | NO | NO | YES | NO | NO | P/MQ |
| NDUAD NADH dehydrogenase [ubiquinone] 1 alpha subcomplex subunit 13 | Q9ERS2 | NO | YES | NO | NO | NO | NO | P |

| | | | | | | | | |
|---|---|---|---|---|---|---|---|---|
| RBP2 E3 SUMO-protein ligase RanBP2 | Q9ERU9 | NO | NO | NO | YES | NO | YES | P |
| SHIP1 Phosphatidylinositol 3,4,5-trisphosphate 5-phosphatase 1 | Q9ES52 | NO | NO | NO | YES | YES | YES | P |
| USH1C Harmonin | Q9ES64 | NO | YES | NO | NO | NO | NO | P |
| HRG Histidine-rich glycoprotein | Q9ESB3 | NO | NO | YES | YES | YES | YES | P/MQ |
| LRBA Lipopolysaccharide-responsive and beige-like anchor protein | Q9ESE1 | YES | YES | NO | NO | NO | NO | P |
| CLTRN Collectrin | Q9ESG4 | NO | YES | NO | NO | NO | NO | P |
| AN32B Acidic leucine-rich nuclear phosphoprotein 32 family member B | Q9EST5 | NO | NO | YES | YES | YES | NO | P/MQ |
| BRD4 Bromodomain-containing protein 4 | Q9ESU6 | NO | NO | NO | NO | YES | NO | P |
| DKC1 H/ACA ribonucleoprotein complex subunit DKC1 | Q9ESX5 | NO | NO | NO | YES | NO | YES | P |
| PYGL Glycogen phosphorylase, liver form | Q9ET01 | NO | NO | YES | YES | YES | YES | P/MQ |
| DPP2 Dipeptidyl peptidase 2 | Q9ET22 | NO | YES | NO | NO | NO | NO | P |
| PALLD Palladin | Q9ET54 | YES | NO | YES | YES | YES | YES | P |
| V-type proton ATPase subunit a | Q9JHF5 | YES | YES | YES | YES | YES | YES | MQ |
| CBPB2 Carboxypeptidase B2 | Q9JHH6 | NO | NO | YES | NO | YES | NO | P |
| IVD Isovaleryl-CoA dehydrogenase, mitochondrial | Q9JHI5 | YES | NO | YES | YES | YES | NO | P |
| TMOD3 Tropomodulin-3 | Q9JHJ0 | YES | NO | YES | YES | YES | YES | P/MQ |
| PLEK Pleckstrin | Q9JHK5 | NO | NO | NO | YES | NO | NO | P |
| NHRF2 Na(+)/H(+) exchange regulatory cofactor NHE-RF2 | Q9JHL1 | YES | NO | YES | YES | YES | NO | P/MQ |
| IDE Insulin-degrading enzyme | Q9JHR7 | NO | YES | NO | NO | NO | NO | P |
| DYHC1 Cytoplasmic dynein 1 heavy chain 1 | Q9JHU4 | YES | NO | YES | YES | YES | YES | P/MQ |
| Ino1 Inositol-3-phosphate synthase 1 | Q9JHU9 | YES | NO | NO | NO | NO | NO | P |
| NIT2 Omega-amidase NIT2 | Q9JHW2 | YES | NO | NO | YES | YES | NO | P |
| STK4 Serine/threonine-protein kinase 4 | Q9JI11 | NO | NO | NO | YES | YES | YES | P |
| Placenta-specific gene 8 protein | Q9JI48 | YES | YES | YES | YES | YES | YES | MQ |
| NQO2 Ribosyldihydronicotinamide dehydrogenase [quinone] | Q9JI75 | NO | NO | NO | YES | YES | NO | P/MQ |
| SPHK2 Sphingosine kinase 2 | Q9JIA7 | NO | YES | NO | NO | NO | NO | P |
| ANM1 Protein arginine N-methyltransferase 1 | Q9JIF0 | NO | YES | NO | NO | NO | NO | P |
| COPB Coatomer subunit beta | Q9JIF7 | NO | NO | NO | YES | YES | YES | P/MQ |
| DAZP1 DAZ-associated protein 1 | Q9JII5 | NO | NO | NO | YES | YES | NO | P |
| AK1A1 Aldo-keto reductase family 1 member A1 | Q9JII6 | YES | NO | YES | YES | YES | YES | P/MQ |
| DDX21 Nucleolar RNA helicase 2 | Q9JIK5 | NO | NO | YES | YES | YES | YES | P/MQ |
| NHRF3 Na(+)/H(+) exchange regulatory cofactor NHE-RF3 | Q9JIL4 | YES | NO | NO | NO | YES | NO | P/MQ |
| SMAD9 Mothers against decapentaplegic homolog 9 | Q9JIW5 | NO | YES | NO | NO | NO | NO | P |
| RALB Ras-related protein Ral-B | Q9JIW9 | NO | NO | NO | YES | YES | NO | P |
| ACINU Apoptotic chromatin condensation inducer in the nucleus | Q9JIX8 | YES | NO | YES | YES | YES | YES | P/MQ |
| FLII Protein flightless-1 homolog | Q9JJ28 | NO | NO | NO | YES | YES | YES | P/MQ |

| | | | | | | | | |
|---|---|---|---|---|---|---|---|---|
| RL38 60S ribosomal protein L38 | Q9JJI8 | NO | NO | NO | YES | YES | NO | P |
| SH3L1 SH3 domain-binding glutamic acid-rich-like protein | Q9JJU8 | NO | NO | NO | YES | NO | YES | P/MQ |
| SHLB1 Endophilin-B1 | Q9JK48 | NO | NO | NO | YES | NO | NO | P |
| PRELP Prolargin | Q9JK53 | NO | NO | NO | YES | NO | NO | P |
| MYG1 UPF0160 protein MYG1, mitochondrial | Q9JK81 | NO | YES | NO | NO | NO | NO | P |
| UCHL3 Ubiquitin carboxyl-terminal hydrolase isozyme L3 | Q9JKB1 | NO | NO | YES | YES | YES | YES | P |
| YBOX3 Y-box-binding protein 3 | Q9JKB3 | NO | NO | NO | YES | YES | NO | P/MQ |
| IQGA1 Ras GTPase-activating-like protein IQGAP1 | Q9JKF1 | YES | NO | YES | YES | YES | YES | P/MQ |
| MBNL1 Muscleblind-like protein 1 | Q9JKP5 | NO | YES | NO | NO | NO | NO | P |
| HYOU1 Hypoxia up-regulated protein 1 | Q9JKR6 | NO | NO | YES | YES | YES | YES | P/MQ |
| ADRM1 Proteasomal ubiquitin receptor ADRM1 | Q9JKV1 | NO | NO | NO | YES | YES | YES | P/MQ |
| NUDT5 ADP-sugar pyrophosphatase | Q9JKX6 | NO | NO | NO | YES | NO | NO | P/MQ |
| HIP1R Huntingtin-interacting protein 1-related protein | Q9JKY5 | NO | NO | NO | NO | YES | NO | P |
| FMNL1 Formin-like protein 1 | Q9JL26 | NO | NO | NO | YES | YES | YES | P |
| HMGN5 High mobility group nucleosome-binding domain-containing protein 5 | Q9JL35 | NO | NO | NO | YES | YES | NO | P |
| GLTP Glycolipid transfer protein | Q9JL62 | NO | NO | NO | YES | NO | NO | P |
| MPP6 MAGUK p55 subfamily member 6 | Q9JLB0 | YES | YES | NO | NO | NO | NO | P |
| CUBN Cubilin | Q9JLB4 | YES | NO | YES | NO | YES | NO | P |
| SCLY Selenocysteine lyase | Q9JLI6 | NO | NO | NO | YES | NO | NO | P |
| Squamous cell carcinoma antigen recognized by T-cells 3 | Q9JLI8 | YES | YES | YES | YES | YES | YES | MQ |
| AL9A1 4-trimethylaminobutyraldehyde dehydrogenase | Q9JLJ2 | YES | NO | YES | YES | YES | YES | P/MQ |
| DCLK1 Serine/threonine-protein kinase DCLK1 | Q9JLM8 | NO | NO | NO | YES | NO | NO | P |
| CD2AP CD2-associated protein | Q9JLQ0 | YES | NO | YES | YES | YES | YES | P/MQ |
| GIT2 ARF GTPase-activating protein GIT2 | Q9JLQ2 | NO | NO | NO | YES | NO | NO | P |
| TRXR2 Thioredoxin reductase 2, mitochondrial | Q9JLT4 | NO | YES | NO | NO | NO | NO | P |
| BAG3 BAG family molecular chaperone regulator 3 | Q9JLV1 | NO | NO | YES | YES | YES | YES | P |
| PNKP Bifunctional polynucleotide phosphatase/kinase | Q9JLV6 | NO | NO | YES | YES | YES | YES | P |
| AUHM Methylglutaconyl-CoA hydratase, mitochondrial | Q9JLZ3 | YES | YES | NO | NO | NO | NO | P |
| NT5C 5'(3')-deoxyribonucleotidase, cytosolic type | Q9JM14 | NO | NO | NO | YES | NO | NO | P |
| ARPC3 Actin-related protein 2/3 complex subunit 3 | Q9JM76 | NO | NO | YES | YES | YES | YES | P |
| BORG4 Cdc42 effector protein 4 | Q9JM96 | NO | NO | NO | YES | NO | NO | P |
| UBP14 Ubiquitin carboxyl-terminal hydrolase 14 | Q9JMA1 | NO | NO | NO | YES | NO | NO | P/MQ |
| STA10 START domain-containing protein 10 | Q9JMD3 | NO | YES | NO | NO | NO | NO | P |
| EDF1 Endothelial differentiation-related factor 1 | Q9JMG1 | NO | NO | NO | YES | NO | NO | P |
| TRXR1 Thioredoxin reductase 1, cytoplasmic | Q9JMH6 | YES | NO | YES | YES | YES | YES | P/MQ |
| MY18A Unconventional myosin-XVIIIa | Q9JMH9 | YES | NO | YES | YES | YES | YES | P/MQ |
| HAOX2 Hydroxyacid oxidase 2 | Q9NYQ2 | YES | NO | YES | NO | YES | NO | P |

| | | | | | | | | |
|---|---|---|---|---|---|---|---|---|
| GLRX1 Glutaredoxin-1 | Q9QUH0 | NO | NO | YES | YES | YES | YES | P |
| RHOA Transforming protein RhoA | Q9QUI0 | YES | NO | YES | YES | YES | YES | P/MQ |
| PSA6 Proteasome subunit alpha type-6 | Q9QUM9 | NO | NO | NO | YES | NO | YES | P |
| PPCE Prolyl endopeptidase | Q9QUR6 | NO | NO | NO | YES | YES | YES | P |
| PIN1 Peptidyl-prolyl cis-trans isomerase NIMA-interacting 1 | Q9QUR7 | NO | NO | NO | YES | NO | YES | P |
| NAGAB Alpha-N-acetylgalactosaminidase | Q9QWR8 | NO | YES | NO | NO | NO | NO | P |
| CYH1 Cytohesin-1 | Q9QX11 | NO | YES | NO | NO | NO | NO | P |
| SON Protein SON | Q9QX47 | NO | NO | YES | YES | YES | NO | P/MQ |
| FBP1 Fructose-1,6-bisphosphatase 1 | Q9QXD6 | YES | NO | NO | NO | YES | NO | P/MQ |
| ACSS2 Acetyl-coenzyme A synthetase, cytoplasmic | Q9QXG4 | NO | YES | NO | NO | NO | NO | P |
| SHRM3 Protein Shroom3 | Q9QXN0 | NO | YES | NO | NO | NO | NO | P |
| MIOX Inositol oxygenase | Q9QXN5 | NO | YES | NO | NO | NO | NO | P |
| PLEC Plectin | Q9QXS1 | YES | NO | YES | YES | YES | YES | P/MQ |
| DREB Drebrin | Q9QXS6 | NO | NO | NO | YES | YES | YES | P/MQ |
| CNPY2 Protein canopy homolog 2 | Q9QXT0 | YES | NO | YES | YES | YES | YES | P/MQ |
| LAT2 Large neutral amino acids transporter small subunit 2 | Q9QXW9 | NO | YES | NO | NO | NO | NO | P |
| CMC2 Calcium-binding mitochondrial carrier protein Aralar2 | Q9QXX4 | YES | YES | NO | NO | NO | NO | P |
| EHD3 EH domain-containing protein 3 | Q9QXY6 | NO | NO | NO | NO | YES | NO | P |
| MACF1 Microtubule-actin cross-linking factor 1 | Q9QXZ0 | YES | NO | NO | YES | YES | YES | P/MQ |
| MYO9B Unconventional myosin-IXb | Q9QY06 | NO | NO | NO | YES | YES | YES | P |
| ZBP1 Z-DNA-binding protein 1 | Q9QY24 | NO | NO | NO | NO | YES | NO | P/MQ |
| VAPB Vesicle-associated membrane protein-associated protein B | Q9QY76 | YES | NO | YES | YES | YES | YES | P/MQ |
| CLIC4 Chloride intracellular channel protein 4 | Q9QYB1 | YES | NO | NO | YES | YES | YES | P/MQ |
| ADDG Gamma-adducin | Q9QYB5 | YES | NO | NO | NO | YES | NO | P |
| ADDA Alpha-adducin | Q9QYC0 | YES | NO | YES | YES | YES | YES | P/MQ |
| Golgin subfamily A member 5 | Q9QYE6 | YES | YES | YES | YES | NO | YES | MQ |
| NDRG2 Protein NDRG2 | Q9QYG0 | NO | YES | NO | NO | NO | NO | P |
| DNJA2 DnaJ homolog subfamily A member 2 | Q9QYJ0 | NO | YES | NO | NO | NO | NO | P |
| ACOT2 Acyl-coenzyme A thioesterase 2, mitochondrial | Q9QYR9 | NO | NO | NO | YES | YES | YES | P |
| QKI Protein quaking | Q9QYS9 | NO | NO | NO | YES | NO | NO | P/MQ |
| PPM1D Protein phosphatase 1D | Q9QZ67 | NO | YES | NO | NO | NO | NO | P |
| IIGP1 Interferon-inducible GTPase 1 | Q9QZ85 | NO | NO | YES | YES | YES | YES | P |
| VPS29 Vacuolar protein sorting-associated protein 29 | Q9QZ88 | NO | YES | NO | NO | NO | NO | P |
| EIF3I Eukaryotic translation initiation factor 3 subunit I | Q9QZD9 | NO | NO | NO | YES | YES | YES | P/MQ |
| COPG1 Coatomer subunit gamma-1 | Q9QZE5 | YES | NO | YES | YES | YES | YES | P/MQ |
| BCAR3 Breast cancer anti-estrogen resistance protein 3 | Q9QZK2 | NO | YES | NO | NO | NO | NO | P |
| DOK3 Docking protein 3 | Q9QZK7 | NO | NO | NO | YES | YES | NO | P |

| | | | | | | | | |
|---|---|---|---|---|---|---|---|---|
| RIPK3 Receptor-interacting serine/threonine-protein kinase 3 | Q9QZL0 | NO | NO | NO | YES | YES | NO | P |
| UBQL2 Ubiquilin-2 | Q9QZM0 | NO | NO | NO | YES | NO | NO | P |
| AFAD Afadin | Q9QZQ1 | NO | NO | NO | YES | YES | NO | P |
| H2AY Core histone macro-H2A.1 | Q9QZQ8 | YES | NO | YES | YES | YES | YES | P/MQ |
| GLYG Glycogenin-1 | Q9R062 | NO | NO | NO | YES | YES | NO | P |
| BCAM Basal cell adhesion molecule | Q9R069 | YES | YES | NO | NO | NO | NO | P |
| GALK1 Galactokinase | Q9R0N0 | NO | NO | NO | YES | NO | NO | P |
| SYT7 Synaptotagmin-7 | Q9R0N7 | NO | YES | NO | NO | NO | NO | P |
| ESTD S-formylglutathione hydrolase | Q9R0P3 | YES | NO | YES | YES | YES | YES | P/MQ |
| SMAP Small acidic protein | Q9R0P4 | NO | NO | NO | YES | YES | YES | P/MQ |
| DEST Destrin | Q9R0P5 | YES | NO | YES | YES | YES | YES | P/MQ |
| ARC1A Actin-related protein 2/3 complex subunit 1A | Q9R0Q6 | NO | YES | NO | NO | NO | NO | P |
| TEBP Prostaglandin E synthase 3 | Q9R0Q7 | NO | YES | NO | NO | NO | NO | P |
| SRS10 Serine/arginine-rich splicing factor 10 | Q9R0U0 | NO | NO | NO | YES | NO | NO | P |
| ACOT9 Acyl-coenzyme A thioesterase 9, mitochondrial | Q9R0X4 | NO | YES | NO | NO | NO | NO | P |
| KAD1 Adenylate kinase isoenzyme 1 | Q9R0Y5 | YES | YES | NO | NO | NO | NO | P |
| GUAD Guanine deaminase | Q9R111 | NO | NO | YES | YES | YES | YES | P/MQ |
| SQOR Sulfide:quinone oxidoreductase, mitochondrial | Q9R112 | YES | NO | YES | YES | YES | YES | P/MQ |
| MTA2 Metastasis-associated protein MTA2 | Q9R190 | NO | NO | NO | YES | NO | NO | P |
| PSA4 Proteasome subunit alpha type-4 | Q9R1P0 | NO | NO | NO | NO | YES | YES | P |
| PSB2 Proteasome subunit beta type-2 | Q9R1P3 | NO | NO | NO | YES | NO | NO | P |
| PSA1 Proteasome subunit alpha type-1 | Q9R1P4 | YES | NO | YES | YES | YES | YES | P/MQ |
| SAE1 SUMO-activating enzyme subunit 1 | Q9R1T2 | NO | NO | YES | YES | YES | YES | P/MQ |
| SEPT6 Septin-6 | Q9R1T4 | NO | NO | NO | YES | NO | NO | P |
| VINEX Vinexin | Q9R1Z8 | NO | YES | NO | NO | NO | NO | P/MQ |
| TPSN Tapasin | Q9R233 | NO | NO | YES | YES | YES | YES | P |
| PEPL Periplakin | Q9R269 | NO | YES | NO | NO | NO | NO | P |
| MYO1C Unconventional myosin-Ic | Q9WTI7 | YES | NO | YES | YES | YES | YES | P/MQ |
| NFKB2 Nuclear factor NF-kappa-B p100 subunit | Q9WTK5 | NO | NO | NO | YES | NO | YES | P |
| RUVB2 RuvB-like 2 | Q9WTM5 | NO | NO | YES | YES | YES | YES | P/MQ |
| KAD2 Adenylate kinase 2, mitochondrial | Q9WTP6 | YES | NO | YES | YES | YES | YES | P/MQ |
| KAD3 GTP:AMP phosphotransferase AK3, mitochondrial | Q9WTP7 | YES | NO | YES | YES | YES | NO | P/MQ |
| AKA12 A-kinase anchor protein 12 | Q9WTQ5 | YES | NO | YES | YES | YES | YES | P/MQ |
| CAD13 Cadherin-13 | Q9WTR5 | NO | YES | NO | NO | NO | NO | P |
| SKP1 S-phase kinase-associated protein 1 | Q9WTX5 | NO | NO | YES | YES | YES | NO | P/MQ |
| PDC6I Programmed cell death 6-interacting protein | Q9WU78 | YES | NO | YES | YES | YES | YES | P/MQ |
| PROD Proline dehydrogenase 1, mitochondrial | Q9WU79 | YES | YES | NO | NO | NO | NO | P |
| CCS Copper chaperone for superoxide dismutase | Q9WU84 | NO | YES | NO | NO | NO | NO | P |
| SYFB Phenylalanine--tRNA ligase beta subunit | Q9WUA2 | YES | NO | YES | YES | YES | YES | P |
| PYGM Glycogen phosphorylase, muscle form | Q9WUB3 | NO | YES | NO | NO | NO | NO | P |

| | | | | | | | | |
|---|---|---|---|---|---|---|---|---|
| CLCKB Chloride channel protein ClC-Kb | Q9WUB6 | NO | YES | NO | NO | NO | NO | P |
| IF4H Eukaryotic translation initiation factor 4H | Q9WUK2 | NO | NO | YES | YES | YES | YES | P/MQ |
| ARL3 ADP-ribosylation factor-like protein 3 | Q9WUL7 | NO | YES | NO | NO | NO | NO | P |
| COR1B Coronin-1B | Q9WUM3 | NO | NO | NO | YES | YES | YES | P |
| COR1C Coronin-1C | Q9WUM4 | NO | NO | NO | YES | NO | NO | P |
| UCHL5 Ubiquitin carboxyl-terminal hydrolase isozyme L5 | Q9WUP7 | NO | YES | NO | NO | NO | NO | P |
| PREB Prolactin regulatory element-binding protein | Q9WUQ2 | NO | NO | NO | NO | YES | NO | P |
| ECI2 Enoyl-CoA delta isomerase 2, mitochondrial | Q9WUR2 | YES | NO | YES | YES | YES | NO | P |
| KAD4 Adenylate kinase 4, mitochondrial | Q9WUR9 | YES | NO | NO | NO | NO | NO | P |
| CATZ Cathepsin Z | Q9WUU7 | NO | NO | NO | YES | YES | YES | P/MQ |
| RBMX RNA-binding motif protein, X chromosome | Q9WV02 | NO | NO | YES | YES | YES | YES | P/MQ |
| ARC1B Actin-related protein 2/3 complex subunit 1B | Q9WV32 | YES | NO | YES | YES | YES | YES | P/MQ |
| VAPA Vesicle-associated membrane protein-associated protein A | Q9WV55 | NO | NO | NO | YES | YES | YES | P |
| DEMA Dematin | Q9WV69 | NO | YES | NO | NO | NO | NO | P |
| SNX1 Sorting nexin-1 | Q9WV80 | YES | NO | YES | YES | YES | YES | P/MQ |
| E41L3 Band 4.1-like protein 3 | Q9WV92 | YES | NO | YES | NO | YES | NO | P/MQ |
| TIM8A Mitochondrial import inner membrane translocase subunit Tim8 A | Q9WVA2 | NO | YES | NO | NO | NO | NO | P |
| BUB3 Mitotic checkpoint protein BUB3 | Q9WVA3 | NO | NO | NO | YES | YES | YES | P |
| TAGL2 Transgelin-2 | Q9WVA4 | YES | NO | YES | YES | YES | YES | P/MQ |
| RBPMS RNA-binding protein with multiple splicing | Q9WVB0 | YES | NO | YES | YES | YES | YES | P |
| PACN2 Protein kinase C and casein kinase substrate in neurons protein 2 | Q9WVE8 | YES | NO | YES | YES | YES | YES | P/MQ |
| EHD1 EH domain-containing protein 1 | Q9WVK4 | NO | NO | YES | YES | YES | YES | P/MQ |
| MAAI Maleylacetoacetate isomerase | Q9WVL0 | YES | YES | NO | NO | NO | NO | P |
| S12A7 Solute carrier family 12 member 7 | Q9WVL3 | NO | YES | NO | NO | NO | NO | P |
| AADAT Kynurenine/alpha-aminoadipate aminotransferase, mitochondrial | Q9WVM8 | NO | NO | NO | YES | YES | NO | P |
| GBP2 Guanylate-binding protein 2 | Q9Z0E6 | NO | NO | YES | YES | YES | YES | P/MQ |
| SYUG Gamma-synuclein | Q9Z0F7 | YES | NO | NO | NO | YES | NO | P/MQ |
| CELF2 CUGBP Elav-like family member 2 | Q9Z0H4 | NO | NO | NO | YES | NO | NO | P |
| CLIP2 CAP-Gly domain-containing linker protein 2 | Q9Z0H8 | NO | NO | NO | YES | NO | NO | P |
| VNN1 Pantetheinase | Q9Z0K8 | YES | YES | NO | NO | NO | NO | P |
| IF2G Eukaryotic translation initiation factor 2 subunit 3, X-linked | Q9Z0N1 | NO | NO | YES | YES | YES | YES | P/MQ |
| PALM Paralemmin-1 | Q9Z0P4 | YES | NO | NO | YES | YES | YES | P/MQ |
| TWF2 Twinfilin-2 | Q9Z0P5 | NO | NO | NO | YES | YES | YES | P |
| ITSN1 Intersectin-1 | Q9Z0R4 | NO | NO | NO | YES | NO | NO | P |
| ITSN2 Intersectin-2 | Q9Z0R6 | NO | NO | NO | YES | NO | NO | P |
| BPNT1 3'(2'),5'-bisphosphate nucleotidase 1 | Q9Z0S1 | NO | YES | NO | NO | NO | NO | P |
| ZO2 Tight junction protein ZO-2 | Q9Z0U1 | YES | NO | YES | YES | YES | YES | P/MQ |

| | | | | | | | | |
|---|---|---|---|---|---|---|---|---|
| TI17B Mitochondrial import inner membrane translocase subunit Tim17-B | Q9Z0V7 | NO | YES | NO | NO | NO | NO | P |
| AIFM1 Apoptosis-inducing factor 1, mitochondrial | Q9Z0X1 | YES | NO | YES | YES | YES | NO | P/MQ |
| STAU1 Double-stranded RNA-binding protein Staufen homolog 1 | Q9Z108 | NO | NO | NO | NO | YES | NO | P |
| HNRDL Heterogeneous nuclear ribonucleoprotein D-like | Q9Z130 | NO | NO | NO | YES | NO | NO | P |
| SUMO3 Small ubiquitin-related modifier 3 | Q9Z172 | NO | YES | NO | NO | NO | NO | P |
| PADI4 Protein-arginine deiminase type-4 | Q9Z183 | NO | NO | NO | YES | YES | YES | P |
| EIF3G Eukaryotic translation initiation factor 3 subunit G | Q9Z1D1 | YES | NO | YES | YES | YES | YES | P/MQ |
| GYS1 Glycogen [starch] synthase, muscle | Q9Z1E4 | NO | NO | NO | YES | NO | NO | P |
| SAE2 SUMO-activating enzyme subunit 2 | Q9Z1F9 | NO | NO | NO | YES | NO | NO | P/MQ |
| VATC1 V-type proton ATPase subunit C 1 | Q9Z1G3 | NO | NO | NO | YES | NO | NO | P/MQ |
| NFS1 Cysteine desulfurase, mitochondrial | Q9Z1J3 | NO | YES | NO | NO | NO | NO | P |
| DX39B Spliceosome RNA helicase Ddx39b | Q9Z1N5 | NO | NO | YES | NO | YES | YES | P |
| NDUA7 NADH dehydrogenase [ubiquinone] 1 alpha subcomplex subunit 7 | Q9Z1P6 | YES | YES | NO | NO | NO | NO | P |
| CLIC1 Chloride intracellular channel protein 1 | Q9Z1Q5 | YES | NO | YES | YES | YES | YES | P/MQ |
| SYVC Valine--tRNA ligase | Q9Z1Q9 | NO | NO | YES | YES | YES | YES | P/MQ |
| BAG6 Large proline-rich protein BAG6 | Q9Z1R2 | NO | NO | NO | YES | NO | NO | P |
| ILF3 Interleukin enhancer-binding factor 3 | Q9Z1X4 | NO | NO | NO | YES | NO | YES | P |
| TRIP6 Thyroid receptor-interacting protein 6 | Q9Z1Y4 | NO | YES | NO | NO | NO | NO | P |
| USO1 General vesicular transport factor p115 | Q9Z1Z0 | NO | NO | NO | YES | YES | YES | P/MQ |
| STRAP Serine-threonine kinase receptor-associated protein | Q9Z1Z2 | YES | NO | NO | NO | YES | NO | P |
| HNRPC Heterogeneous nuclear ribonucleoproteins C1/C2 | Q9Z204 | YES | NO | YES | YES | YES | YES | P/MQ |
| MECP2 Methyl-CpG-binding protein 2 | Q9Z2D6 | YES | NO | YES | YES | YES | YES | P/MQ |
| MBD2 Methyl-CpG-binding domain protein 2 | Q9Z2E1 | NO | NO | NO | YES | NO | NO | P |
| E41L1 Band 4.1-like protein 1 | Q9Z2H5 | NO | YES | NO | NO | NO | NO | P |
| GIPC2 PDZ domain-containing protein GIPC2 | Q9Z2H7 | YES | YES | NO | NO | NO | NO | P |
| LETM1 Mitochondrial proton/calcium exchanger protein | Q9Z2I0 | YES | NO | NO | YES | NO | NO | P |
| S23A1 Solute carrier family 23 member 1 | Q9Z2J0 | NO | YES | NO | NO | NO | NO | P |
| PMM2 Phosphomannomutase 2 | Q9Z2M7 | NO | NO | YES | YES | NO | YES | P |
| ACL6A Actin-like protein 6A | Q9Z2N8 | NO | NO | NO | YES | NO | NO | P |
| K0P1 Lysine-rich nucleolar protein 1 | Q9Z2Q2 | NO | YES | NO | NO | NO | NO | P |
| SEPT5 Septin-5 | Q9Z2Q6 | NO | NO | NO | YES | NO | NO | P |
| U119A Protein unc-119 homolog A | Q9Z2R6 | NO | NO | NO | YES | NO | NO | P |
| PSA7 Proteasome subunit alpha type-7 | Q9Z2U0 | YES | NO | YES | YES | YES | YES | P/MQ |
| PSA5 Proteasome subunit alpha type-5 | Q9Z2U1 | NO | NO | NO | YES | NO | YES | P/MQ |

| | | | | | | | | |
|---|---|---|---|---|---|---|---|---|
| PCK1 Phosphoenolpyruvate carboxykinase, cytosolic [GTP] | Q9Z2V4 | YES | NO | NO | NO | YES | NO | P |
| DNPEP Aspartyl aminopeptidase | Q9Z2W0 | NO | YES | NO | NO | NO | NO | P |
| HNRPF Heterogeneous nuclear ribonucleoprotein F | Q9Z2X1 | NO | NO | YES | YES | YES | YES | P/MQ |
| PLPHP Pyridoxal phosphate homeostasis protein | Q9Z2Y8 | NO | NO | YES | NO | YES | YES | P |
| SNUT1 U4/U6.U5 tri-snRNP-associated protein 1 | Q9Z315 | YES | NO | YES | YES | YES | YES | P/MQ |
| ITPR2 Inositol 1,4,5-trisphosphate receptor type 2 | Q9Z329 | NO | YES | NO | NO | NO | NO | P |
| 2-oxoglutarate dehydrogenase, mitochondrial | Z4YJV4 | YES | YES | YES | YES | YES | YES | MQ |

Table S1: List of host-derived proteins found by presence at time point and biological region, including platform used for identification (P = Protalizer, MQ = MaxQuant).

| Protein Name | Protein Id | Gene product | Function | 4 dpi SAC | 10 dpi SAC | 4 dpi Interface | 10 dpi Interface | Platform |
|---|---|---|---|---|---|---|---|---|
| 50S ribosomal protein L30 | Q2FEQ7 | RpmD | Ribosome | YES | YES | NO | YES | P/MQ |
| 50S ribosomal protein L17 | Q2FER6 | RplQ | Ribosome | YES | YES | NO | YES | P/MQ |
| Threonine--tRNA ligase | Q2FG54 | ThrS | Translation | NO | YES | YES | YES | P/MQ |
| Elongation factor Ts | Q2FHI1 | Tsf | Translation | YES | YES | NO | YES | P/MQ |
| Elongation factor Tu | Q2FJ92 | Tuf | Translation | YES | YES | YES | YES | P/MQ |
| Elongation factor G | Q2FJ93 | FusA | Ribosome | NO | YES | YES | YES | P |
| 50S ribosomal protein L7/L12 | Q2FJA0 | RplL | Ribosome | YES | YES | NO | YES | P |
| Uncharacterized leukocidin-like protein 2 | Q2FFA2 | LukA | Virulence | NO | YES | NO | YES | P/MQ |
| Extracellular matrix-binding protein ebh | Q2FH04 | Ebh | Virulence | YES | YES | YES | YES | P |
| Extracellular adherence protein | A6QIG2 | Eap | Virulence | YES | YES | YES | YES | P |
| FPRL1 inhibitory protein | Q2FHS7 | FLIPr | Virulence/ Immune Evasion | NO | YES | NO | YES | P/MQ |
| Extracellular matrix protein-binding protein emp | Q2FIK4 | Emp | Virulence | YES | YES | YES | YES | P/MQ |
| Iron-regulated surface determinant protein A | Q2FHV1 | IsdA | Metal homeostasis | NO | YES | NO | YES | P/MQ |
| Iron-regulated surface determinant protein B | Q2FHV2 | IsdB | Metal homeostasis | NO | NO | NO | YES | P |
| L-lactate dehydrogenase 2 | Q2FDQ7 | Ldh2 | Anaerobic metabolism | YES | YES | YES | YES | P/MQ |
| Enolase | Q2FIL7 | Eno | Anaerobic metabolism | YES | YES | NO | YES | P/MQ |
| Alcohol dehydrogenase | Q2FJ31 | Adh | Anaerobic metabolism | YES | YES | YES | YES | P/MQ |
| Formate acetyltransferase | Q2FK44 | PflB | Anaerobic metabolism | YES | YES | YES | YES | P/MQ |
| ATP-dependent Clp protease ATP-binding subunit ClpL | Q2FDV8 | ClpL | Stress | NO | YES | NO | YES | P |
| Putative universal stress protein SAUSA300_1656 | Q2FG28 | USP2 | Stress | YES | YES | YES | YES | P/MQ |
| Alanine dehydrogenase 1 | Q2FH00 | Ald1 | Cell Wall/Stress | NO | YES | NO | YES | P/MQ |
| Alkyl hydroperoxide reductase C | Q2FJN4 | AhpC | Stress | YES | YES | YES | YES | P/MQ |
| Trigger factor | Q2FG61 | Tig | Protein folding | NO | YES | NO | YES | P/MQ |

| | | | | | | | | |
|---|---|---|---|---|---|---|---|---|
| Chaperone protein DnaK | Q2FGE3 | DnaK | Protein folding | YES | YES | YES | YES | P/MQ |
| UPF0337 protein SAUSA300_0816 | Q2FIG2 | SAUSA300_0816 | Unknown function | NO | NO | NO | YES | P/MQ |
| GTP-sensing transcriptional pleiotropic repressor CodY | Q2FHI3 | CodY | Regulation/ Nutrition | YES | YES | NO | YES | P/MQ |
| Fructose-bisphosphate aldolase class 1 | Q2FDQ4 | Fda | Glycolysis | YES | YES | YES | YES | P/MQ |
| ATP synthase subunit alpha | Q2FF22 | AtpA | Energy | YES | YES | YES | YES | P |
| ATP synthase subunit beta | Q2FF24 | AtpD | Energy | YES | YES | YES | YES | P |
| S-adenosylmethionine synthase | Q2FFV6 | MetK | Amino acid biosynthesis | YES | YES | YES | YES | P |
| Pyruvate kinase | Q2FG40 | Pyk | Glycolysis | NO | YES | NO | YES | P |
| Glucose-6-phosphate isomerase | Q2FIB3 | Pgi | Glycolysis | NO | YES | NO | YES | P/MQ |

Table S2: List of bacterial proteins detected and their localizations based on stringent search criteria from Protalizer and MaxQuant. No bacterial proteins were detected in the cortex.

| Bacteria Name | | Gene | Category | 4 dpi SAC | 10 dpi SAC | 4 dpi Interface | 10 dpi Interface |
|---|---|---|---|---|---|---|---|
| 30S ribosomal protein S10 | Q2FEN8 | RpsJ | Ribosome | NO | YES | NO | YES |
| 50S ribosomal protein L2 | Q2FEP2 | RplB | Ribosome | YES | YES | NO | YES |
| 30S ribosomal protein S8 | Q2FEQ3 | RpsH | Ribosome | NO | YES | NO | YES |
| 50S ribosomal protein L6 | Q2FEQ4 | RplF | Ribosome | NO | NO | NO | YES |
| 50S ribosomal protein L18 | Q2FEQ5 | RplR | Ribosome | NO | NO | NO | YES |
| 30S ribosomal protein S5 | Q2FEQ6 | RpsE | Ribosome | NO | NO | NO | YES |
| 50S ribosomal protein L15 | Q2FEQ8 | RplO | Ribosome | NO | YES | NO | YES |
| 50S ribosomal protein L21 | Q2FG80 | RplU | Ribosome | NO | YES | NO | YES |
| 30S ribosomal protein S20 | Q2FGD8 | RpsT | Ribosome | NO | YES | NO | YES |
| Ribosome-recycling factor | Q2FHH9 | Frr | Ribosome | NO | YES | NO | YES |
| 50S ribosomal protein L25 | Q2FJE0 | RplY | Ribosome | NO | YES | NO | YES |
| Uncharacterized leukocidin-like protein 1 | Q2FFA3 | LukB | Virulence | NO | YES | NO | YES |
| Chemotaxis inhibitory protein | Q2FFF7 | Chp | Virulence/Immune Evasion | NO | YES | NO | YES |
| Serine-aspartate repeat-containing protein E | Q2FJ77 | SdrE | Virulence/Immune Evasion | NO | YES | NO | YES |
| Putative heme-dependent peroxidase SAUSA300_0569 | Q2FJ56 | ChdC | Metal homeostasis | NO | YES | NO | YES |
| Sulfur carrier protein FdhD | Q2FEL2 | FdhD | Sulfur Carrier Protein | NO | NO | NO | YES |
| Acetate kinase | Q2FG27 | AckA | Anaerobic metabolism | NO | YES | NO | NO |
| L-lactate dehydrogenase 1 | Q2FK29 | Ldh1 | Anaerobic metabolism | YES | YES | NO | YES |
| UPF0342 protein SAUSA300_1795 | Q2FFQ0 | SAUSA300_1795 | Unknown function | NO | YES | NO | YES |
| 60 kDa chaperonin | Q2FF95 | GroL | Protein folding | NO | YES | NO | YES |
| Thioredoxin | Q2FHT6 | TrxA | Redox reactions | NO | YES | NO | YES |

| | | | | | | | |
|---|---|---|---|---|---|---|---|
| HTH-type transcriptional regulator ArcR | Q2FDM8 | ArcR | Regulation/Anaerobic Conditions | NO | NO | YES | NO |
| Transcription elongation factor GreA | Q2FGB6 | GreA | Transcription Elongation Factor | NO | YES | NO | YES |
| Nuclease SbcCD subunit C | Q2FH88 | SbcC | DNA Replication | YES | NO | YES | NO |
| Pyrimidine-nucleoside phosphorylase | Q2FEZ3 | Pdp | Phosphorolysis of the pyrimidine nucleosides | YES | YES | NO | YES |
| ATP synthase subunit b | Q2FF20 | AtpF | Energy | NO | YES | NO | YES |
| Probable manganese-dependent inorganic pyrophosphatase | Q2FFH6 | PpaC | Unknown function | YES | NO | YES | NO |
| Phosphoenolpyruvate carboxykinase (ATP) | Q2FFV5 | PckA | Gluconeogenesis | NO | YES | NO | YES |
| L-threonine dehydratase catabolic TdcB | Q2FH01 | TdcB | L-threonine degradation via propanoate pathway | NO | YES | NO | YES |
| Bifunctional protein FolD | Q2FI15 | FoldD | Folate biosynthesis | NO | YES | NO | YES |

Table S3: List of bacterial proteins detected and their localizations with lower identification confidence.
